# Functional orthogonality of WRN inhibitor resistance enables alternating therapy and mutation-tolerant inhibitor design in MSI-H cancers

**DOI:** 10.64898/2026.06.09.731233

**Authors:** Guizhen Zhou, Dan Teng, Dan Wang, Miao Pang, Zisheng Fan, Yuyan Tao, Chen Shi, Xinyu Yang, Manlin Huang, Panpan Shao, Yun Liu, Rongrong Cui, Zhiyin Xie, Yuanyang Zhou, Zhiming Ge, Xinyi Ma, Jiahang Xu, Shuqing Chu, Yang Chen, Zhehuan Fan, Qibang Sui, Ruirui Yang, Chong Qin, Xutong Li, Kaixian Chen, Xiaomin Luo, Mingliang Wang, Sulin Zhang, Mingyue Zheng

## Abstract

Werner syndrome helicase (WRN) is a synthetic-lethal vulnerability in microsatellite instability–high (MSI-H)/mismatch repair–deficient (dMMR) cancers^1–5^, and non-covalent and covalent WRN inhibitors are now entering clinical development^6–9^. A central unresolved question is whether resistance to one WRN inhibitor class inevitably compromises the target, or instead creates actionable vulnerabilities to an alternate modality. Here we generated ten stable acquired-resistance models across three MSI-H cell lines using the non-covalent inhibitor HRO761 and the covalent inhibitor VVD-214. Whole-exome sequencing, biochemical reconstitution and isogenic knock-in models identified recurrent on-target WRN missense mutations as dominant resistance drivers, but with sharply modality-specific spectra: HRO761 resistance clustered at G729/F730/I852, whereas VVD-214 resistance concentrated at E846. These mutations impaired inhibitor engagement and abolished the canonical WRN inhibitor (WRNi)-induced pharmacodynamic cascade, including WRN reduction, DNA damage response (DDR) activation and G_2_/M arrest. Crucially, most resistance mutations retained biochemical, cellular and in vivo sensitivity to the alternate inhibitor modality, revealing a functional orthogonality that enabled a 7-day cyclic alternating regimen to delay tumour regrowth in xenografts. We further identified F730L as an engineered cross-resistant bottleneck model and used structure-guided, artificial intelligence (AI)-enabled optimization to generate GBA-007, a proof-of-concept mutation-tolerant WRN inhibitor candidate with promising activity against F730L in vitro and in vivo. Thus, clinically relevant WRN inhibitor classes impose distinct on-target resistance trajectories that can be exploited through schedule design, while cross-resistant bottlenecks can be addressed by rapid mutation-aware inhibitor engineering.

## Introduction

Microsatellite instability–high (MSI-H) or mismatch repair–deficient (dMMR) tumours represent a clinically important molecular subtype across multiple cancer types, including colorectal and endometrial cancers^10–13^. Defective DNA mismatch repair drives the progressive accumulation of insertion/deletion mutations and repeat expansions within microsatellite sequences, thereby increasing genomic heterogeneity and promoting replication-associated stress^5,10–13^. Immune checkpoint inhibitors (ICIs) have transformed the treatment landscape of MSI-H/dMMR disease^14–17^, yet a substantial fraction of patients exhibit primary resistance or ultimately experience progression after an initial response, underscoring the need for additional precision therapies that exploit MSI-H–specific vulnerabilities^18,19^.

Large-scale functional genomics studies have consistently identified Werner syndrome helicase (WRN) as a highly selective synthetic lethal dependency in MSI-H/dMMR contexts^1–5^. Genetic depletion of WRN induces profound DNA damage, checkpoint activation and growth arrest preferentially in MSI-H models, while exerting comparatively limited effects in microsatellite-stable (MSS) backgrounds^2–5^. Mechanistically, mismatch repair (MMR) deficiency–associated repeat expansions—particularly TA dinucleotide repeat expansions—are thought to form abnormal DNA secondary structures that create a non-redundant requirement for WRN helicase activity to maintain replication fork integrity and genome stability^5^. Together, these data establish WRN as a therapeutically actionable target with a clear biological rationale in MSI-H/dMMR tumours.

Building on this genetic foundation, the development of WRN-targeted therapeutics has advanced rapidly. Multiple small-molecule WRN inhibitors (WRNi) have been reported with selective activity in MSI-H/dMMR cellular and xenograft models, often accompanied by induction of DNA damage response (DDR) signalling and marked tumour growth inhibition^6,8,20–23^. Importantly, two clinically relevant WRNi modalities have entered early clinical development^6–9^. HRO761 is a non-covalent inhibitor with strong preclinical activity in MSI-H models and encouraging early clinical signals^6,7^. VVD-214/RO7589831 (also known as VVD-133214) represents a covalent inhibitor class with early clinical safety and preliminary antitumour activity reported^8,9^. The concurrent clinical emergence of mechanistically distinct WRNi classes creates a near-term opportunity: WRN inhibition may become an actionable precision therapy for MSI-H/dMMR disease, and multiple agents may be available for rational sequencing or combination^24–27^.

However, experience across targeted oncology strongly suggests that acquired resistance will be a central determinant of long-term benefit^28–30^. This concern is particularly salient in MSI-H/dMMR tumours, where elevated mutational burden and clonal diversity may expand accessible evolutionary trajectories under drug selection^10–13,31^. In other targeted therapy settings, resistance mechanisms frequently converge on on-target alterations that diminish inhibitor engagement, but can also involve pathway re-wiring, phenotypic adaptation, drug efflux, or microenvironmental effects^28–30^. Understanding the molecular basis of WRNi resistance is therefore essential not only for anticipating clinical failure modes, but also for informing trial design, biomarker development, and next-generation inhibitor strategies.

Recent preclinical studies have begun to illuminate features of WRNi resistance. Evidence suggests that on-target alterations in WRN can emerge under pharmacologic selection, whereas genetic screens have not identified a universal bypass mechanism capable of broadly circumventing WRN dependency^24–27^. In parallel, structural and biochemical studies indicate that distinct WRNi classes can modulate WRN through different conformational trapping mechanisms^6,8,27^. This raises a clinically consequential possibility: different WRNi classes may impose distinct selective pressures, leading to drug-specific resistance mutation patterns; some resistance events may confer cross-resistance, whereas others may preserve sensitivity to an alternative inhibitor class^24–27^. Despite these advances, several fundamental questions remain unanswered. First, the molecular basis of acquired resistance has not been systematically defined across clinically relevant WRNi modalities in a way that links genomic events to biochemical mechanism and cellular phenotype. Second, even if different inhibitor classes select distinct resistance events, it remains unclear whether such differences translate into functionally actionable cross-sensitivity that can be exploited therapeutically—particularly in long-term in vivo settings where clonal dynamics and pharmacologic constraints shape resistance evolution. Third, even if cross-sensitivity exists for many resistance events, it is unknown whether resistance evolution nonetheless generates cross-resistant bottlenecks that compromise both inhibitor classes, and how such bottlenecks might be efficiently overcome in a clinically relevant timeframe.

These considerations motivate three key questions. First, what is the molecular basis of acquired resistance to WRN inhibition, and does resistance evolve through distinct molecular events under non-covalent versus covalent inhibitory mechanisms? Second, do orthogonal functional resistance profiles exist—such that resistance driven by one WRNi class remains tractable to another—and if so, how can this be translated into a rational treatment strategy to extend tumour control, including the potential for cyclic alternating therapy? Third, does resistance evolution generate cross-resistant bottlenecks that limit both inhibitor classes, and if so, can rapid, structure-informed and artificial intelligence (AI)-enabled molecular design provide an efficient route to next-generation WRN inhibitors capable of overcoming such bottlenecks?

Here, we address these questions using two clinically relevant WRN inhibitors with distinct mechanisms of action, HRO761 and VVD-214^6,8^. We establish acquired resistance models in multiple MSI-H tumour cell lines and integrate genomic profiling with biochemical reconstitution and isogenic modelling to define the molecular basis and functional consequences of resistance. We then evaluate whether resistance-associated cross-sensitivity can be translated into a rational treatment strategy in vivo, with an emphasis on long-term tumour control. Finally, motivated by the possibility that cross-resistance bottlenecks may arise during resistance evolution, we explore a structure-informed, AI-enabled design framework for rapidly generating next-generation WRN inhibitor candidates designed to retain activity against resistance-associated WRN variants. Collectively, this work aims to advance WRNi resistance research beyond mutation cataloguing toward resistance-informed treatment design and next-generation inhibitor development, with the long-term goal of enabling more durable WRN-targeted therapies for MSI-H/dMMR cancers.

## Results

### Parallel resistance models reveal durable escape from clinically relevant WRN inhibitor classes

To define the molecular basis of acquired resistance to WRN inhibition and to enable a head-to-head comparison of resistance evolution under distinct inhibitory modalities, we generated parallel panels of stable resistance models to a clinically relevant non-covalent WRN inhibitor (HRO761) and a covalent WRN inhibitor (VVD-214)^6,8^. We focused on three MSI-H tumour cell lines—HCT116, SW48, and Ishikawa—that are highly sensitive to WRN inhibition^6,8^. Resistant populations were established using a long-term selection strategy with alternating high-dose and low-dose drug exposure^32,33^, yielding six HRO761-resistant populations and four VVD-214-resistant populations (Fig. 1a).

**Figure 1.**
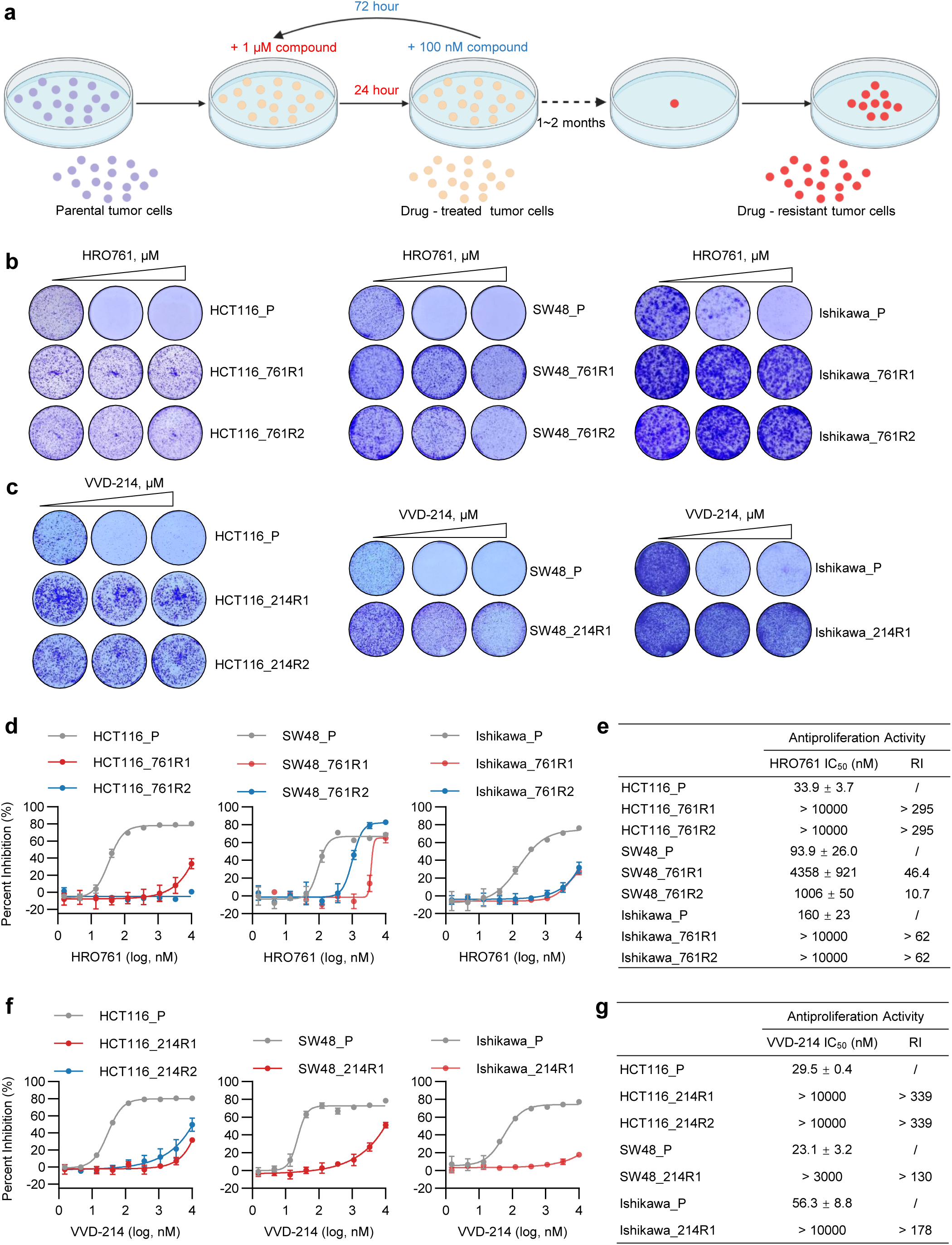
Generation of parallel acquired-resistance models to non-covalent and covalent WRN inhibitors in MSI-H cells. **a,** Schematic overview of the generation of WRN inhibitor-resistant cells. Resistant populations were established by cyclic drug exposure: cells were treated with high-dose inhibitor for 24 h, followed by low-dose inhibitor treatment for 72 h. This cycle was repeated for 1-2 months until stable resistant populations were successfully obtained. **b, c,** Colony formation assays assessing the clonogenic capacity of parental cells and the corresponding resistant cells after treatment with HRO761 (**b**) or VVD-214 (**c**) at 0, 0.3 and 1 μM for 5 days. **d–g,** Antiproliferative activity of HRO761 (**d, e**) or VVD-214 (**f, g**) in parental cells and the corresponding resistant cells. **d** and **f** show dose–response curves, and **e** and **g** summarize the IC₅₀ values and resistance index (RI). Cells were treated with the indicated compounds for 4 days. RI was defined as the IC₅₀ of resistant cells divided by the IC₅₀ of the corresponding parental cells. Experiments were independently repeated three times. Data are presented as mean ± standard deviation (SD).

We first confirmed that these selected populations exhibited a stable and robust resistant phenotype. In clonogenic assays, HRO761 or VVD-214 markedly suppressed colony formation in parental cells, whereas the corresponding resistant populations maintained strong colony-forming capacity under the same treatment conditions; even at higher drug concentrations, colony growth was only modestly affected in most resistant cells (Fig. 1b, c). Consistently, short-term proliferation assays showed substantially decreased sensitivity to the selecting inhibitor across all resistant lines. For HRO761, parental HCT116, SW48, and Ishikawa cells displayed IC₅₀ values of 33.9 ± 3.7 nM, 93.9 ± 26.0 nM, and 160 ± 23 nM, respectively, whereas the matched resistant populations showed markedly elevated IC₅₀ values, with most exceeding 10,000 nM (Fig. 1d, e). For VVD-214, parental IC₅₀ values were 29.5 ± 0.4 nM, 23.1 ± 3.2 nM, and 56.3 ± 8.8 nM, respectively, while the corresponding resistant populations generally shifted into the micromolar range; notably, HCT116_214R1, HCT116_214R2, and Ishikawa_214R1 exhibited IC₅₀ values greater than 10,000 nM (Fig. 1f, g).

Overall, all resistant populations showed at least a 10-fold increase in IC₅₀ relative to their parental counterparts, with some exhibiting more than a 300-fold increase. These modality-matched panels therefore provided a controlled experimental system to ask the central translational question of this study: whether resistance selected by one WRN inhibitor class creates vulnerabilities that can be exploited by the alternate class, or instead converges on cross-resistant bottlenecks that require next-generation inhibitor design.

### Resistant cells fail to mount the canonical WRNi-induced pharmacodynamic cascade

To define the phenotypic consequence of acquired resistance in a manner that can be linked to inhibitor action, we profiled WRNi-induced pharmacodynamic responses in parental and resistant cells at the levels of DDR signalling, G_2_/M cell cycle arrest, and downstream functional outcomes. Because activation of the DDR is a hallmark downstream consequence of effective WRN inhibition in MSI-H contexts^6,8^, we first compared DDR pathway engagement following treatment with HRO761 or VVD-214.

In parental cells, both inhibitors triggered a clear dose-dependent pharmacodynamic response. WRN protein abundance decreased markedly with increasing inhibitor concentrations, consistent with previous observations that WRN inhibition can promote chromatin retention of WRN accompanied by reduced WRN protein levels^34^. Because WRN reduction is a downstream pharmacodynamic readout rather than a direct phenotypic measurement, we interpreted it together with DDR activation, including induction of γ-H2AX and activation of the ATM–CHK2 and p53–p21 axes. In resistant populations, WRN protein levels were largely maintained upon drug treatment, and γ-H2AX induction as well as ATM–CHK2 and p53–p21 activation were substantially attenuated or abolished (Fig. 2a, b; Supplementary Fig. S1a–e). These results indicate that acquired resistance is characterized by failure to mount the canonical WRNi-induced pharmacodynamic cascade.

**Figure 2.**
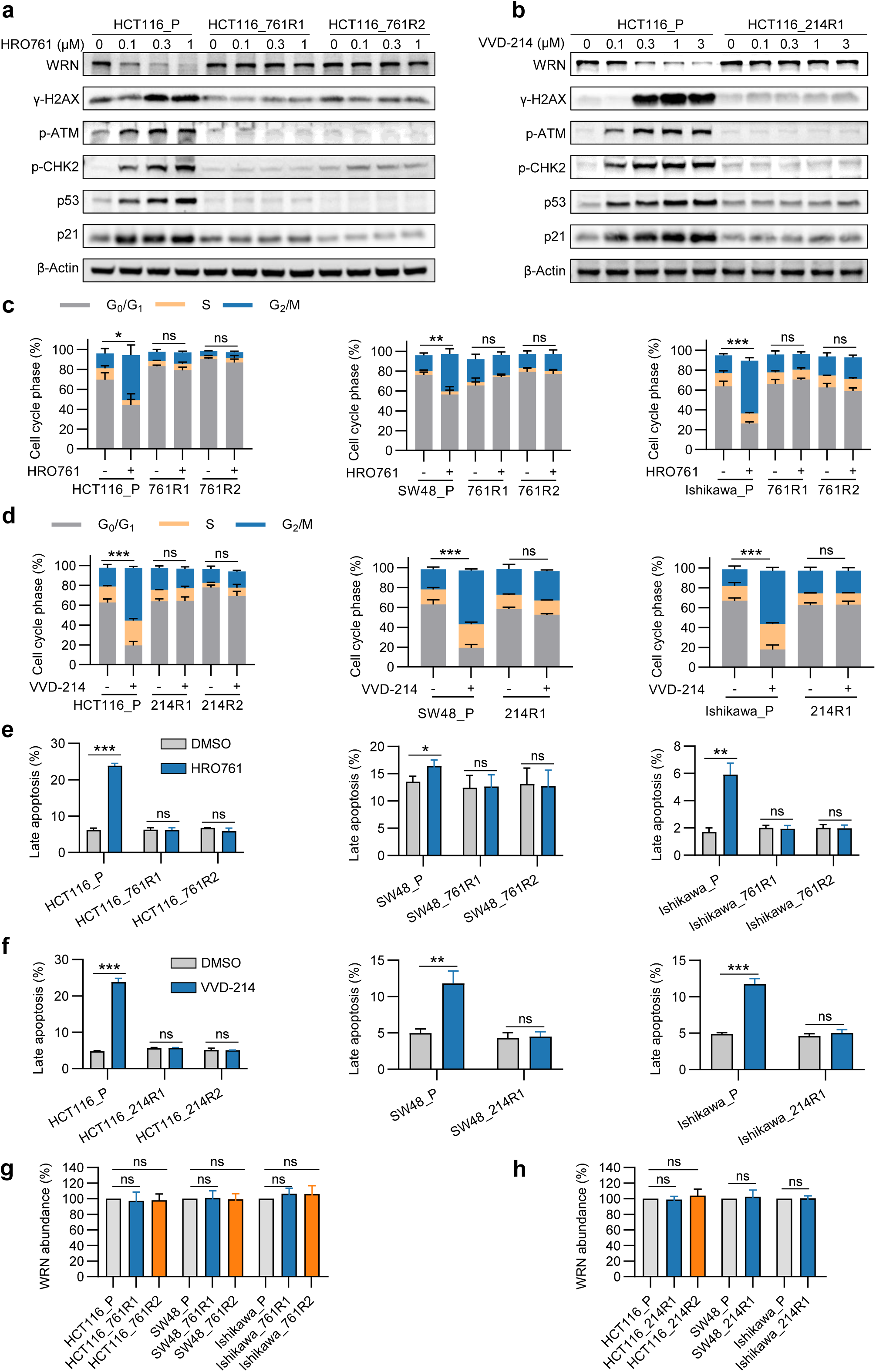
Acquired resistance reflects loss of on-target WRN inhibitor pharmacodynamics, blunting WRN protein reduction, DDR activation, G_2_/M cell cycle arrest and apoptosis. **a, b,** Western blot analysis of WRN and DDR pathway-related proteins in parental cells and resistant cells after treatment with increasing concentrations of HRO761 (**a**) or VVD-214 (**b**) for 24 h. The analyzed proteins included WRN, γ-H2AX (Ser139), p-ATM (Ser1981), p-CHK2 (Thr68), p53, and p21. **c, d,** Flow cytometric analysis of cell-cycle distribution in parental cells and resistant cells after treatment with 0.3 μM HRO761 (**c**) or 0.3 μM VVD-214 (**d**) for 24 h. Experiments were independently repeated three times. **e, f,** Flow cytometric analysis of late apoptosis in parental cells and resistant cells after treatment with 0.3 μM HRO761 (**e**) or 0.3 μM VVD-214 (**f**) for 24 h. Experiments were independently repeated three times. **g, h,** Western blot analysis of basal WRN protein levels in parental cells and resistant cells. Protein band intensities were quantified by densitometric analysis. Experiments were independently repeated three times. Data are presented as mean ± SD. *P < 0.05, **P < 0.01, ***P < 0.001; ns, not significant.

We next asked whether this loss of DDR engagement translated into downstream functional escape from WRN inhibitor-induced growth suppression. Consistent with effective DDR activation, parental cells displayed pronounced G₂/M accumulation and an increased fraction of late apoptotic cells following inhibitor exposure. By contrast, resistant cells did not exhibit significant G₂/M accumulation under the same treatment conditions (Fig. 2c, d), and late apoptosis was not significantly increased (Fig. 2e, f), indicating a coordinated loss of both G_2_/M cell cycle arrest and apoptotic commitment in the resistant state.

Finally, to exclude baseline WRN overexpression as a trivial explanation for reduced drug responsiveness, we compared basal WRN protein abundance between parental and resistant cells and observed no obvious differences (Fig. 2g, h; Supplementary Fig. S1f, g). Collectively, these data establish a shared resistance-associated phenotype across independently derived models: WRN inhibitors no longer elicit the expected pharmacodynamic cascade—WRN protein reduction, DDR activation, G_2_/M cell cycle arrest and apoptosis—thereby motivating a mechanistic search for genetic and biochemical alterations that disrupt inhibitor engagement.

### Non-covalent and covalent WRN inhibitors select distinct on-target resistance spectra

To identify the genetic determinants of acquired resistance to WRN inhibitors—and to test whether non-covalent and covalent WRN inhibitor modalities impose distinct selective pressures—we performed whole-exome sequencing on all ten independently derived resistant populations. Given the pronounced genomic instability and background heterogeneity characteristic of MSI-H tumours^11–13^, we focused on recurrent coding variants and compared the distribution of altered genes across resistance models to prioritize candidate events that consistently track with stable resistance.

Despite the highly heterogeneous mutational landscape, resistance evolution converged strongly on WRN. Among the top 30 coding variants ranked by variant frequency, WRN was the only gene mutated across all ten resistant populations, and all detected WRN alterations were missense mutations. By contrast, variants in other genes were observed only sporadically and at markedly lower recurrence frequencies, lacking consistent enrichment across models (Fig. 3a). These results indicate that recurrent on-target WRN alterations are the predominant events associated with acquired resistance in these populations, highlighting WRN missense mutations as the dominant candidate drivers of the resistant phenotype.

**Figure 3.**
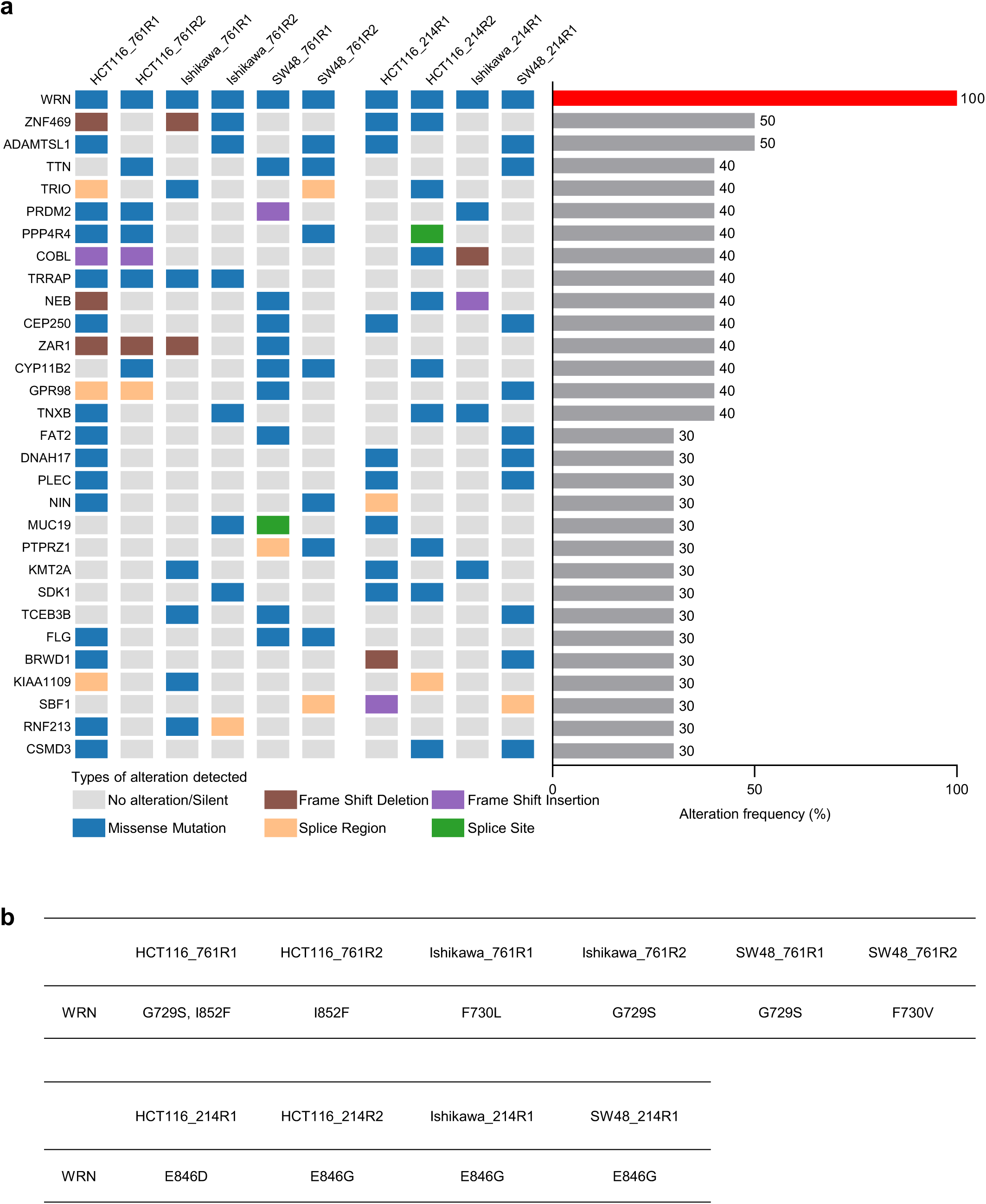
Whole-exome sequencing reveals modality-specific, on-target WRN mutation spectra underlying acquired resistance. **a,** Mutation types and alteration frequencies of the top 30 genes across ten WRN inhibitor-resistant cells. “No alteration/Silent” indicates either no detectable coding alteration or a synonymous variant. Other categories include missense mutation, frameshift deletion, frameshift insertion, splice-region mutation, and splice-site mutation. **b,** Distribution of WRN missense mutation sites in HRO761- and VVD-214-resistant cells.

Importantly, WRN mutations displayed clear inhibitor-modality–specific positional preferences. In HRO761-resistant populations, WRN mutations clustered predominantly at G729, F730, and I852, with several mutation types recurring across independent selections. In contrast, VVD-214-resistant populations exhibited WRN mutations exclusively at E846, most commonly E846G (Fig. 3b). This striking segregation of mutation sites suggests that non-covalent versus covalent WRN inhibition constrains resistance evolution through distinct molecular trajectories, suggesting the possibility that resistance mutations selected by one inhibitor class may preserve susceptibility to the other.

Together, these exome data define WRN missense mutations as recurrent on-target resistance events and reveal that HRO761 and VVD-214 impose distinct genetic routes to escape. This modality-specific architecture provided the genetic basis for testing whether resistance is functionally orthogonal and therefore therapeutically actionable through inhibitor switching or alternating schedules.

### Resistance mutations impair inhibitor engagement and are sufficient to disable WRNi pharmacodynamics and drive resistance

To establish a causal link between recurrent WRN missense mutations and the resistant phenotype, and to determine whether these alterations act by directly impairing inhibitor engagement, we evaluated resistance-associated WRN variants using complementary biochemical and isogenic cellular systems. We first expressed and purified wild-type (WT) WRN and the corresponding mutant proteins, and assessed inhibitor binding by protein thermal shift assays. Both HRO761 and VVD-214 markedly increased the thermal stability of WT WRN, consistent with direct target engagement. In contrast, the inhibitor-induced ΔT_m_ values were substantially reduced for the matched resistance-associated WRN mutants, with stabilization nearly abolished for some variants (Fig. 4a, b), indicating that these mutations weaken the interaction between WRN and the selecting inhibitor.

**Figure 4.**
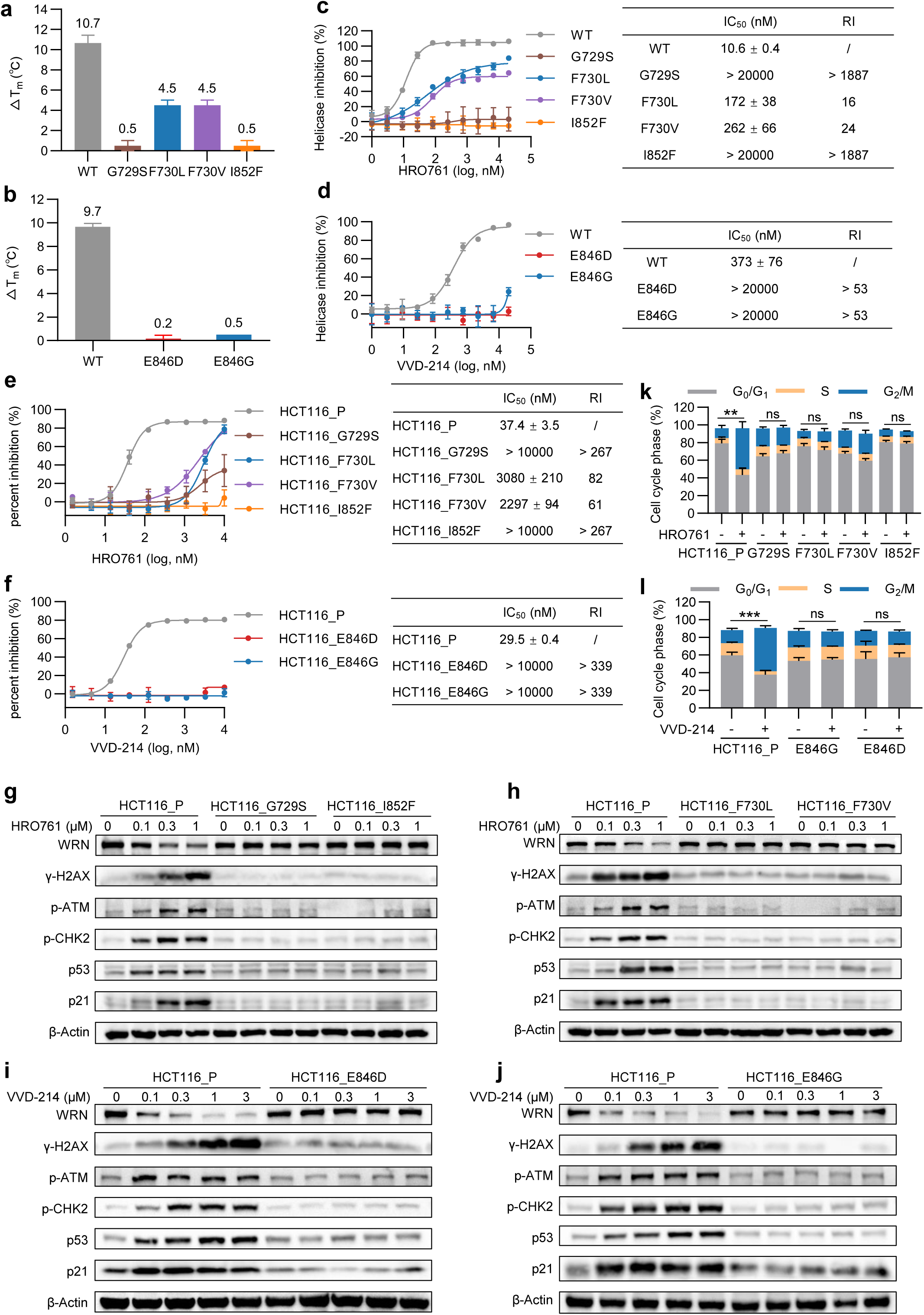
On-target WRN point mutations disrupt inhibitor engagement and are sufficient to abolish WRN inhibitor pharmacodynamics and drive resistance. **a, b,** Protein thermal shift assays showing inhibitor-induced thermal stabilization of WT and mutant WRN proteins after treatment with 5 μM HRO761 (**a**) or VVD-214 (**b**). T_m_ indicates protein melting temperature. ΔT_m_ values were calculated relative to the DMSO control. Experiments were independently repeated three times. **c, d,** Helicase activity inhibition assays assessing the inhibitory effects of HRO761 (**c**) or VVD-214 (**d**) on WT WRN and the corresponding WRN mutants. IC₅₀ was defined as the compound concentration required to inhibit 50% of WRN helicase activity. RI was calculated relative to WT WRN. Experiments were independently repeated three times. **e, f,** Antiproliferative activity of HRO761 (e) or VVD-214 (f) in WRN point-mutation knock-in cells and parental HCT116 cells (HCT116_P). Cells were treated with the indicated compounds for 4 days. RI was defined as the IC₅₀ of WRN point-mutation knock-in cells divided by the IC₅₀ of parental HCT116 cells. Experiments were independently repeated three times. **g–j,** Western blot analysis of WRN and DDR-related proteins in WRN point mutation knock-in cells and parental HCT116 cells after treatment with increasing concentrations of HRO761 (**g, h**) or VVD-214 (**i, j**) for 24 h. The analyzed DDR-related proteins included γ-H2AX (Ser139), p-ATM (Ser1981), p-CHK2 (Thr68), p53, and p21. **k, l,** Flow cytometric analysis of cell-cycle distribution in WRN point mutation knock-in cells and parental HCT116 cells after treatment with 0.3 μM HRO761 (**k**) or 0.3 μM VVD-214 (**l**) for 24 h. Experiments were independently repeated three times. Data are presented as mean ± SD. **P < 0.01, ***P < 0.001; ns, not significant.

We next asked whether reduced engagement translates into impaired functional inhibition of WRN helicase activity. In in vitro helicase assays, resistance-associated WRN mutations diminished inhibition by the corresponding inhibitors to varying degrees. For HRO761-selected variants, G729S and I852F nearly eliminated HRO761-mediated inhibition (IC₅₀ > 20,000 nM; RI > 1887), while F730L and F730V also substantially reduced HRO761 potency (RI = 16 and 24, respectively) (Fig. 4c). For VVD-214-selected variants, E846D and E846G similarly impaired VVD-214 inhibition (IC₅₀ > 20,000 nM; RI > 53) (Fig. 4d). These results support a model in which on-target WRN substitutions directly compromise inhibitor-mediated enzymatic suppression.

To test sufficiency in a controlled genetic background, we generated isogenic HCT116 knock-in cell lines carrying single WRN point mutations (G729S, F730L, F730V, I852F, E846D, and E846G). In proliferation assays, each mutant line exhibited markedly reduced sensitivity to the selecting WRN inhibitor. Under HRO761 treatment, G729S and I852F conferred high-level resistance (RI > 267), and F730L and F730V also produced pronounced resistance (RI = 82 and 61, respectively) (Fig. 4e). Under VVD-214 treatment, E846D and E846G likewise conferred high-level resistance (RI > 339) (Fig. 4f). Thus, introduction of a single WRN point mutation is sufficient to drive resistance in an otherwise matched cellular context.

Finally, because a key objective of this study is to connect resistance genotypes to actionable pharmacodynamic consequences, we examined whether these WRN point mutations disrupt the canonical WRNi response program. Whereas parental HCT116 cells showed dose-dependent WRN protein reduction, γ-H2AX induction, and activation of the ATM–CHK2 and p53–p21 pathways following WRN inhibitor treatment, each WRN mutant cell line displayed markedly attenuated or abolished responses (Fig. 4g–j). Consistently, cell-cycle profiling showed robust G₂/M accumulation in parental cells but not in the mutant lines (Fig. 4k, l). Together, these biochemical and isogenic data establish WRN missense mutations as direct on-target resistance determinants that impair inhibitor engagement, reduce enzymatic inhibition, and abrogate WRNi-induced pharmacodynamic responses—providing a mechanistic foundation for subsequent analyses of whether resistance mutations selected by one inhibitor modality retain functional sensitivity to the alternative modality.

### Modality-specific resistance creates actionable cross-sensitivity to the alternate WRN inhibitor class

The above results showed that HRO761 and VVD-214 select distinct WRN resistance mutation spectra, raising the possibility that these inhibitors impose different selective pressures and that resistance to one inhibitor class may retain sensitivity to the other. To contextualize these inhibitor-class–specific preferences, we mapped resistance-associated residues onto previously reported structural information. HRO761 primarily engages the D1–D2 ATPase region of the WRN helicase domain^6^, consistent with the proximity of HRO761-associated resistance residues G729, F730, and I852 to this region (Fig. 5a). In contrast, VVD-214 acts as a covalent allosteric inhibitor engaging Cys727 within the ATPase domain hinge region^8^, and its major resistance-associated residue E846 localizes to this structural context (Fig. 5b). These observations provide a structural rationale for the divergent resistance-site preferences observed across the two inhibitor modalities.

**Figure 5.**
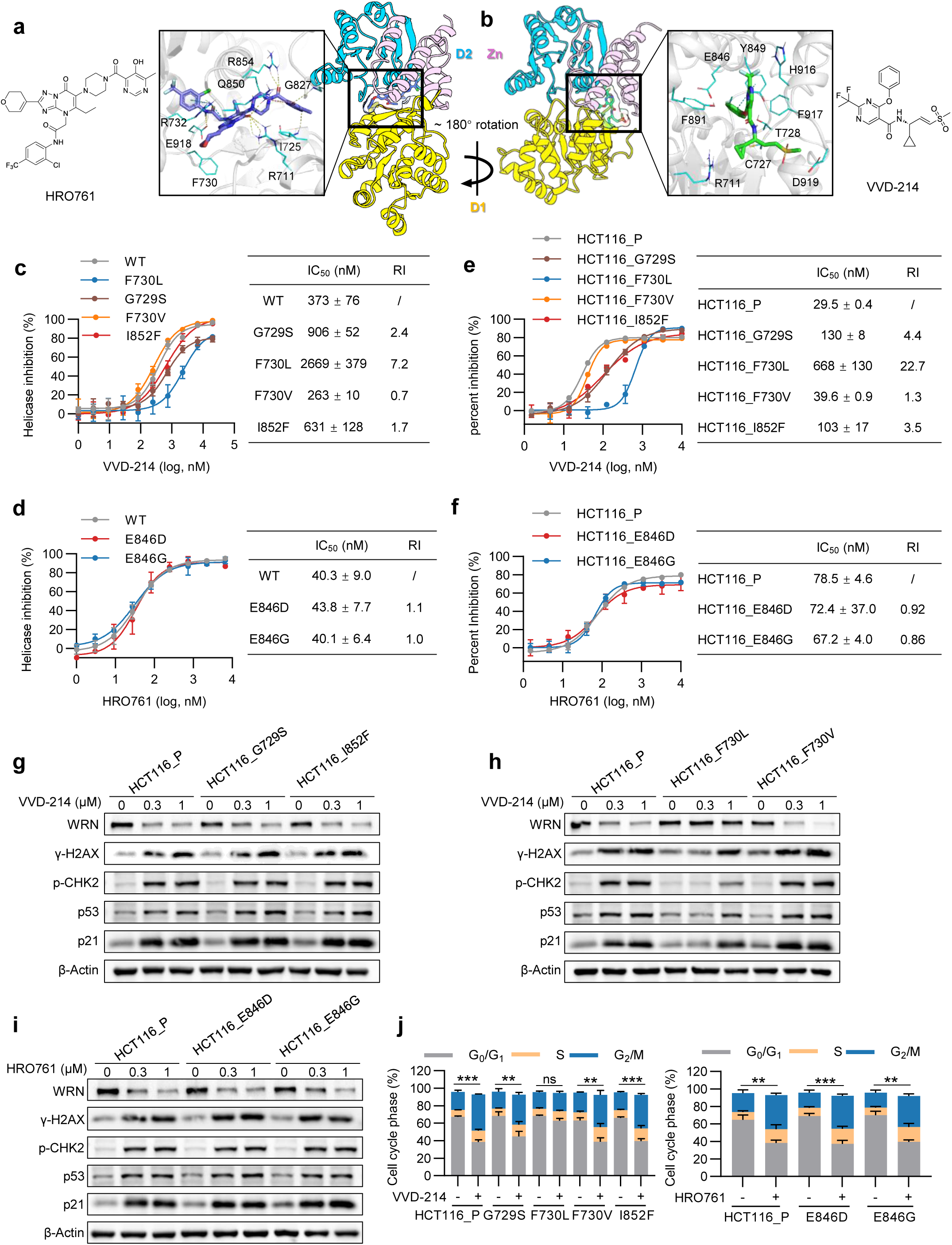
Orthogonal functional resistance profiles enable cross-sensitivity between non-covalent and covalent WRN inhibitors. **a, b,** Binding modes of HRO761 (**a**) and VVD-214 (**b**) on WRN based on previously reported structural information. **c, d,** Helicase activity inhibition assays assessing the inhibitory activity of VVD-214 against HRO761-associated WRN mutants (**c**) and HRO761 against VVD-214-associated WRN mutants (**d**). Experiments were independently repeated three times. **e, f,** Antiproliferative activity of VVD-214 (**e**) or HRO761 (**f**) in WRN point-mutation knock-in cells associated with resistance to HRO761 or VVD-214 respectively. Cells were treated with the indicated compounds for 4 days. Experiments were independently repeated three times. **g–i,** Western blot analysis of WRN and DDR-related proteins, including γ-H2AX (Ser139), p-CHK2 (Thr68), p53, and p21, in WRN point mutation knock-in cells after treatment with increasing concentrations of VVD-214 (**g, h**) or HRO761 (**i**) for 24 h. β-Actin was used as the loading control. **j,** Flow cytometric analysis of cell-cycle distribution in WRN point mutation knock-in cells after treatment with DMSO, 0.3 μM HRO761, or 0.3 μM VVD-214 for 24 h. Experiments were independently repeated three times. Data are presented as mean ± SD. **P < 0.01, ***P < 0.001; ns, not significant.

We next asked whether resistance-associated WRN variants compromise only the selecting inhibitor or also confer cross-resistance, and thus whether an orthogonal functional resistance profile exists between non-covalent and covalent WRN inhibition. In vitro helicase inhibition assays showed that VVD-214 largely retained inhibitory activity against the HRO761-associated mutants G729S, F730V, and I852F, with RI below 3, whereas its inhibitory activity against the engineered F730L mutant was markedly reduced (RI = 7.2) (Fig. 5c). Conversely, HRO761 fully retained inhibitory activity against the VVD-214-associated mutants E846D and E846G, with RIs of approximately 1 (Fig. 5d). Thus, with the exception of the F730L bottleneck model, the tested resistance mutations did not confer complete cross-resistance at the biochemical level.

Cell proliferation assays further supported this cross-sensitivity pattern. VVD-214 maintained potent antiproliferative activity against HCT116_G729S, HCT116_F730V, and HCT116_I852F cells, with RIs below 5, whereas HCT116_F730L cells showed markedly reduced sensitivity to VVD-214 (RI = 22.7) (Fig. 5e). In parallel, HRO761 retained strong antiproliferative activity against HCT116_E846D and HCT116_E846G cells, with RIs of approximately 1 (Fig. 5f). These data indicate that most resistance mutations selected by one inhibitor class remain functionally tractable to the alternate inhibitor class at the cellular level, again with the engineered F730L bottleneck model representing a notable exception.

To determine whether cross-sensitivity reflected restoration of on-target pathway engagement rather than isolated viability effects, we examined downstream DDR activation and cell cycle arrest under cross-treatment conditions. Following VVD-214 treatment, γ-H2AX, p-CHK2, p53, and p21 remained upregulated in HCT116_G729S, HCT116_F730V, and HCT116_I852F cells with responses comparable to parental cells, indicating preserved DDR activation; by contrast, in HCT116_F730L cells, only 1 μM VVD-214 induced mild DDR activation (Fig. 5g, h). Conversely, after HRO761 treatment, DDR markers were similarly induced in HCT116_E846D and HCT116_E846G cells (Fig. 5i). Consistently, cell-cycle profiling showed that cross-treatment with the alternate inhibitor significantly induced G_2_/M-phase arrest in the tested mutant cells, except for the F730L context (Fig. 5j).

Together, these results move the resistance map from genotype to therapeutic logic. Across most resistance-associated WRN variants, mutations that compromised the selecting inhibitor preserved sensitivity to the alternate modality in biochemical inhibition, cellular antiproliferative activity and re-engagement of WRNi-linked DDR activation and G_2_/M arrest. This functional orthogonality provided a mechanistic foundation for resistance-informed inhibitor switching and for the cyclic alternating treatment strategy tested in vivo. F730L was included as an engineered stress-test mutation to model a plausible cross-resistant escape route at a recurrent resistance hotspot, and its clinical frequency should be defined as WRNi-treated patient samples become available.

### Functional orthogonality enables mutation-informed switching and cyclic alternating therapy in vivo

To determine whether the cross-sensitivity and functional orthogonality observed in vitro can be maintained in vivo, we established HCT116 cell-derived xenograft models carrying distinct WRN point mutations and treated them with either HRO761 or VVD-214. Both inhibitors significantly suppressed the growth of WT WRN tumours (Fig. 6a). Consistent with inhibitor-class–specific resistance, HRO761 showed markedly reduced or nearly abolished antitumour activity against tumours harbouring HRO761-associated resistance mutations (G729S, F730V, and I852F), yet retained significant efficacy against tumours carrying VVD-214-associated resistance mutations (E846D and E846G). Conversely, VVD-214 exhibited markedly reduced activity against E846D- and E846G-mutant tumours, while maintaining clear antitumour activity against G729S-, F730V-, and I852F-mutant tumours. In contrast, engineered F730L-mutant tumours displayed pronounced resistance to both inhibitors (Fig. 6b-g). Together, these data support the notion that, for on-target resistance contexts, inhibitor-class switching can restore antitumour activity in vivo, whereas a subset of resistance contexts may compromise both modalities.

**Figure 6.**
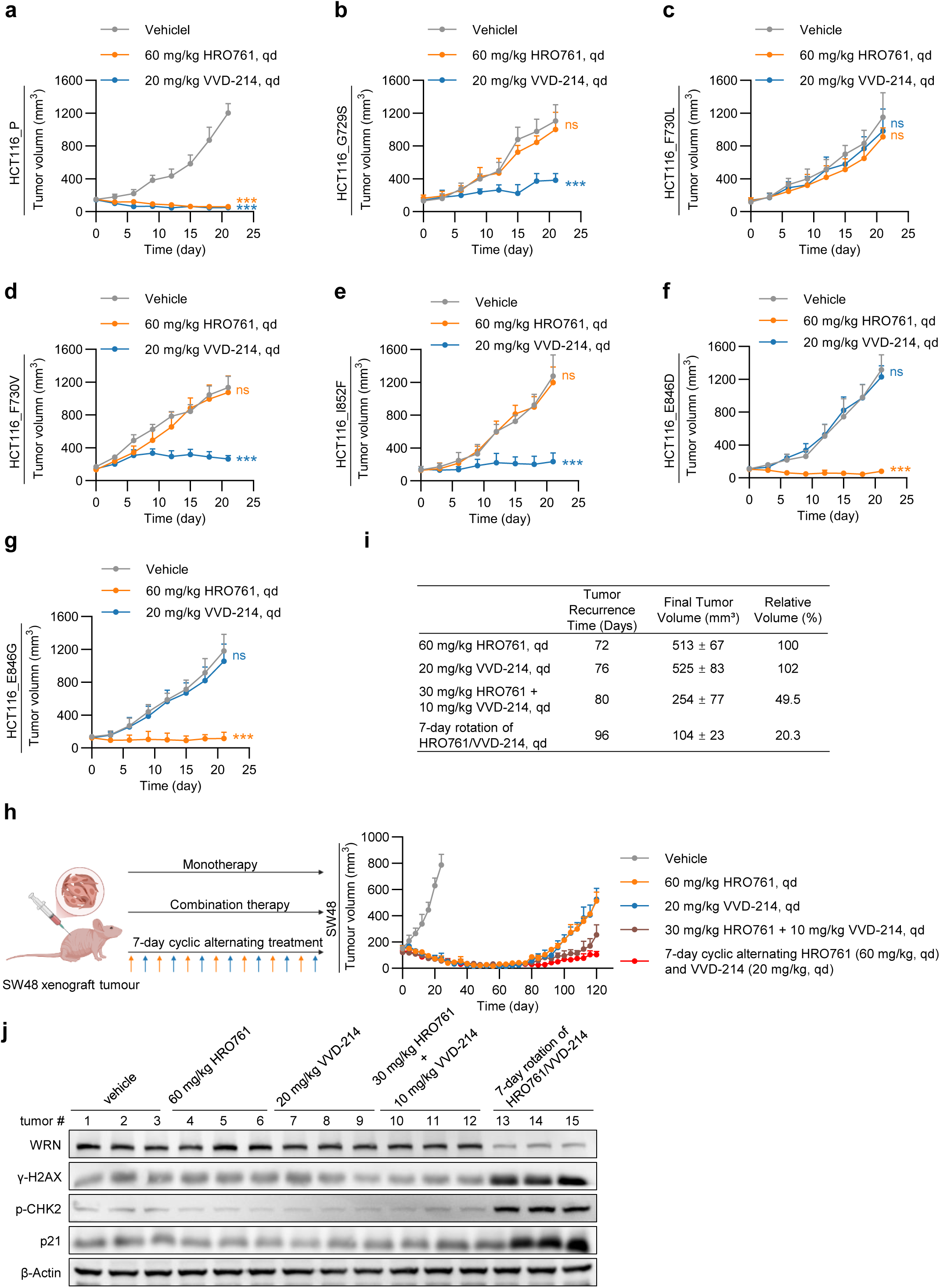
Orthogonal resistance profiles translate into mutation-informed switching and cyclic alternating therapy to extend tumour control in vivo. **a–g,** Tumour growth curves of WT and WRN point-mutant HCT116 xenografts (n = 6) treated with HRO761 monotherapy (60 mg/kg, qd) or VVD-214 monotherapy (20 mg/kg, qd). qd, once daily. All treatments were administered by oral gavage. Statistical comparisons were made against the vehicle group at the endpoint unless otherwise indicated. ***P < 0.001; ns, not significant. **h,** Tumour growth curves of SW48 xenografts (n = 6) under different long-term treatment regimens. Monotherapy groups received HRO761 (60 mg/kg, qd) or VVD-214 (20 mg/kg, qd). The combination group received HRO761 (30 mg/kg, qd) plus VVD-214 (10 mg/kg, qd). The 7-day cyclic alternating treatment group alternated every 7 days between HRO761 (60 mg/kg, qd) and VVD-214 (20 mg/kg, qd). All treatments were administered by oral gavage. **i,** Summary of tumour regrowth time, endpoint tumour volume, and relative tumour volume compared with the HRO761 monotherapy group in the long-term efficacy study. Tumour regrowth was defined as an increase in tumour volume in two consecutive measurements compared with the previous measurement point, exceeding the minimum tumour volume recorded during treatment. **j,** Western blot analysis assessed expression levels of WRN and DDR-related proteins, including γ-H2AX (Ser139), p-CHK2 (Thr68), and p21, in endpoint tumour tissues from the long-term efficacy study. Data are presented as mean ± SD.

To link these efficacy patterns to on-target pathway engagement in tumours, we assessed WRN abundance and DDR-associated markers following treatment. Molecular responses were broadly aligned with tumour growth outcomes: in treatment-responsive settings, WRN protein levels were markedly reduced and DDR markers (γ-H2AX, p-CHK2, and p21) were induced, whereas in non-responsive settings these outputs were attenuated or not effectively triggered. Specifically, HRO761 induced pronounced WRN downregulation and DDR activation in E846D/E846G-mutant tumours but had markedly reduced effects in G729S/F730V/I852F-mutant tumours, while VVD-214 showed the opposite pattern. In engineered F730L-mutant tumours, neither inhibitor induced obvious WRN reduction or DDR activation (Supplementary Fig. S2a–g), consistent with the observed in vivo resistance. These findings indicate that, with the exception of the F730L bottleneck model, tumours carrying on-target resistance mutations selected by one WRN inhibitor can retain functional sensitivity to the alternate inhibitor modality in vivo.

Given prior reports of acquired resistance and tumour regrowth under continuous HRO761 treatment in SW48 xenografts^6^, we next asked whether orthogonal resistance landscapes could be leveraged proactively to extend tumour control in a long-term setting. We compared three dosing strategies in the SW48 model: continuous monotherapy, combination therapy, and a 7-day cyclic alternating regimen. In the combination group, HRO761 and VVD-214 were co-administered at half of their respective monotherapy doses. Long-term efficacy studies showed that tumours in the HRO761 and VVD-214 monotherapy arms regrew on days 72 and 76 after treatment initiation, respectively, with endpoint tumour volumes of 513 ± 67 mm³ and 525 ± 83 mm³. Tumours in the combination group regrew on day 80, with an endpoint volume of 254 ± 77 mm³ (49.5% of the HRO761 monotherapy endpoint). Strikingly, tumour regrowth in the 7-day cyclic alternating group was delayed until day 96, with an endpoint volume of 104 ± 23 mm³ (20.3% of the HRO761 monotherapy endpoint) (Fig. 6h, i).

To assess whether the improved long-term outcome under alternating therapy was accompanied by sustained on-target pathway engagement, we examined WRN inhibition-associated molecular responses in endpoint tumours. Compared with vehicle controls, only the cyclic alternating group retained pronounced WRN protein reduction together with sustained activation of DDR markers, including γ-H2AX, p-CHK2, and p21 (Fig. 6j). Collectively, these in vivo results demonstrate that inhibitor-class–specific resistance can remain therapeutically exploitable in tumour settings. They support two linked principles: mutation-informed switching can restore antitumour activity for many on-target resistance contexts, and proactive cyclic alternation of mechanistically distinct WRN inhibitors can extend tumour control while preserving endpoint WRNi pharmacodynamic responses.

### F730L supports a hinge-remodelling model for cross-resistance and guides mutation-aware inhibitor design

The above results established an orthogonal functional resistance landscape between non-covalent and covalent WRN inhibition for many resistance contexts, yet also revealed that the engineered F730L model compromises both modalities and may represent a potential cross-resistant bottleneck. Notably, F730L conferred marked resistance to both HRO761 and VVD-214, whereas F730V lost sensitivity to HRO761 but retained responsiveness to VVD-214. Because these two substitutions differ only in the side-chain properties at residue 730 but display markedly distinct drug-response patterns, we hypothesized that the cross-resistant phenotype is unlikely to be explained by a single disrupted local contact. Instead, it may arise from broader hinge-region and pocket-environment remodelling that destabilizes productive inhibitor engagement, particularly for covalent inhibition that depends on maintaining a reaction-competent geometry.

To explore the structural basis of this differential sensitivity and to develop a mechanistic rationale for bottleneck formation, we constructed complexes of VVD-214 bound to the WRN helicase domain in the WT, F730V, and F730L contexts and performed 1 μs molecular dynamics simulations. At the protein level, the F730L system displayed increased conformational fluctuations and reduced compactness compared with WT and F730V, as reflected by a markedly increased protein backbone root-mean-square deviation (RMSD) (Fig. 7a). Residue-level root-mean-square fluctuation (RMSF) analysis further indicated increased protein flexibility in the F730L context, with pronounced effects in the D1–D2 hinge/linker region where residue 730 resides (Fig. 7b), suggesting perturbed interdomain coupling. Consistent with these observations, radius of gyration (Rg) analysis showed a broader distribution and an upward shift in the median value for the F730L system, indicating a more relaxed ensemble and reduced compactness (Fig. 7c).

**Figure 7.**
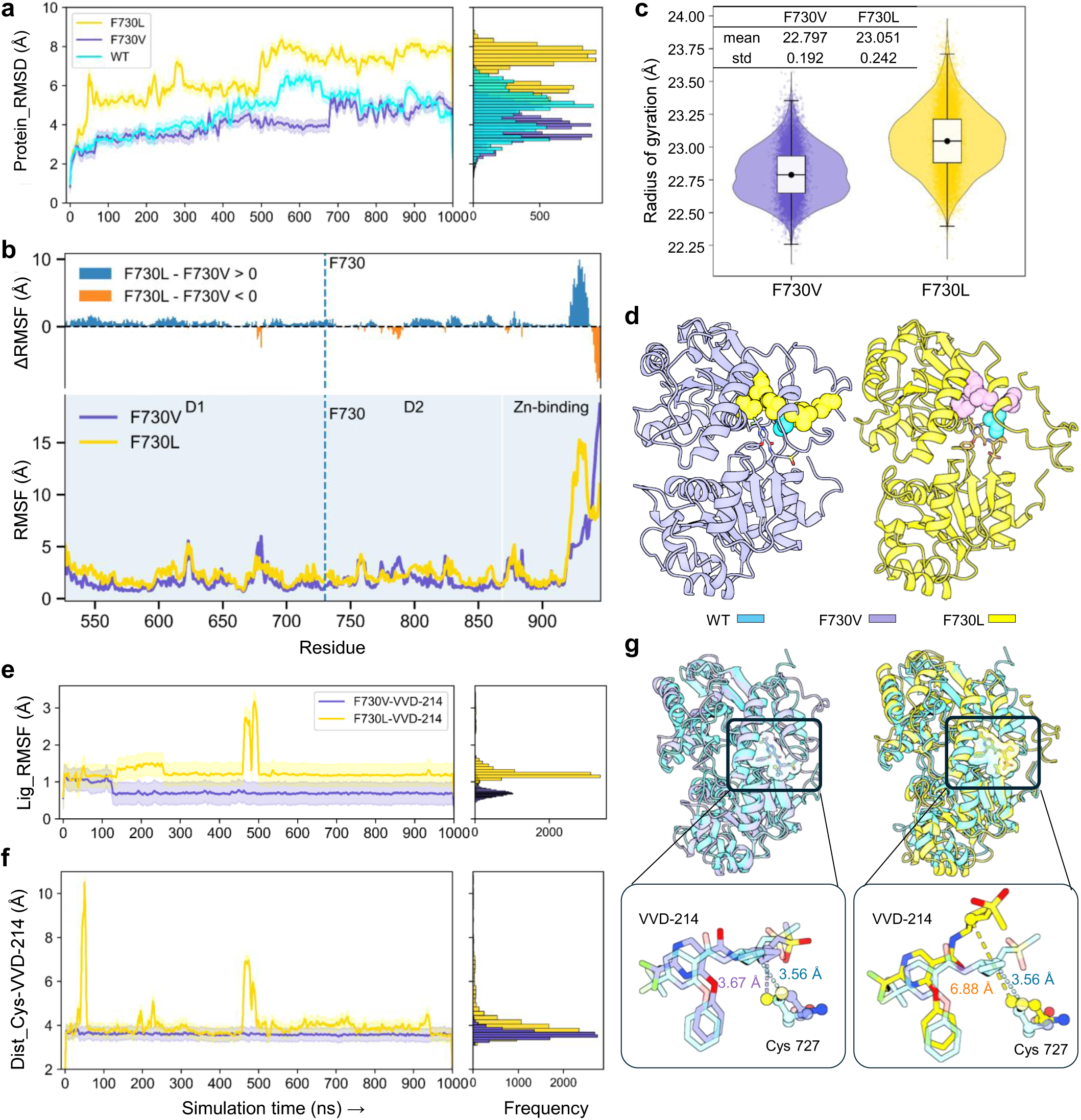
Hinge-region remodeling provides a structural–dynamic rationale for cross-resistant bottlenecks and guides next-generation inhibitor design. **a,** Protein backbone RMSD analysis showing increased global conformational fluctuations in the F730L system. **b,** RMSF analysis showing markedly increased flexibility in the D1–D2 hinge region of the F730L system. **c,** Rg distribution indicating a more relaxed conformation in the F730L system. **d,** Analysis of the local hydrophobic cluster surrounding F730 showing that the F730L mutation weakens the local hydrophobic core. **e,** Ligand RMSD analysis indicating reduced binding stability of VVD-214 in the F730L system. **f,** Time-dependent distance between the sulfur atom of Cys727 and the β-carbon of the acrylamide warhead, showing loss of the reaction-competent conformation in the F730L system. **g,** Comparison of representative binding conformations: WT and F730V on the left, and WT and F730L on the right.

We next examined local packing features that could account for this hinge destabilization. Analysis of the hydrophobic cluster surrounding residue 730 showed that the F730V context retained a relatively stable hydrophobic core comprising eight hydrophobic residues with a surface area of 741.4 Å², whereas the F730L substitution was associated with a reorganization and reduction of the local hydrophobic cluster, yielding a smaller cluster of six hydrophobic residues with a surface area of 611.3 Å² (Fig. 7d). These data support a model in which F730L remodels the hinge-adjacent hydrophobic environment, thereby reducing local structural stability and propagating increased conformational plasticity to the binding pocket.

At the ligand level, this destabilized protein environment was accompanied by reduced binding pose stability of VVD-214. Ligand RMSD remained low and stable in the F730V system (average RMSD of approximately 0.6 Å), whereas the F730L system displayed substantially larger ligand fluctuations (average RMSD of approximately 1.5 Å) together with evident conformational rearrangements during the simulation (Fig. 7e). Importantly, covalent inhibition requires the inhibitor to repeatedly sample—and maintain—a reaction-competent geometry. Accordingly, the distance between the sulfur atom of Cys727 and the β-carbon of the acrylamide warhead remained stably within a reaction-favorable range in the F730V system but fluctuated markedly in the F730L system, with excursions away from the reactive site (Fig. 7f). Representative conformations further indicated that while VVD-214 binding poses were largely consistent between WT and F730V, the ligand was displaced in the F730L context, moving the warhead away from Cys727 and disrupting geometries favorable for covalent engagement (Fig. 7g).

Collectively, these simulations support a structural–dynamic model in which hinge-region remodelling and destabilization of the local hydrophobic core can propagate to the inhibitor-binding environment, reduce ligand pose stability, and—critically for covalent modalities—impair maintenance of a reaction-competent geometry. These data provide a plausible mechanistic rationale for how a residue-730 substitution could create cross-resistance and motivate a next-generation design strategy that explicitly prioritizes pocket compatibility and mutation tolerance under hinge remodelling, rather than optimizing potency solely in the WT structural context.

### Mutation-aware, structure-guided optimization yields a proof-of-concept F730L-tolerant WRN inhibitor

The above results identified an engineered cross-resistant bottleneck model in which F730L conferred marked resistance to both HRO761 and VVD-214, motivating the development of WRN inhibitors capable of maintaining activity under this resistance context. Guided by the mechanistic analyses described above, we reasoned that F730L-mediated resistance is unlikely to arise from loss of a single local interaction. Instead, it may reflect a combination of (i) reduced conformational stability of the D1–D2 hinge region and adjacent binding pocket and (ii) remodelling of the local hydrophobic environment surrounding residue 730, which together decrease the compatibility of the original HRO761 scaffold with the mutant pocket. Accordingly, our design objective was not simply to optimize affinity in the WT background, but to improve pocket compatibility and reduce mutational penalty across both WT and F730L contexts, with particular attention to the ring scaffold proximal to residue 730 and its peripheral substituents.

Based on this rationale, we adopted a two-stage structure-guided optimization strategy to develop HRO761 derivatives with improved F730L tolerance. In the first stage, we enumerated ring scaffolds around the original oxygen-containing unsaturated six-membered ring and used PBCNet2.0^35^ to evaluate cross-system compatibility of 44 aromatic and saturated six-membered ring scaffolds in both WT and F730L WRN conformations. Rather than prioritizing candidates solely by predicted binding affinity in a single system, we jointly assessed mutation-induced binding penalties, binding-pose consistency, and local pocket occupancy in WT and F730L backgrounds, using HRO761 as the scoring reference. This screening identified four classes of aromatic and saturated six-membered ring scaffolds with improved tolerance to the mutant environment, which were advanced to second-stage optimization (Supplementary Table S1). Binding-mode analysis suggested that the para position of the HRO761 scaffold remained readily modifiable; therefore, in the second stage we introduced R-group substitutions at this position, generating a customized library of 103 molecules. PBCNet2.0 was used to score binding against both WT and F730L systems, and—together with synthesizability filters (synthetic accessibility (SA) score < 4 and RAscore > 0.7) and structural clustering—five candidates were selected for synthesis to balance accessibility and chemical diversity (Fig. 8a, Supplementary Table S1–S3). Among the synthesized compounds, GBA-007 showed the strongest activity in a preliminary helicase screen, achieving 85.9% and 92.9% inhibition of WRN helicase activity at 1 μM and 10 μM, respectively, outperforming the other candidates and HRO761 (Supplementary Fig. S3a). We therefore prioritized GBA-007 for detailed evaluation in WT and F730L models.

**Figure 8.**
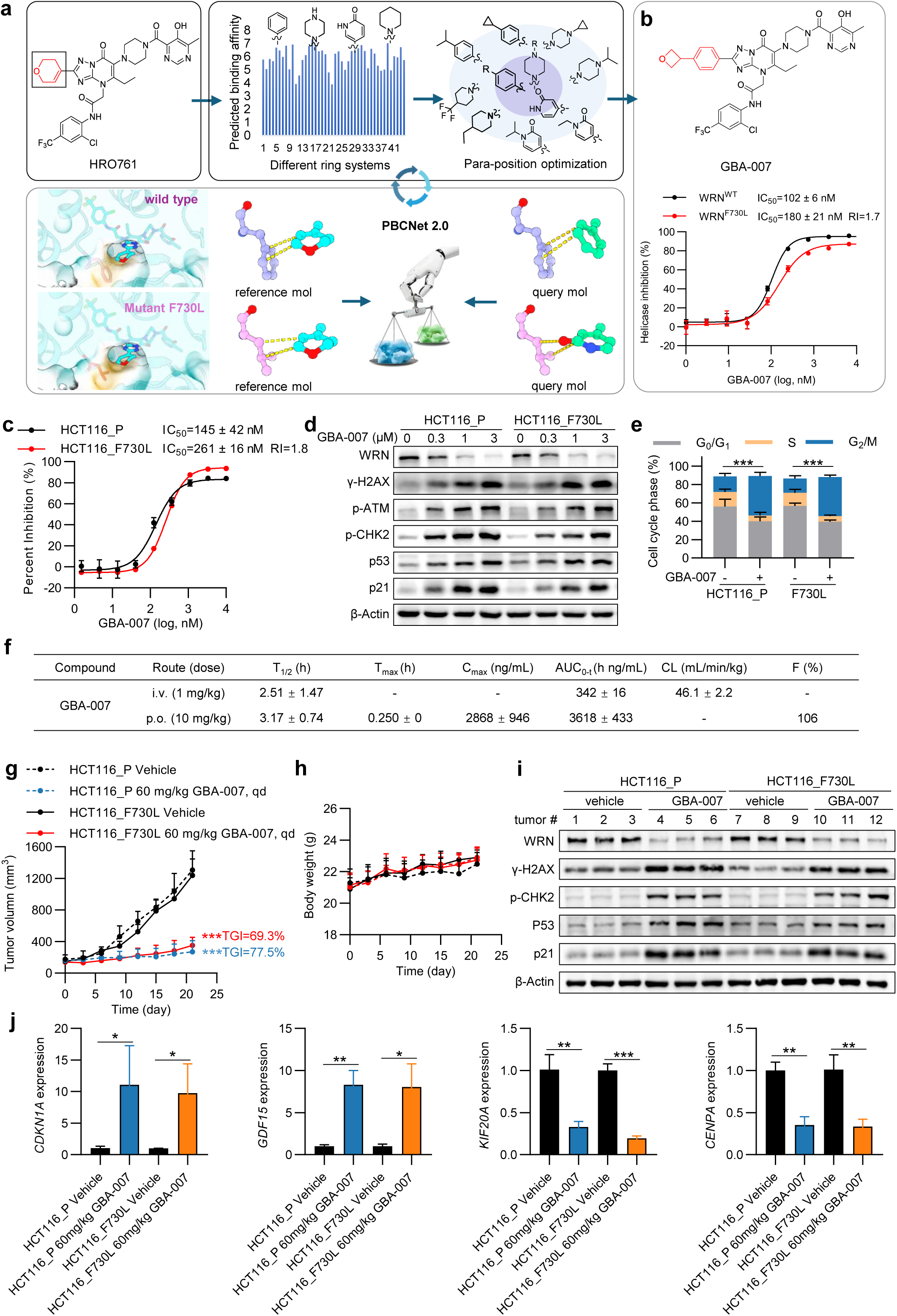
AI-enabled, structure-guided optimization yields a next-generation WRN inhibitor that overcomes a cross-resistant bottleneck. **a,** Schematic overview of the design workflow for the next-generation WRN inhibitor GBA-007. **b,** Helicase activity inhibition assays assessing the inhibitory activity of GBA-007 against WT WRN and the F730L mutant. IC₅₀ was defined as the compound concentration required to inhibit 50% of WRN helicase activity. Experiments were independently repeated three times. **c,** Antiproliferative activity of GBA-007 in HCT116_P and HCT116_F730L cells. Cells were treated with the indicated compound for 4 days. Experiments were independently repeated three times. **d,** Western blot analysis of WRN and DDR-related proteins in HCT116_P and HCT116_F730L cells after treatment with increasing concentrations of GBA-007 for 24 h. The analyzed DDR-related proteins included γ-H2AX (Ser139), p-ATM (Ser1981), p-CHK2 (Thr68), p53, and p21. β-Actin was used as the loading control. **e,** Flow cytometric analysis of cell-cycle distribution in HCT116_P and HCT116_F730L cells after treatment with 0.5 μM GBA-007 for 24 h. Experiments were independently repeated three times. **f,** Pharmacokinetic profile of GBA-007 in BALB/c mice (n = 3) after intravenous administration at 1 mg/kg and oral gavage at 10 mg/kg. i.v., intravenous administration; p.o., oral gavage; T₁_/_₂, terminal elimination half-life; Tₘₐₓ, time to maximum plasma concentration; Cₘₐₓ, maximum plasma concentration; AUC_0-t_, area under the plasma concentration–time curve from time zero to the last measurable time point; CL, clearance; F, oral bioavailability calculated from dose-normalized AUC_0-t_ values after p.o. and i.v. administration. -, not applicable. **g,h,** Tumour growth curves (**g**) and body weight changes (**h**) of HCT116_P and HCT116_F730L cell-derived xenograft models (n = 6) treated with GBA-007 at 60 mg/kg once daily by oral gavage. **i,** Western blot analysis of WRN and DDR-related proteins in xenograft tumour tissues after in vivo GBA-007 treatment. The analyzed DDR-related proteins included γ-H2AX (Ser139), p-CHK2 (Thr68), p53, and p21. β-Actin was used as the loading control. **j,** RT-qPCR analysis of DDR- and cell-cycle-related genes, including *CDKN1A*, *GDF15*, *KIF20A*, and *CENPA*, in xenograft tumour tissues after in vivo GBA-007 treatment. Data are presented as mean ± SD. *P < 0.05, **P < 0.01, ***P < 0.001.

We first assessed biochemical inhibition of WRN helicase activity. GBA-007 potently inhibited WT WRN with an IC₅₀ of 102 nM, and retained strong inhibitory activity against WRN F730L with an IC₅₀ of 180 nM (Fig. 8b). To assess the RecQ family biochemical selectivity of GBA-007, we tested its inhibitory activity against BLM, RECQ1, RECQ4 and RECQ5. GBA-007 showed negligible inhibition of all four helicases with IC₅₀ values exceeding 10,000 nM (Supplementary Fig. S3b), demonstrating excellent intra-family target selectivity. Notably, GBA-007 showed potent antiproliferative activity against both parental HCT116 cells and HCT116_F730L cells, with IC₅₀ values of 145 nM and 261 nM, respectively (Fig. 8c). Extended profiling across a panel of additional cell lines demonstrated that GBA-007 exerted potent inhibitory activity against all tested MSI-H cell lines, with IC₅₀ values ranging from 56.7 nM to 93.4 nM in LS411N, LoVo and Ishikawa cells. In contrast, no meaningful inhibitory activity was observed against all tested MSS cell lines, with all IC₅₀ values exceeding 20,000 nM (Supplementary Fig. S3c–e), supporting the MSI-H-selective activity profile of GBA-007. To determine whether the potent inhibitory activity of GBA-007 against F730L-mutant HCT116 cells arises from the restoration of on-target pathway engagement, we evaluated WRN abundance, DDR activation and G_2_/M cell cycle arrest. GBA-007 induced dose-dependent downregulation of WRN in both parental and F730L-mutant cells and robustly upregulated DDR markers, including γ-H2AX, p-ATM, p-CHK2, p53 and p21 (Fig. 8d). Consistently, cell-cycle profiling confirmed pronounced G₂/M arrest in both cell types (Fig. 8e). Together, these biochemical and cellular data indicate that GBA-007 exhibits excellent target selectivity, overcomes F730L-associated resistance while maintaining on-target WRN inhibition phenotypes.

We next tested whether GBA-007 could overcome F730L-mediated resistance in vivo. A pharmacokinetic study revealed that GBA-007 exhibited favorable oral pharmacokinetic properties in BALB/c mice. After oral gavage at 10 mg/kg, GBA-007 was rapidly absorbed, with a Tₘₐₓ of 0.250 h, Cₘₐₓ of 2868 ng/mL, AUC_0-t_ of 3618 h ng/mL, T₁_/_₂ of 3.17 h, and an oral bioavailability of 106% (Fig. 8f). Oral administration of GBA-007 significantly suppressed tumour growth in both parental HCT116 and HCT116_F730L xenograft models (Fig. 8g), with no obvious body-weight loss during treatment (Fig. 8h). Western blotting and reverse transcription-quantitative polymerase chain reaction (RT-qPCR) analyses of tumour tissues showed reduced WRN protein levels and increased expression of γ-H2AX, p-CHK2, p53, and p21 in both parental and F730L tumours after treatment (Fig. 8i). In parallel, DDR-associated genes (*CDKN1A* and *GDF15*) were upregulated, whereas cell-cycle-associated genes (*KIF20A* and *CENPA*) were downregulated (Fig. 8j). These results indicate that GBA-007 induces WRN inhibition–associated molecular responses and antitumour activity in vivo in both WT and F730L settings.

Collectively, these findings demonstrate that a structure-guided, AI-enabled optimization workflow can rapidly convert a resistance bottleneck into a tractable design problem. Together with the alternating-therapy results, these data support a two-pronged resistance-management framework: exploit functional orthogonality through schedule and inhibitor selection where possible, and deploy mutation-aware design to address cross-resistant variants such as F730L.

## Discussion

As WRN inhibitors progress through early clinical development for MSI-H/dMMR tumours, acquired resistance is likely to be a major determinant of long-term benefit^28–30^. Recent preclinical studies have begun to define this landscape: independent efforts have shown that MSI cancer cells readily acquire on-target WRN alterations under WRN inhibitor pressure, with recurrent hotspot mutations and, in some settings, chemotype-selective resistance profiles that preserve sensitivity to alternative WRNi modalities^24–27^. In parallel, emerging structural work has clarified that clinically relevant WRNi can act by stabilizing distinct inactive, “off-DNA” conformations of the WRN helicase and by perturbing the conformational dynamics required for productive DNA engagement^27^. Together, these advances establish two key principles: first, on-target resistance to WRNi can emerge rapidly in hypermutable MSI backgrounds; second, mechanistically distinct WRNi can impose distinct selective pressures, creating the possibility that resistance to one modality may remain tractable to another^24–27^. Building on this foundation, our study advances WRNi resistance research in two translationally actionable directions. We first demonstrate that inhibitor-class–specific resistance landscapes can be leveraged prospectively to extend tumour control through a cyclic alternating regimen. We then use engineered F730L as a prospective cross-resistant bottleneck model and show that such bottlenecks can be addressed through a rapid, structure-guided and AI-enabled optimization workflow, providing a generalizable path to mutation-tolerant next-generation inhibitors.

Across independently derived resistance models in multiple MSI-H cell lines, we observed a convergent resistance phenotype characterized by a coordinated loss of WRNi pharmacodynamics: resistant cells failed to exhibit the canonical cascade of WRN protein reduction, DDR activation, G_2_/M cell cycle arrest and apoptotic commitment that is observed in parental cells upon WRN inhibition. Importantly, this resistant state was not explained by marked baseline WRN overexpression, but instead reflected failure of drug treatment to re-engage the on-target programme, consistent with impaired inhibitor engagement. Whole-exome sequencing further showed that resistance evolution converged strongly on WRN missense mutations despite the heterogeneous mutational background typical of MSI cells. Moreover, resistance trajectories were strikingly modality-specific: HRO761-resistant models predominantly selected mutations clustered at G729, F730 and I852, whereas VVD-214-resistant models selected mutations concentrated at E846. These patterns support the concept that non-covalent and covalent WRNi modalities apply distinct selective pressures on the WRN helicase, consistent with structural frameworks in which different inhibitors stabilize different inactive conformational states or depend on distinct local interaction networks within the helicase domain and its hinge region. In this regard, our data extend the emerging consensus from recent resistance-mapping studies by providing a direct, modality-matched comparison across clinically relevant WRNi classes and by linking mutation spectra to functional consequences^24–27^.

Critically, we moved beyond mutation cataloguing to establish causality and mechanism across biochemical and isogenic systems. Protein thermal shift assays indicated that resistance-associated mutations reduce inhibitor-induced stabilization of WRN, consistent with impaired binding. In vitro helicase assays showed that these mutations diminish inhibitor-mediated suppression of WRN activity, and isogenic knock-in models demonstrated sufficiency: introduction of single WRN point mutations into an otherwise matched HCT116 background was sufficient to confer marked resistance to the selecting inhibitor. These mutations also attenuated downstream pharmacodynamic outputs, including WRN protein reduction, DDR marker induction and G₂/M arrest. Together, these data establish a coherent causal chain from recurrent on-target sequence variants to impaired inhibitor engagement, reduced enzymatic inhibition, and failure to trigger the cytotoxic DDR activation and G_2_/M cell cycle arrest programme that underpins WRN–MSI synthetic lethality under pharmacologic perturbation^6,8^.

A central finding of this work is that inhibitor-class–specific resistance differences are not merely structural observations but can be operationalized as a functionally orthogonal resistance relationship. With the notable exception of F730L, most tested mutations that compromised the selecting inhibitor retained sensitivity to the alternate modality across biochemical inhibition, cellular antiproliferative activity, and re-induction of WRN inhibition-associated pharmacodynamic outputs. This cross-sensitivity was further supported in vivo using xenograft models carrying defined WRN point mutations, where inhibitor switching restored antitumour activity in many resistance contexts. In the broader field, recent preclinical efforts have suggested that certain WRN variants can preserve sensitivity to alternative WRNi, particularly when inhibitors have distinct binding modes or mechanisms^24–27^. Our study strengthens and extends this concept by defining orthogonality in functional terms—requiring not only viability effects but also restoration of on-target pharmacodynamics—and by demonstrating that such orthogonality holds across multiple experimental levels, including in vivo tumour response.

We then asked whether this orthogonality can be exploited proactively to delay the emergence of resistance and prolong tumour control, rather than being applied only reactively after resistance has been detected. In a long-term SW48 xenograft experiment, a 7-day cyclic alternating regimen between HRO761 and VVD-214 delayed tumour regrowth and reduced endpoint tumour burden compared with continuous monotherapy and a reduced-dose combination regimen. Notably, endpoint tumours in the alternating group retained pronounced WRN reduction and DDR activation, indicating sustained on-target engagement over prolonged treatment. These findings are consistent with evolutionary concepts in which temporal modulation of selective pressure can delay the rise of resistant clones in settings where multiple partially non-overlapping resistance trajectories exist^36,37^. While our data do not establish this 7-day cycle as optimal, they provide proof of principle that schedule design informed by functional orthogonality can extend tumour control beyond what is achievable with continuous single-agent exposure, representing a translational step beyond resistance mutation mapping.

The existence of functional orthogonality, however, also clarifies its limits. Our engineered F730L model underscores that switching and alternating strategies may be constrained by variants that compromise multiple modalities. Although the clinical frequency of F730L remains to be established in WRNi-treated patient samples, residue 730 lies within a recurrent resistance hotspot, making this mutation a useful stress test for prospective resistance management. In our models, F730L conferred pronounced resistance to both HRO761 and VVD-214, limiting the efficacy of modality switching and blunting the benefits of orthogonality-based scheduling. Importantly, our structural–dynamic analyses suggest that this bottleneck is not readily explained by the disruption of a single local contact. Instead, molecular dynamics simulations support a model in which F730L remodels the D1–D2 hinge-adjacent environment and destabilizes the local hydrophobic core, increasing conformational plasticity and reducing binding pose stability. For covalent inhibition modalities in particular, this destabilization can impair maintenance of reaction-competent geometries, providing a plausible mechanistic rationale for why closely related substitutions at the same residue (e.g., F730V versus F730L) can yield divergent cross-sensitivity behaviours. This structural–dynamic perspective complements recent structural insights into WRN conformational cycling and off-DNA inhibitor trapping by highlighting how hinge-region remodelling could propagate to the binding environment and create cross-resistance even between mechanistically distinct inhibitors^6,8,27^.

A major translational contribution of our study is that we did not treat this bottleneck model as a terminal failure mode. Instead, we demonstrate a rapid path from bottleneck identification to mutation-tolerant inhibitor design, and we argue that this approach represents a broader research paradigm for drug discovery against targets prone to rapid on-target resistance evolution. In hypermutable MSI-H/dMMR settings, resistance can arise on clinically relevant time scales, creating a mismatch between tumour evolution and conventional sequential drug development workflows in which next-generation inhibitors are often initiated only after resistance is observed clinically. Here, guided by our mechanism-based diagnosis of hinge-region remodelling, we reframed the design objective from maximizing WT potency to minimizing mutation-induced penalties across both WT and F730L contexts. Using a structure-guided, AI-enabled scoring strategy that jointly evaluated candidates in both conformational backgrounds and integrated synthesizability-aware filtering, we rapidly prioritized next-generation candidates and identified GBA-007 as a proof-of-concept lead compound. GBA-007 retained potent biochemical inhibition against F730L WRN, restored cellular pharmacodynamic responses and G_2_/M cell cycle arrest, and suppressed both WT and F730L xenograft growth without obvious toxicity signals in the tested setting. Conceptually, this establishes an integrated framework that combines (i) resistance mapping, (ii) mechanistic diagnosis of bottleneck variants, and (iii) rapid, mutation-aware design–make–test cycles. We propose that such a resistance-evolution–aware discovery workflow could be broadly applicable to drug discovery programmes in which on-target variants arise readily and bypass mechanisms are limited, and in which durable clinical benefit will likely require both schedule-level interventions and mutation-tolerant next-generation molecules.

Collectively, our findings support a stratified resistance-management framework. For resistance mutations that preserve sensitivity to an alternate WRNi modality, mutation-informed inhibitor selection and, potentially, orthogonality-informed cyclic alternating strategies may prolong tumour control. For engineered cross-resistant bottleneck models such as F730L, next-generation inhibitor development guided by structural and dynamic mechanisms may be required to restore tractability if analogous variants emerge clinically. While our work provides proof of concept for both arms of this framework, several limitations should be acknowledged. First, our conclusions are based primarily on cell lines and cell-derived xenograft (CDX) models; validation in patient-derived models and clinical resistance samples is needed to define the frequency, temporal emergence and clinical relevance of specific WRN resistance variants during patient treatment, including whether engineered bottleneck models such as F730L arise clinically. Second, although alternating therapy was associated with prolonged tumour control in our long-term model, the optimal cycle length, dosing intensity and generalizability across tumour contexts remain undefined, and will require pharmacokinetic/pharmacodynamic studies and longitudinal tracking of clonal dynamics. Third, GBA-007 remains a proof-of-concept candidate; its therapeutic window and activity across broader resistance backgrounds require further optimization. Finally, although our data underscore the prominence of on-target resistance in these models, clinical resistance may also be shaped by pharmacologic constraints, tumour-intrinsic adaptation, microenvironmental influences and non-genetic mechanisms, which should be incorporated into future resistance modelling.

Overall, this study advances WRNi resistance research from mutation identification toward resistance-informed therapeutic design and mutation-tolerant inhibitor discovery. By defining a functionally orthogonal resistance relationship between a clinically relevant non-covalent and covalent WRNi, translating this orthogonality into a feasible cyclic alternating regimen that prolongs tumour control, and providing a rapid AI-enabled route to overcome a cross-resistant bottleneck, our work offers an integrated conceptual and experimental framework to improve the durability of WRN-targeted therapies in MSI-H/dMMR cancers.

## Methods

### Chemistry

VVD-214 was purchased from Bide Pharmatech. HRO761 was synthesized following the method described previously^6^. GBA-007 was synthesized as described in the Supplementary Information. All compounds were verified for purity (>95%) by high-performance liquid chromatography (HPLC).

### Cell culture

HCT116, SW48, SW620, LS513, AGS, Caco-2, LoVo and Ishikawa cells were purchased from Procell Life Science & Technology Co., Ltd. LS411N cells were purchased from Nanjing Cobioer Biosciences Co., Ltd. HCT116 cells were cultured in McCoy’s 5A medium (Meilunbio, MA0314); Caco-2 cells were cultured in MEM with NEAA (Meilunbio, MA0217); LS411N and LS513 cells were cultured in RPMI-1640 medium (Meilunbio, MA0215); LoVo cells were cultured in Ham’s F-12K medium (Meilunbio, MA0230); AGS cells were cultured in Ham’s F-12 medium (Meilunbio, MA0229); Ishikawa cells were cultured in high-glucose DMEM (Meilunbio, MA0212); and SW48 and SW620 cells were cultured in Leibovitz’s L-15 medium (Meilunbio, MA0547). Except for SW48 and SW620, which were maintained at 37 °C without CO₂ supplementation, all other cell lines were maintained at 37 °C in a humidified incubator containing 5% CO₂. All media were supplemented with 10% fetal bovine serum (Meilunbio, PWL002) and 1% penicillin–streptomycin solution (Meilunbio, PWL062), except that Caco-2 cells were cultured in MEM with NEAA supplemented with 20% fetal bovine serum and 1% penicillin–streptomycin solution. Cell lines were authenticated by short tandem repeat profiling and were routinely tested to be negative for mycoplasma contamination.

### Generation of WRN inhibitor-resistant cell populations

WRN inhibitor-resistant cell populations were established by cyclic drug exposure. Parental HCT116, SW48 and Ishikawa cells were treated with 1 μM HRO761 or VVD-214 for 24 h, followed by exposure to 0.1 μM of the same inhibitor for 72 h. This treatment cycle was repeated for 1–2 months until cells maintained stable growth under drug selection. After the selection period, resistant cell populations were expanded and maintained under the corresponding inhibitor. Before subsequent experiments, resistant cells were cultured for at least three passages in drug-free medium, and the stability of the resistant phenotype was confirmed by determining the RI.

### Clonogenic assay

Cells were seeded in 6-well plates at 200–1,000 cells per well, with the exact seeding density optimized for each cell line and allowed to adhere overnight. Cells were then treated with DMSO, HRO761 or VVD-214 for 5 days. After treatment, colonies were washed with PBS, fixed with methanol and stained with Crystal Violet Staining Solution (Beyotime, C0121). Plates were then washed with PBS, air-dried and imaged.

### Cell proliferation assay

Cells were seeded in 96-well plates (500–1,000 cells/well), allowed to adhere overnight, and treated with serial dilutions of compounds or DMSO for 96 h. Proliferation was assessed using the CellTiter-Glo (CTG) Luminescent Cell Viability Assay (Meilunbio, PWL214), and luminescence was normalized to DMSO controls. Dose–response curves were fitted with a four-parameter logistic model in GraphPad Prism 9.5.1, and IC₅₀ values were calculated.

### Western blot analysis

Cells were washed with cold PBS and lysed in RIPA buffer (Meilunbio, MA0152) containing protease and phosphatase inhibitors (Meilunbio, MB2678 and MB12707). Xenograft tumour tissues were homogenized in the same buffer. Lysates were centrifuged at 12,000 × g for 15 min at 4 °C, and supernatants were collected. Protein concentrations were measured using the Pierce™ BCA Protein Assay Kit (Thermo Fisher Scientific, 23227). Samples were separated by SDS-PAGE, transferred to nitrocellulose membranes, blocked with 5% non-fat milk in TBST, and incubated overnight at 4 °C with primary antibodies against WRN (CST, 4666S), phospho-ATM (Ser1981) (CST, 5883S), phospho-CHK2 (Thr68) (CST, 2197S), γ-H2AX (Ser139) (CST, 9718S), p53 (CST, 9282S), p21 (CST, 2947S), and β-Actin (CST, 4970S). After TBST washes, membranes were incubated with HRP-conjugated secondary antibodies (Promega, W4011 and W4021), washed again, and bands visualized using chemiluminescent substrate (Meilunbio, MA0186) on a Syngene GeneGnome XRQ. Band intensities were quantified with ImageJ.

### Flow cytometric cell-cycle assay

Cells were treated with the indicated compounds for 24 h. After treatment, cells were harvested using Trypsin-EDTA Solution (Meilunbio, MA0302), washed with cold PBS and fixed in 75% ethanol at 4 °C overnight. Fixed cells were washed with PBS and stained with propidium iodide/RNase A staining solution from the Cell Cycle and Apoptosis Analysis Kit (Beyotime, C1052) at 37 °C for 30 min in the dark. DNA content was analyzed using an Agilent NovoCyte flow cytometer, and the proportions of cells in G₀/G₁, S and G₂/M phases were quantified using NovoExpress software. The gating strategy is shown in Supplementary Fig. S4a.

### Flow cytometric apoptosis assay

Cells were treated with the indicated compounds for 24 h. After treatment, cells were harvested using Trypsin Solution without EDTA (Meilunbio, MA0234), washed with cold PBS and stained using the Annexin V-FITC Cell Apoptosis Detection Kit (Beyotime, C1062) according to the manufacturer’s instructions. Stained cells were analyzed using an Agilent NovoCyte flow cytometer, and apoptosis was quantified using NovoExpress software. Late apoptotic cells were defined as Annexin V-positive and PI-positive cells. The gating strategy is shown in Supplementary Fig. S4b.

### Whole-exome sequencing and variant analysis

Whole-exome sequencing was performed by Tsingke Biotechnology Co., Ltd. Genomic DNA was extracted from parental and WRN inhibitor-resistant cells, and DNA quality and quantity were assessed prior to library preparation. DNA was fragmented, end-repaired, A-tailed, ligated to sequencing adapters, and amplified by polymerase chain reaction (PCR). Exonic regions were enriched using biotin-labelled DNA probes and streptavidin magnetic beads, amplified, and subjected to library quality control. Sequencing was performed on a DNBSEQ platform with paired-end 150 bp reads.Raw reads were processed with fastp to remove adapters and low-quality bases (<35 bp reads, >5 undetermined bases, or >40% bases with Q ≤ 15). Clean reads were aligned to GRCh37 using Sentieon and SAMtools, with duplicate marking and base quality score recalibration. Somatic single-nucleotide variants (SNVs) and Indels were called with Genome Analysis Toolkit (GATK) Mutect2 using parental cells as matched controls and annotated with Variant Effect Predictor (VEP). Coding variants were extracted from resistant populations, and recurrently mutated genes were ranked by mutation frequency.

### Expression and purification of wild-type and mutant human WRN (D1D2RH)

cDNAs encoding wild-type and mutant human WRN (D1D2RH; residues Asn517–Pro1238) were cloned into pFastBac1 with a C-terminal 8×His tag and PreScission cleavage site. Recombinant baculoviruses were generated in Sf9 cells using Cellfectin™ II (Thermo Fisher Scientific, 10362100) and used to infect High Five cells. Seventy-two hours post-infection, cells were harvested and lysed in 50 mM Tris-HCl, pH 8.0, 300 mM NaCl, 20 mM imidazole, 1 mM TCEP, 10% glycerol, and protease inhibitors (Meilunbio, MB2678) by sonication. Lysates were clarified by centrifugation and loaded onto a HisTrap HP column (Cytiva, 17524801). Bound proteins were eluted with an imidazole gradient and further purified by size-exclusion chromatography on a Superdex 200 Increase 10/300 GL column (Cytiva, 28990944) equilibrated with 50 mM Tris-HCl, pH 8.0, 150 mM NaCl, 1 mM TCEP, 10% glycerol. Purified proteins were analyzed by SDS-PAGE, concentrated, aliquoted, flash-frozen in liquid nitrogen, and stored at −80 °C.

### Protein thermal shift assay

Purified wild-type and mutant human WRN (D1D2RH) proteins were diluted to 1 μM in assay buffer (20 mM Tris-HCl, pH 8.0, 100 mM NaCl, 1 mM TCEP) and incubated with HRO761, VVD-214, or DMSO control. SYPRO Orange dye (Sigma-Aldrich, S5692) was added, and protein thermal shift assays were performed on a Bio-Rad real-time PCR system. The temperature was ramped from 25 °C to 90 °C in 0.5 °C increments, with fluorescence recorded after a 10 s hold at each step. Melting temperatures (T_m_) were calculated from fluorescence curves, and ΔT_m_ values determined relative to DMSO controls.

### WRN helicase activity assay

WRN helicase activity was measured using a fluorescence-quenching DNA unwinding assay. The fluorescent DNA substrate was prepared by annealing equimolar OLIGOA-BHQ2 and OLIGOB-TAMRA oligonucleotides (see Supplementary Table S4) in 50 mM NaCl by heating to 95 °C for 5 min and slow cooling to 4 °C. Reactions were performed in assay buffer (30 mM Tris-HCl, pH 7.5, 0.5 mM MgCl₂, 50 mM NaCl, 0.02% BSA, 1 mM TCEP) in 384-well black plates (Corning, 3575). Compounds or DMSO control (1 µL) were added to wells, followed by 10 µL of 100 nM WRN protein with 200 nM trap ssDNA (plus 0.2 mM ATP for VVD-214 assays) and incubation for 30 min. Reactions were initiated by adding 10 µL of 200 nM fluorescent DNA substrate and 6 mM ATP. Fluorescence was monitored kinetically on an EnVision plate reader (PerkinElmer) at 544 nm/590 nm. Helicase activity was normalized to DMSO control, and IC₅₀ values were determined from nonlinear dose–response curves in GraphPad Prism 9.5.1.

### WRN ATPase activity assay

WRN ATPase activity was measured using the ADP-Glo kinase assay kit (Promega, V9102). WRN and serially diluted compounds were prepared in assay buffer containing 25 mM Tris-HCl pH 8.0, 5 mM NaCl, 2 mM MgCl₂, 1 mM DTT, 0.01% Tween-20 and 2.5 μg/mL calf thymus DNA, and incubated in 384-well plates (Corning, 3575) at room temperature for 15 min. The reaction was initiated by adding DNA substrate and ATP, with final concentrations of 200 nM WRN, 300 nM DNA substrate and 3 mM ATP, followed by incubation at room temperature with shaking for 3 h. Reactions were diluted 5000-fold with assay buffer, and 10 μL of the diluted mixture was transferred to white 384-well plates (PerkinElmer, 6007299). ADP-Glo reagent and kinase detection reagent were sequentially added with 40 min incubation after each addition. Luminescence was measured using an EnVision plate reader. Percent inhibition was calculated relative to the DMSO control, and IC₅₀ values were determined by fitting dose–response curves using a four-parameter logistic model in GraphPad Prism 9.5.1.

### BLM, RECQ1, RECQ4 and RECQ5 ATPase activity assay

The ATPase activities of BLM, RECQ1, RECQ4 and RECQ5 were measured by ICE Bioscience Co., Ltd. using an ADP-Glo-based enzymatic activity assay. Briefly, test compounds were first prepared in DMSO and serially diluted at a 1:3 ratio in a 384-well dilution plate. Then, 0.10 μL of the diluted compound solution was transferred into each well of a 384-well assay plate. Subsequently, 5 μL of enzyme working solution containing BLM, RECQ1, RECQ4 or RECQ5 was added to the corresponding wells and incubated at 25 °C for 10 min. The enzymatic reaction was initiated by adding 5 μL of substrate working solution, followed by incubation at 25 °C for 60 min. After the reaction, 5 μL of ADP-Glo reagent was added to terminate the enzymatic reaction and deplete the remaining ATP, and the plate was incubated at 25 °C for 40 min. Next, 10 μL of detection solution was added to convert ADP into ATP and generate a luminescent signal, followed by another incubation at 25 °C for 40 min. Luminescence signals were recorded using a multimode plate reader. The percentage inhibition was calculated relative to DMSO-treated control wells, and IC_50_ values were determined by nonlinear regression using a four-parameter logistic model.

### Generation of isogenic WRN point-mutant cells

Isogenic WRN point-mutant HCT116 cells were generated by CRISPR/Cas9-mediated homology-directed repair. sgRNAs targeting G729/F730, E846, and I852 regions were cloned into pSpCas9(BB)-2A-Puro (PX459). Parental HCT116 cells were co-transfected with PX459-sgRNA plasmids and mutation-specific donor templates using PolyJet™ DNA In Vitro Transfection Reagent (SignaGen, SL100688). Single-cell clones were isolated by limiting dilution, expanded, and genomic DNA was extracted for PCR amplification and Sanger sequencing of target regions. sgRNA and donor template sequences are provided in the Supplementary Table S5.

### Pharmacokinetic study

Female BALB/c mice aged 8-9 weeks and weighing approximately 20 g were used for pharmacokinetic analysis with approval from the Institutional Animal Care and Use Committee at the Shanghai Institute of Materia Medica, Chinese Academy of Sciences (IACUC No. 2026-04-ZMY-18). GBA-007 was administered intravenously at 1 mg/kg or orally by gavage at 10 mg/kg, with three mice per route. The intravenous formulation consisted of 5% DMSO, 20% PEG300 and 75% saline and was dosed at 5 mL/kg. The oral formulation consisted of 5% DMSO, 5% PEG400 and 90% 10% HP-β-CD in water and was dosed at 10 mL/kg. Plasma concentrations of GBA-007 were measured after dosing, and pharmacokinetic parameters were calculated by non-compartmental analysis. Oral bioavailability was calculated from dose-normalized AUC_0-t_ values after oral and intravenous administration.

### In vivo efficacy studies

Female BALB/c nude mice (6–8 weeks) were used with approval from the Institutional Animal Care and Use Committee at the Shanghai Institute of Materia Medica, Chinese Academy of Sciences (IACUC No. 2025-04-ZMY-11). HRO761, VVD-214, and GBA-007 were formulated in an aqueous vehicle containing 10% N-methyl-2-pyrrolidone, 15% Kolliphor EL, 1% Tween-20, and 15% hydroxypropyl-β-cyclodextrin, with sterile water as the balance, and administered once daily by oral gavage.

For short-term studies, mice were subcutaneously inoculated with 5 × 10⁶ parental HCT116 or isogenic WRN point-mutant HCT116 cells. When tumours reached 100–200 mm³, mice were randomized to vehicle or treatment groups (n = 6 per group) and treated with vehicle, HRO761 (60 mg/kg, qd, p.o.), VVD-214 (20 mg/kg, qd, p.o.), or GBA-007 (60 mg/kg, qd, p.o.). Tumour volume and body weight were measured every 3 days, and tumour volume calculated as length × width² / 2.

For the long-term SW48 xenograft study, mice were inoculated with 5 × 10⁶ SW48 cells and, at 100–200 mm³ tumour volume, randomized to vehicle or treatment groups (n = 6 per group) including HRO761 monotherapy (60 mg/kg, qd, p.o.), VVD-214 monotherapy (20 mg/kg, qd, p.o.), combination (HRO761 30 mg/kg + VVD-214 10 mg/kg, qd, p.o.), or 7-day cyclic alternating treatment (HRO761 60 mg/kg and VVD-214 20 mg/kg in alternating 7-day cycles). Tumour volume was monitored every 4 days.

### RT-qPCR assay

Total RNA was extracted from ∼50 mg PBS-washed tumour tissue using RNA extraction reagent (Vazyme, R701-02). Tissue was lysed in 600 µL reagent, mechanically disrupted with grinding beads (Servicebio, G0203-150G), and centrifuged at 12,700 rpm for 15 min. Supernatants were mixed with 200 µL RNase-free H₂O, incubated 5 min at room temperature, and centrifuged again. RNA was precipitated with isopropanol, washed twice with 75% ethanol, air-dried, and dissolved in 20–100 µL RNase-free H₂O. RNA concentration and purity were measured with a NanoDrop spectrophotometer. Reverse transcription was performed using HiScript II Q RT SuperMix (Vazyme, R233-01), and RT-qPCR was carried out with ChamQ SYBR RT-qPCR Master Mix (Vazyme, Q331-03) on a QuantStudio 5 Real-Time PCR System (Thermo Fisher Scientific). Relative gene expression was calculated using the 2^−ΔΔCt method and normalized to *ACTB*. Primer sequences are provided in the Supplementary Table S6.

### Protein Preparation and System Setup for Molecular Dynamics Simulations

To investigate the structural basis underlying the differential sensitivity of the F730V and F730L mutants to VVD-214, molecular dynamics (MD) simulations were performed for VVD-214 in complex with the WRN helicase domain in the WT, F730V, and F730L backgrounds. Protein structures were prepared and optimized using the Protein Preparation Wizard implemented in Schrödinger Maestro version 13.5.128 (Release 2023-1). Protein protonation states were checked and assigned under physiologically relevant conditions, and local hydrogen-bonding networks and side-chain orientations were optimized during preparation.

The resulting WT and mutant protein–ligand complexes were subsequently used to build MD simulation systems. The protein was described using the CHARMM36 force field, whereas the ligand VVD-214 was parameterized using the CGenFF force field. Each system was solvated in explicit TIP3P water and neutralized with NaCl ions. Standard energy minimization and equilibration procedures were applied prior to the production phase.

### Molecular Dynamics Simulations and Trajectory Analysis

All MD simulations were performed using GROMACS 2026.1. For each of the WT, F730V, and F730L complex systems, two independent 1-μs production simulations were carried out using different random velocity seeds to improve sampling robustness. Production simulations were performed with a 2-fs integration time step at 303 K under periodic boundary conditions.

Trajectory analyses were conducted to characterize both protein- and ligand-level dynamic behavior. Protein conformational stability and flexibility were evaluated by calculating the RMSD, RMSF and Rg. Ligand conformational stability was assessed by ligand RMSD analysis. To monitor the geometry relevant to covalent engagement, the distance between the sulfur atom of Cys727 and the β-carbon of the acrylamide warhead of VVD-214 was measured throughout the simulations. Representative conformations were extracted from the trajectories for comparative structural analysis and visualization.

### Generation and Prioritization of Candidate Resistance-Overcoming Analogues

To develop second-generation HRO761 analogues capable of overcoming F730L-mediated resistance, focused analogue generation was performed using the Custom R-group Enumeration module in the Schrödinger suite. Different ring systems and substituents were introduced at the designated optimization sites to construct a chemically tractable virtual library. All generated molecules were subjected to synthesizability filtering, and only compounds with SA score < 4 and RAscore > 0.7 were retained for subsequent evaluation. For structure-based prioritization, molecular docking was performed based on the crystal structure of the WRN–HRO761 complex (PDB ID: 8PFO). Candidate molecules were docked into both the WT and F730L WRN structures using the Schrödinger docking module in SP precision, with ligand sampling set to refine only, so as to preserve the core binding mode of the reference scaffold while evaluating local structural modifications.

For PBCNet2.0 evaluation, the co-crystallized HRO761 in the WRN complex was used as the reference ligand, and the docked poses of the newly generated analogues in the corresponding WT and F730L complex structures were used as the query input structures. Candidate prioritization was not based solely on predicted affinity in a single protein background. Instead, compounds were selected by jointly considering their predicted binding performance in both WT and F730L, with emphasis on mutation tolerance, preservation of the core binding pose, and overall cross-system compatibility. Based on this integrated evaluation, five compounds with favorable predicted binding behavior in both the WT and mutant systems were selected for synthesis and experimental validation.

### Statistical analysis

Statistical analyses were performed using GraphPad Prism 9.5.1. Data are presented as mean ± SD. Dose– response curves were fitted by nonlinear regression using a four-parameter logistic model to calculate IC_50_ values. Comparisons between two independent groups were performed using unpaired two-tailed Student’s t-tests. Comparisons among three or more groups were performed using one-way ANOVA followed by Dunnett’s multiple-comparisons test. For CDX tumour studies, statistical analyses were performed on endpoint tumour volumes. Two-group comparisons were analysed using unpaired two-tailed Student’s t-tests, whereas multiple-group comparisons against a single control group were analysed using one-way ANOVA followed by Dunnett’s multiple-comparisons test. Statistical significance was defined as follows: ns, P > 0.05; *P < 0.05; **P < 0.01; and ***P < 0.001.

## Acknowledgments

We thank the staff members of the Large-scale Protein Preparation System (https://cstr.cn/31129.02.NFPS.LSPS) at the National Facility for Protein Science in Shanghai (https://cstr.cn/31129.02.NFPS), for providing technical support and assistance in data collection and analysis.

## Author contributions

Conceptualization: G.Z., D.T., X.L., K.C., X.Lu., M.W., S.Z., and M.Z.; Methodology: G.Z., D.T., M.P., D.W., Z.F., Y.T., C.S., X.Y., M.H., P.S., Y.L., Z.X., R.C., Y.Z., Z.G., S.C., X.M., J.X., Y.C., Z.Fa., Q.S., R.Y., and C.Q.; Investigation: G.Z., D.T., M.P., X.L., K.C., X.Lu., M.W., S.Z., and M.Z.; Visualization: G.Z., D.T., and M.P.; Funding acquisition: S.Z. and M.Z.; Writing - original draft: G.Z., D.T., and M.P.; Writing - review & editing: G.Z., D.T., M.P., S.Z., and M.Z..

## Data availability

Source data are provided with this paper. Whole-exome sequencing data have been deposited in the NCBI Sequence Read Archive (SRA) under BioProject accession PRJNA1474530. Molecular dynamics input files, representative structures and processed analysis data supporting the MD analyses have been deposited in Zenodo under accession https://doi.org/10.5281/zenodo.20505169. PBCNet2.0 screening-library information and dual-background ranking results for scaffold and analogue prioritization are provided in Supplementary Tables S2 and S3. All other data supporting the findings of this study are available within the Article and its Supplementary Information or from the corresponding authors upon reasonable request.

## Code availability

No custom code central to the conclusions of this study was generated. PBCNet2.0 is a previously published open-source model and is available at https://github.com/YuJie-0202/PBCNet2.0. Molecular dynamics simulations, docking, scoring and data analyses were performed using publicly available or commercial software as described in the Methods.

## Supplementary information

### Supplementary Information

Supplementary Information file contains Supplementary Figs. S1–S4; Supplementary Tables S1 and S4–S6; Supplementary Chemistry Methods; Supplementary Schemes S1–S3; Synthetic procedures and characterization data for GBA-001, GBA-006, GBA-007, GBA-008, GBA-029; ^1^H NMR, ^13^C NMR, ^19^F NMR, HRMS and HPLC spectra of GBA-001, GBA-006, GBA-007, GBA-008, GBA-029; and uncropped western blot images.

### Supplementary Data

Supplementary Data file contains Supplementary Tables S2 and S3.

## Reporting Summary

**Code and Software Submission Checklist.**

## Source data

Source Data file contains all source data excluding those provided in the Supplementary Information file, Supplementary Data file, and datasets deposited in public databases.

