## Supplementary material for "Functional orthogonality of WRN inhibitor resistance enables alternating therapy and mutation-tolerant inhibitor design in MSI-H cancers": Supplementary Information_20260608.docx

**This Supplementary Information includes:**

Supplementary Figs. S1–S4

Supplementary Tables S1 and S4–S6

Supplementary Chemistry Methods

Supplementary Schemes S1–S3

Synthetic procedures and characterization data for GBA-001, GBA-006, GBA-007, GBA-008, GBA-029

^1^H NMR, ^13^C NMR, ^19^F NMR, HRMS and HPLC spectra of GBA-001, GBA-006, GBA-007, GBA-008, GBA-029

Uncropped western blot images

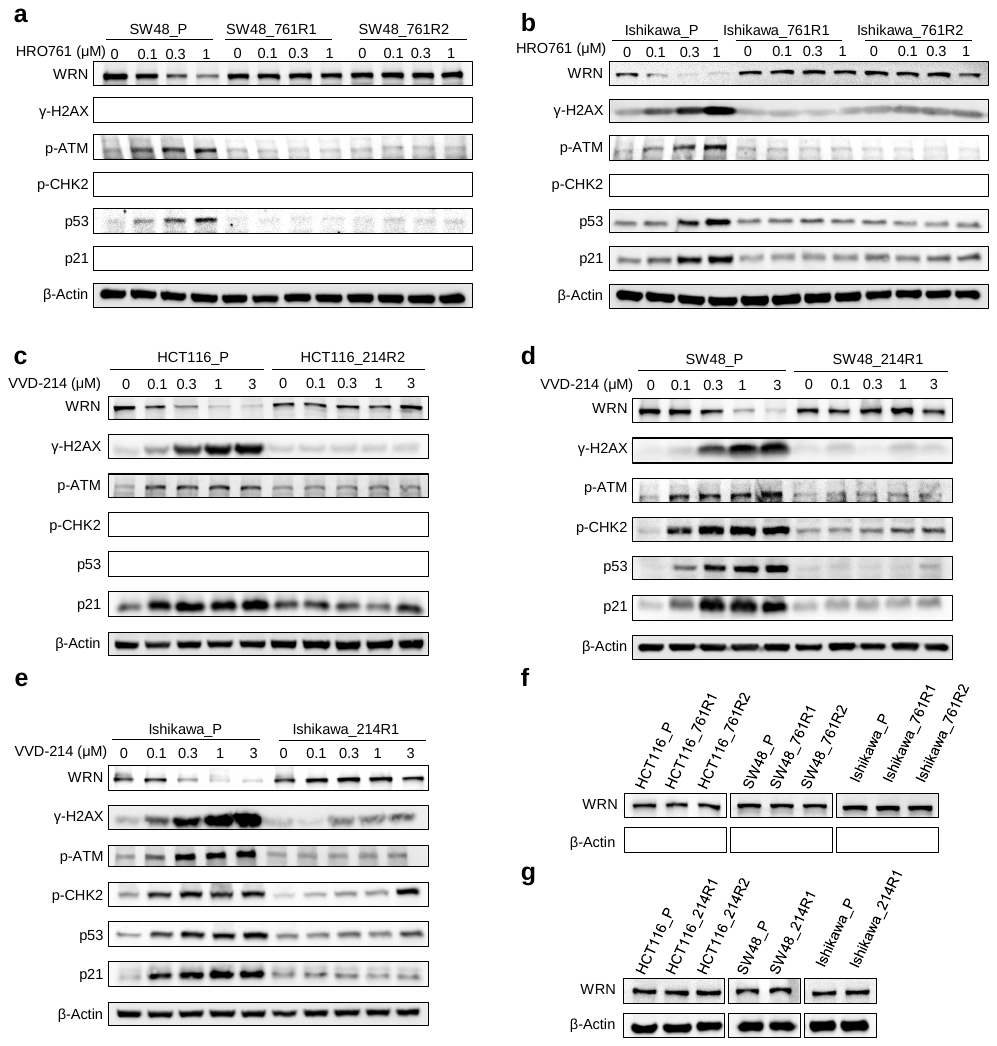

**Supplementary Fig. S1. Western blot analysis of WRN and DNA damage response-related proteins in parental and WRN inhibitor-resistant cells.**

**a–e,** Western blot analysis of WRN, γ-H2AX and DDR pathway-related proteins, including components of the ATM–CHK2 and p53–p21 pathways, in parental cells and the corresponding WRN inhibitor-resistant cells after treatment with HRO761 or VVD-214 for 24 h. **f, g,** Western blot analysis of basal WRN protein levels in parental cells and the corresponding WRN inhibitor-resistant cells under untreated conditions.

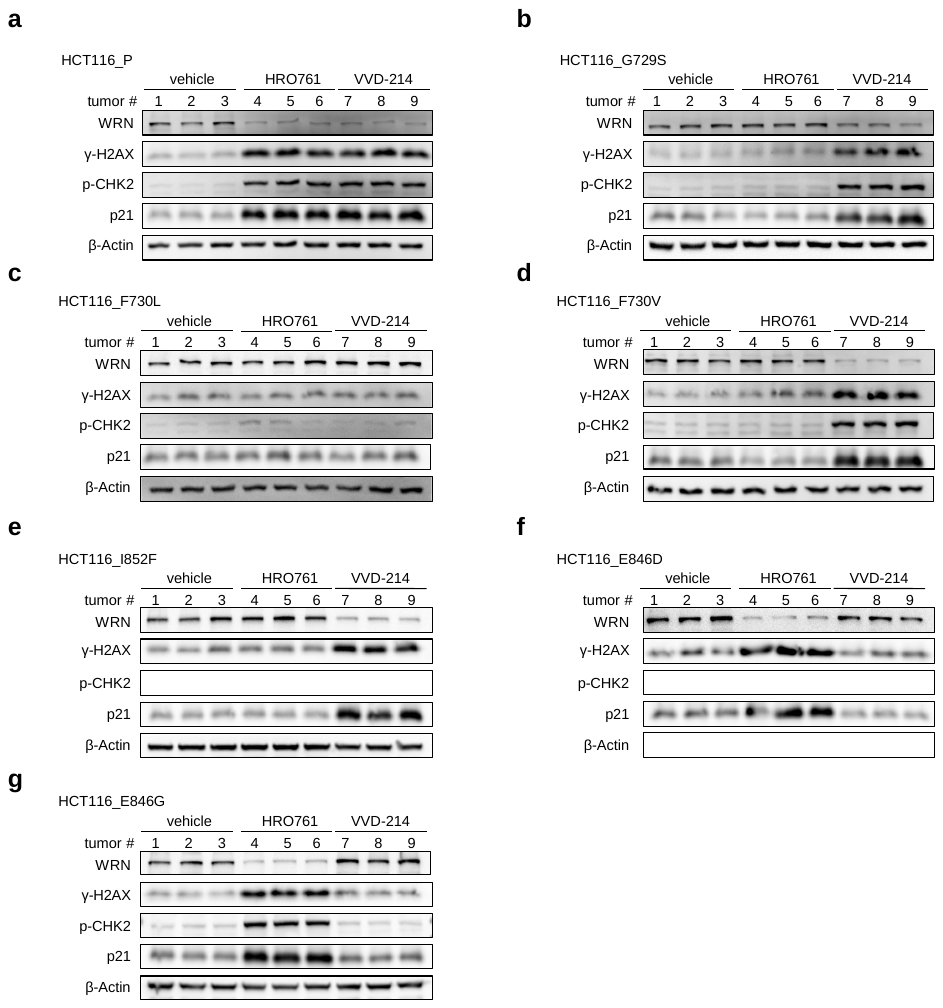

**Supplementary Fig. S2. Western blot analysis of WRN and DDR-related proteins in cell-derived xenograft (CDX) tumour tissues after WRN inhibitor treatment.**

**a–g,** Western blot analysis of WRN, γ-H2AX and DDR pathway-related proteins, including p-CHK2 and p21, in HCT116 CDX tumour tissues derived from cells expressing wild-type (WT) WRN or carrying the indicated WRN point mutations (G729S, F730L, F730V, I852F, E846D and E846G) after treatment with HRO761 or VVD-214.

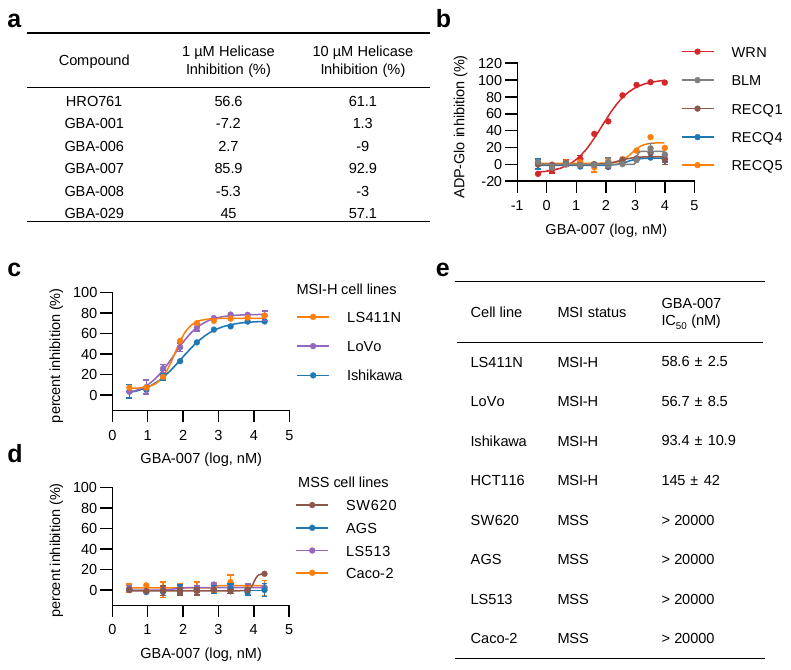

**Supplementary Fig. S3. Biochemical selectivity and cellular sensitivity profile of GBA-007.**

**a,** Helicase activity inhibition assays assessing the inhibitory effects of HRO761 and five candidate compounds on F730L-mutant WRN. **b,** ADP-Glo-based ATPase activity assays assessing the inhibitory effects of GBA-007 on WRN, BLM, RECQ1, RECQ4 and RECQ5. GBA-007 showed weak or negligible inhibition against BLM, RECQ1, RECQ4 and RECQ5, with IC₅₀ values greater than 10,000 nM. **c–e, Dose–response analysis of GBA-007 in MSI-H and MSS cell lines. c, Dose–response curves showing the effects of GBA-007 on cell proliferation in MSI-H cell lines, including LS411N, LoVo and Ishikawa. d, Dose–response curves showing the effects of GBA-007 on cell proliferation in** microsatellite-stable (MSS) cell lines **cell lines, including SW620, AGS, LS513 and Caco-2. e, IC₅₀ values of GBA-007 grouped by MSI status, comparing the sensitivity of MSI-H and MSS cell lines. Experiments were independently repeated three times.** Data are presented as mean ± standard deviation (SD).

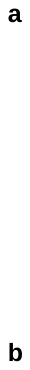

**Supplementary Fig. S4. Representative gating strategy for flow-cytometry analysis.**

**a,** Representative gating strategy for cell-cycle analysis. Cell debris was excluded based on FSC-A and SSC-A, and single cells were gated using FSC-A and FSC-H. G0/G1, S and G_2_/M populations were assigned based on PI DNA-content profiles. **b,** Representative gating strategy for apoptosis analysis. Cell debris was excluded based on FSC-A and SSC-A, and single cells were gated using FSC-A and FSC-H. Apoptotic populations were analyzed by Annexin V-FITC and PI staining. Late apoptotic cells were defined as Annexin V-positive and PI-positive cells.

**Supplementary Table S1. Dual-background ranking of scaffold and analogue candidates by PBCNet2.0-derived affinity scores and Glide SP docking scores.**

| **Candidate class** | **Candidate ID** | **Chemical structure** | **PBCNet2.0 dual-background balanced rank** | **Glide SP dual-background balanced rank** |
| --- | --- | --- | --- | --- |
| Scaffold | Scaffold 1 | 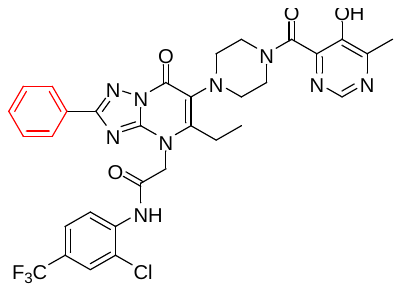 | 2/44 | 2/44 |
|  | Scaffold 2 | 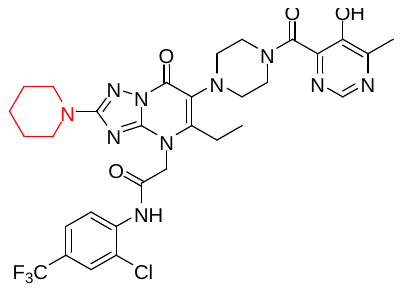 | 6/44 | 16/44 |
|  | Scaffold 3 | 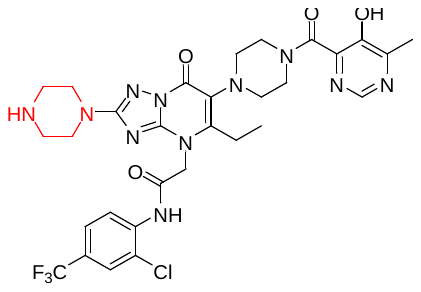 | 7/44 | 18/44 |
|  | Scaffold 4 | 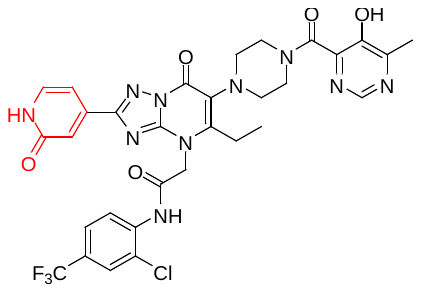 | 8/44 | 1/44 |
| Analogue | GBA-006 | 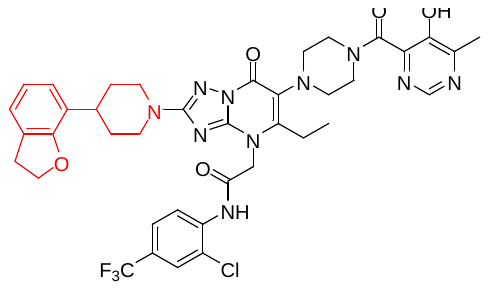 | 1/103 | 6/103 |
|  | GBA-008 | 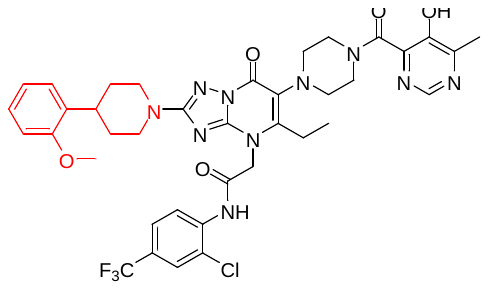 | 4/103 | 7/103 |
|  | GBA-029 | 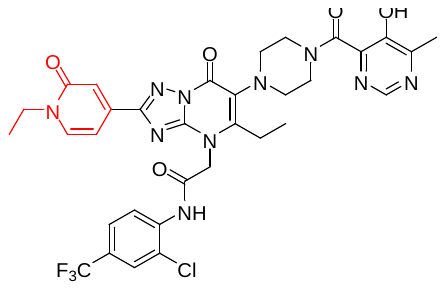 | 5/103 | 37/103 |
|  | GBA-007 | 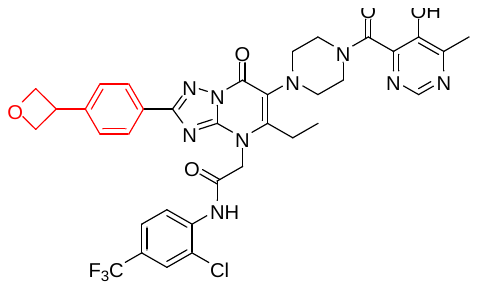 | 8/103 | 21/103 |
|  | GBA-001 | 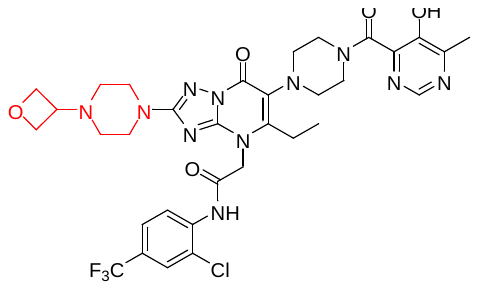 | 27/103 | 102/103 |

Note: Candidate scaffolds and analogues were ranked using two dual-background scoring schemes based on either PBCNet2.0-derived predicted affinity scores or Glide SP docking scores in WT and F730L WRN structures. For each scoring scheme, scores in the two protein backgrounds were first normalized to a 0–1 scale, with 1 corresponding to the most favorable score. For PBCNet2.0-derived affinity scores, higher values were treated as more favorable; for Glide docking scores, more negative values were treated as more favorable. A balanced dual-background score was then calculated as: balanced score = 0.60 × min (SWT, SF730L) + 0.40 × mean (SWT, SF730L), where SWT and SF730L are the normalized scores in the WT and F730L systems, respectively. This ranking strategy prioritizes candidates that maintain favorable predicted performance in both protein backgrounds and penalizes candidates that perform well in only one system. Rank values are shown as x/44 for scaffold candidates and x/103 for analogue candidates, where smaller x indicates higher priority. The analogue ranking was performed on 103 newly generated analogues [after excluding the reference compound/invalid entry; revise as appropriate]. This analysis compares the prioritization behavior of PBCNet2.0-derived affinity scoring and Glide docking-score ranking and was used to support mutation-aware candidate selection.

**Supplementary Table S4. Oligonucleotide sequences used in the WRN helicase assay**

| **Name** | **Sequence (5′→3′)** |
| --- | --- |
| OLIGOA-BHQ2 | TTTTTTTTTTTTTTTTTTTTTTTTTTTTTTCGTACCCGATGTGTTCGTTC-BHQ2 |
| OLIGOB-TAMRA | TAMRA-GAACGAACACATCGGGTACGTTTTTTTTTTTTTTTTTTTTTTTTTTTTTT |
| Trap ssDNA | GCACTGGCCGTCGTTTTACG |

**Supplementary Table S5. sgRNA and donor template sequences used for generation of isogenic WRN point-mutant HCT116 cells**

| **Category** | **Name** | **Sequence (5′→3′)** |
| --- | --- | --- |
| sgRNA | G729/F730 sgRNA | TGGTTCGATCAAAACCAGTAC |
| sgRNA | E846 sgRNA | TTACGGTGCTCCTAAGGACA |
| sgRNA | I852 sgRNA | GAATCATATTATCAGGAGAT |
| Donor template | G729S | GACATTGTACGTTGCTTAAATCTGAGAAATCCTCAGATCACCTGTACTAGCTTTGATCGACCAAACCTGTATTTAGAAGTTAGGCGAAAAACAGGGAA |
| Donor template | F730L | GACATTGTACGTTGCTTAAATCTGAGAAATCCTCAGATCACCTGTACTGGTCTGGATCGACCAAACCTGTATTTAGAAGTTAGGCGAAAAACAGGGAA |
| Donor template | F730V | GACATTGTACGTTGCTTAAATCTGAGAAATCCTCAGATCACCTGTACTGGTGTGGATCGACCAAACCTGTATTTAGAAGTTAGGCGAAAAACAGGGAA |
| Donor template | E846D | AAGCTGACATTCGCCAAGTCATTCATTACGGTGCTCCTAAGGACATGGACTCATATTATCAGGAGATTGGTAGAGCTGGTCGTGATGGACTTCAAAGTTC |
| Donor template | E846G | AAGCTGACATTCGCCAAGTCATTCATTACGGTGCTCCTAAGGACATGGGCTCATATTATCAGGAGATTGGTAGAGCTGGTCGTGATGGACTTCAAAGTTC |
| Donor template | I852F | AGTCATTCATTACGGTGCTCCTAAGGACATGGAATCATATTATCAGGAATTTGGTAGAGCTGGTCGTGATGGACTTCAAAGTTCTTGTCACGTCCTCTGG |

**Supplementary Table S6. Primer sequences used for quantitative real-time PCR**

| **Target** | **Primer** | **Sequence (5′→3′)** |
| --- | --- | --- |
| *ACTB* | Forward | CATGTACGTTGCTATCCAGGC |
| *ACTB* | Reverse | CTCCTTAATGTCACGCACGAT |
| *CDKN1A* | Forward | TGTCCGTCAGAACCCATGC |
| *CDKN1A* | Reverse | AAAGTCGAAGTTCCATCGCTC |
| *GDF15* | Forward | GACCCTCAGAGTTGCACTCC |
| *GDF15* | Reverse | GCCTGGTTAGCAGGTCCTC |
| *CENPA* | Forward | GACGCCTATCTCCTCACCTTA |
| *CENPA* | Reverse | GTTGCACATCCTTTGGGAAGA |
| *KIF20A* | Forward | TGCTGTCCGATGACGATGTC |
| *KIF20A* | Reverse | AGGTTCTTGCGTACCACAGAC |

**Chemistry.** General information: Starting materials, reagents and solvents were purchased from Bide Pharmatech, Adamas-beta, Energy Chemical, and J&K, and were used without further purification. ^1^H Nuclear magnetic resonance (NMR) spectroscopy was performed on a Bruker Ascend 500 MHz NMR or 600 MHz NMR (IS as TMS). ^13^C Nuclear magnetic resonance (NMR) spectroscopy was performed on a Bruker Ascend 151 MHz NMR (IS as TMS). ^19^F Nuclear magnetic resonance (NMR) spectroscopy was performed on a Bruker Ascend 471 MHz NMR or 565 MHz NMR (IS as TMS). Chemical shifts were reported in parts per million (ppm, δ) downfield from tetramethylsilane. Proton coupling patterns were described as singlet (s), doublet (d), triplet (t), quartet (q), multiplet (m), and broad (br). High-resolution mass spectra (HRMS) and low-resolution mass spectra (LRMS) were obtained by electrospray ionization (ESI) using Thermo Exactive Plus and Agilent 6125C MS. HPLC data analysis of compounds was performed on an Agilent 1290 with a quaternary pump and diode-array detector (DAD), the peak purity was verified by UV spectra. Unless otherwise specified, all compounds are > 95% pure by HPLC analysis. The NMR and MS spectra, HPLC methods and HPLC traces are contained in supporting information.

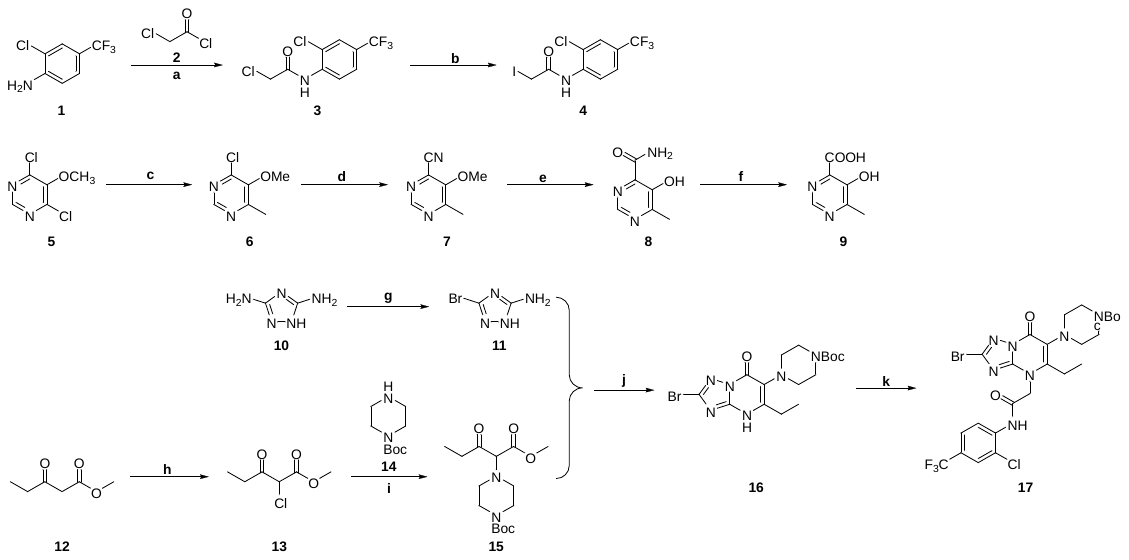

**Supplementary Scheme 1. Synthesis of Intermediate 17*^a^***. *^a^*Reagents and conditions: (a) DCM, 0 ℃, 64%; (b) KI, acetone, 80 ℃, reflux, 96%; (c) MeMgCl, THF, 0 ℃, 80%; (d) Zn(CN)_2_, Pd(PPh_3_)_4_, DMF, 120 ℃, Ar, 50%; (e) 48% HBr aq., 40 ℃, 50%; (f) 1 M NaOH aq., 100 ℃, 84%; (g) NaNO_2_, HBr/H_2_O, 100 ℃, 60%; (h) SOCl_2_, DCM, r.t.; (i) Et_3_N, ACN, 60 ^o^C, 56% for two steps; (j) H_3_PO_4_, EtOH, 80 ^o^C, 45%; (k) **4**, DIPEA, DMF, 0 ℃, 34%.

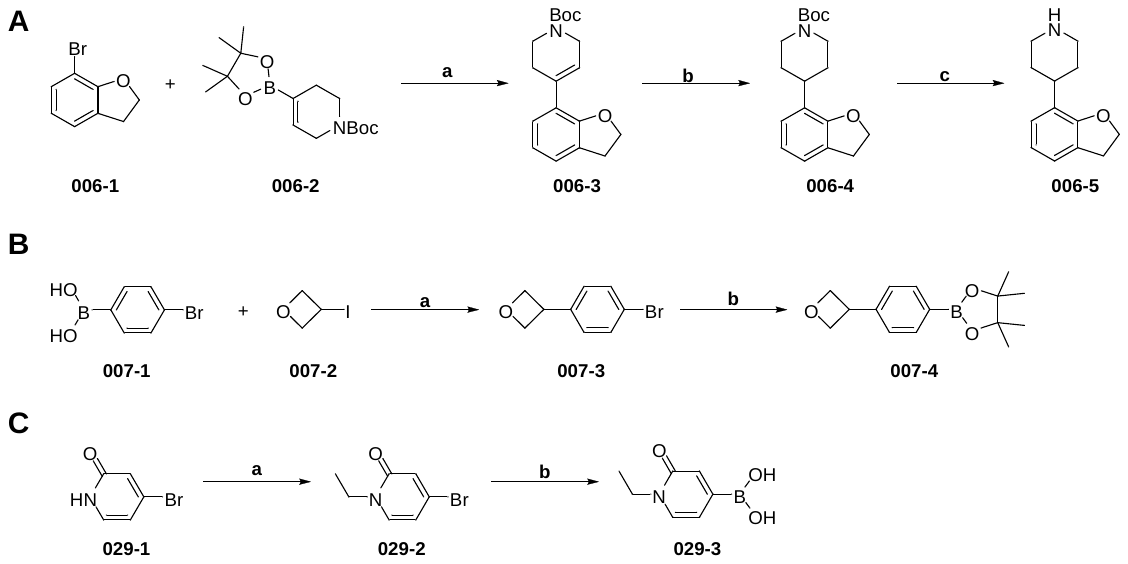

**Supplementary Scheme 2. Synthesis of Intermediates 006-5, 007-4, 029-3*^a^***. *^a^*Reagents and conditions: **A.** (a) Pd(dppf)Cl_2_, K_3_PO_4_, 1,4-dioxane/H_2_O, Ar, 80 ℃, 72%; (b) H_2_, Pd/C, MeOH, r.t.; (c) TFA, DCM, r.t., 56% for two steps. **B.** (a) (*1R, 2R*)-2-aminocyclohexan-1-ol; NaHMDS, NiI_2_, isopropyl alcohol, 80 ^o^C, 67%; (b) B_2_pin_2_, KOAc, Pd(dppf)Cl_2_, 1,4-dioxane, 80 ^o^C, 78%. **C.** (a) (a-i) NaH, THF, 0 ℃; (a-ii) Iodoethane, r.t. to 50 ^o^C, 65%; (b) B_2_pin_2_, KOAc, Pd(dppf)Cl_2_, 1,4-dioxane, 100 ^o^C, 42%.

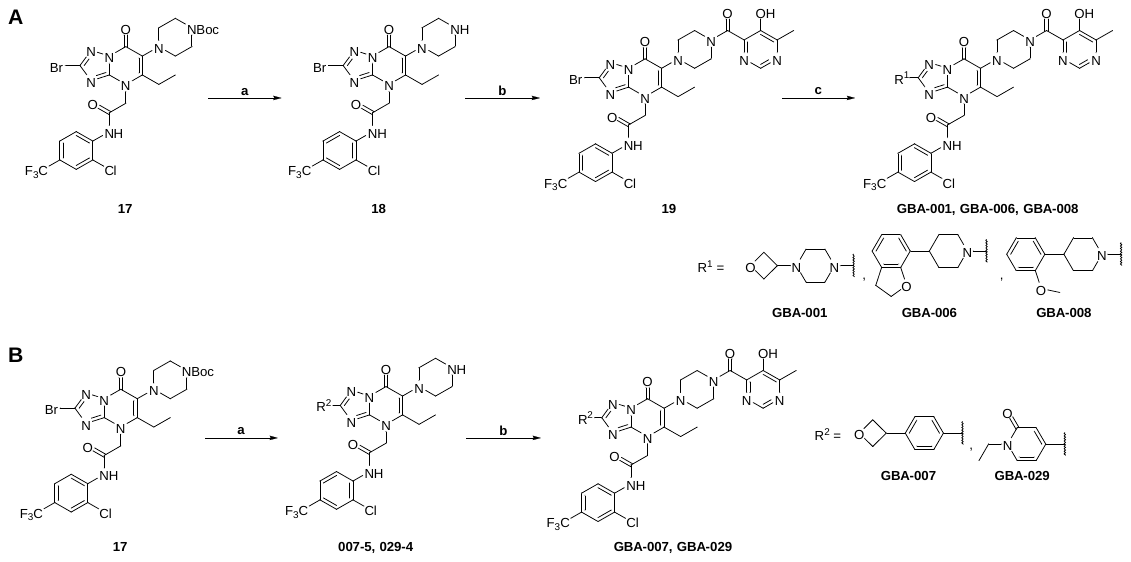

**Supplementary Scheme 3. Synthesis of Compounds GBA-001**, **GBA-006**, **GBA-007**, **GBA-008**, **GBA-029*^a^***. *^a^*Reagents and conditions: **A.** (a) TFA, DCM, r.t., 100%; (b) **9**, HOBt, EDCI, DIPEA, DMF, r.t., 71%-80%; (c) amine derivatives, KOAc, DMF, DMSO, 120 ℃, 12%-36%. **B.** (a) (a-i) borate ester, Pd(dppf)Cl_2_, K_3_PO_4_, 1,4-dioxane/H_2_O, Ar, 80 ℃; (a-ii) TFA, DCM, r.t., 23%-56% for two steps; (b) **9**, HOBt, EDCI, DIPEA, DMF, r.t., 71%-82%.

**4-chloro-5-methoxy-6-methylpyrimidine (6)*.*** 4,6-Dichloro-5-methoxypyrimidine **5** (17.90 g, 100.0 mmol, 1.0 equiv) were dissolved in 150 mL of DCM and cooled to 0 °C before the addition of Methylmagnesium chloride 3 M in THF (37 mL, 110.0 mmol, 1.1 equiv). After stirring for 10.0 h at room temperature, the reaction mixture was quenched with saturated NH_4_Cl solution (50 mL), extracted with EtOAc. The combined organic layers were washed with brine (100 ml), dried over anhydrous Na_2_SO_4_, filtered and concentrated. The residue was purified via silica gel chromatography (20% EA/PE) to afford compound **6** (12.50 g, 80% yield) as a yellow solid.

**5-methoxy-6-methylpyrimidine-4-carbonitrile (7)*.*** To a solution of **6** (3.17 g, 20.0 mmol, 1.0 equiv) in dry DMF (20 mL) was added Zn(CN)_2_ (1.64 g, 14.0 mmol, 0.7 equiv), Pd(PPh_3_)_4_ (1.16 g, 1.0 mmol, 0.05 equiv). Then the mixture was heated at 120 ^o^C for overnight. After reaction completed, the mixture was cooled to ambient temperature. EtOAc (20 mL) was added and the insoluble matter was removed by filtration through 100-200 mesh silica gel, and the filtrate was washed with 10% LiCl (3 × 20 mL), dried over Na_2_SO_4_, filtered and concentrated under reduced pressure. The residue was purified via silica gel chromatography (10-25% EA/PE) to afford the compound **7** (1.49 g, 50% yield) as a yellow solid. ^1^H NMR (500 MHz, DMSO-*d6*) *δ* 8.90 (s, 1H), 4.11 (s, 3H), 2.52 (s, 3H).

**5-hydroxy-6-methylpyrimidine-4-carboxamide (8)*.*** To a solution of **7** (3.00 g, 20.0 mmol, 1.0 equiv) in 48% hydrogen bromide solution (80 mL) was heated at 40 ^o^C for 20.0 h. After the end of the reaction monitored by TLC, the mixture was cooled to ambient temperature. The mixture was cooled in an ice bath and the pH was adjusted to approximately 3 with 50% NaOH. The resulting solid was filtered, washed with water, and dried at 50 °C to afford the compound **8** (1.53 g, 50% yield) as a yellow solid. ^1^H NMR (500 MHz, DMSO-*d_6_*) *δ* 12.69 (s, 1H), 8.80 (s, 1H), 8.62 (s, 1H), 8.47 (s, 1H), 2.44 (s, 3H).

**5-hydroxy-6-methylpyrimidine-4-carboxylic acid (9).** To a solution of **8** (1.53 g, 10.0 mmol, 1.0 equiv) in 1 M aqueous sodium hydroxide solution (45 mL) was heated at 100 ^o^C for 12.0 h. After the end of the reaction monitored by TLC, the mixture was cooled to ambient temperature. The mixture was cooled in an ice bath and the resulting aqueous residue was acidified to pH < 3 with 3 M HCl. Then the residue was filtered and washed with 95% ice-cold ethanol, and dried at 50 °C to afford the compound **9** (1.46 g, 84% yield) as a yellow solid. ^1^H NMR (500 MHz, DMSO-*d_6_*) *δ* 8.66 (s, 1H), 2.46 (s, 3H).

Intermediates **4** and Intermediates **17** were prepared through published synthetic methods^1^.

**2-(2-bromo-5-ethyl-6-(4-(5-hydroxy-6-methylpyrimidine-4-carbonyl)piperazin-1-yl)-7-oxo-[1,2,4]triazolo[1,5-*a*]pyrimidin-4(7*H*)-yl)-*N*-(2-chloro-4-(trifluoromethyl)phenyl)acetamide (19)*.*** To a solution of **17** (1.00 g, 1.5 mmol, 1.0 equiv) was dissolved in DCM (9 mL) and added TFA (3 mL), the mixture was stirred at room temperature for 1.5 h. After reaction completed, the reaction mixture was quenched with 30 mL of saturated aqueous NaHCO₃ and diluted with 30 mL of DCM. The organic phase was separated, and the aqueous phase was extracted with DCM (30 mL × 3). The combined organic phases were washed with brine (30 mL × 3), dried over Na_2_SO_4_, and concentrated to afford intermediate **18**. To a solution of **9** (349 mg, 2.3 mmol, 1.2 equiv) in DMF (7 mL) were added HOBt (51 mg, 0.4 mmol, 0.2 equiv) and EDCI (544 mg, 2.8 mmol, 1.5 equiv) successively. The mixture was stirred room temperature under Ar for 30 min, then **18** (1.06 g, 1.9 mmol, 1.0 equiv) and DIPEA (3 mL, 5.7 mmol, 3.0 equiv ) was added and the mixture was stirred room temperature under Ar for 2.0 h. After reaction completed, the reaction mixture was diluted with 30 mL of DCM and washed with water and brine. The organic phase was dried and removed through rotary evaporation to afford the crude product, which was further purified via silica gel chromatography (0-10% MeOH/DCM) to afford intermediate **19** (940 mg, 72% yield) as a white solid. ^1^H NMR (500 MHz, DMSO-*d_6_*) *δ* 10.37 (s, 1H), 8.49 (s, 1H), 8.08 (d, *J* = 8.6 Hz, 1H), 7.96 (d, *J* = 2.2 Hz, 1H), 7.72 (dd, *J* = 8.8, 2.2 Hz, 1H), 5.30 (s, 2H), 4.54 – 4.49 (m, 1H), 3.51 – 3.19 (m, 4H), 3.03 – 2.94 (m, 3H), 2.84 – 2.78 (m, 1H), 2.65 – 2.60 (m, 1H), 2.42 (s, 3H), 1.17 (t, *J* = 7.5 Hz, 3H).

***N*-(2-chloro-4-(trifluoromethyl)phenyl)-2-(5-ethyl-6-(4-(5-hydroxy-6-methylpyrimidine-4-carbonyl)piperazin-1-yl)-2-(4-(oxetan-3-yl)piperazin-1-yl)-7-oxo-[1,2,4]triazolo[1,5-*a*]pyrimidin-4(7*H*)-yl)acetamide (GBA-001)*.*** To a solution of **19** (70 mg, 0.1 mmol, 1.0 equiv) and 1-(Oxetan-3-yl)piperazine (22 mg, 0.2 mmol, 1.5 equiv) in DMSO/DMF (1.0 mL/1.0 mL) were added KOAc (59 mg, 0.6 mmol, 6.0 equiv) successively. The mixture was stirred at 120 °C under Ar for 16.0 h. After reaction completed, the mixture was cooled to ambient temperature and concentrated under reduced pressure. Water (10 mL) was added and the layers were separated. The aqueous layer was extracted with EtOAc (3 × 10 mL). The combined organic layers were washed with brine (10 mL), dried over Na_2_SO_4_, filtered and concentrated under reduced pressure. The residue was purified via flash chromatography (acetonitrile : water 0.1% HCOOH = 10-100%) to afford the title compound **GBA-001** (27 mg, 36% yield) as a white solid. ^1^H NMR (500 MHz, DMSO-*d_6_*) *δ* 10.34 (s, 1H), 10.23 (s, 1H), 8.57 (s, 1H), 8.03 (d, *J* = 8.6 Hz, 1H), 7.96 (d, *J* = 2.2 Hz, 1H), 7.72 (dd, *J* = 8.8, 2.2 Hz, 1H), 5.21 (s, 2H), 4.56 – 4.51 (m, 2H), 4.48 – 4.41 (m, 2H), 3.55 – 3.38 (m, 9H), 3.27 – 3.18 (m, 1H), 3.01 – 2.89 (m, 3H), 2.82 – 2.75 (m, 1H), 2.65 – 2.55 (m, 1H), 2.44 (s, 3H), 2.37 – 2.25 (m, 4H), 1.16 (t, *J* = 7.4 Hz, 3H); ^13^C NMR (151 MHz, DMSO-*d_6_*) *δ* 166.17, 164.38, 163.97, 156.89, 154.59, 153.60, 150.91, 148.89, 146.90, 146.44, 138.14, 126.83 (d, *J* = 4.0 Hz), 126.43 (d, *J* = 32.9 Hz), 126.06, 125.50, 124.79 (d, *J* = 4.1 Hz), 123.37 (q, *J* = 272.3 Hz), 121.62, 74.41, 58.47, 50.35, 50.16, 49.83, 48.36, 47.02, 45.24, 41.95, 31.20, 29.07, 22.15, 20.76, 19.29, 12.90; ^19^F NMR (471 MHz, DMSO-*d_6_*) *δ* -60.80; HRMS (ESI) calcd for C_33_H_37_ClF_3_N_11_O_5_ 760.2692 [M+H]^+^, found 760.2690; HPLC retention time = 6.179 min, purity = 98.45%, (λ = 254 nm).

**4-(2,3-dihydrobenzofuran-7-yl)piperidine (006-5)*.*** To a solution of 7-bromo-2,3-dihydrobenzofuran **006-1** (1.00 g, 5.0 mmol, 1.0 equiv) and tert-butyl 4-(4,4,5,5-tetramethyl-1,3,2-dioxaborolan-2-yl)-3,6-dihy- dropyridine-1(2*H*)-carboxylate **006-2** (1.70 g, 5.0 mmol, 1.1 equiv) in 1,4-dioxane/H_2_O (16/8 mL) were added K_3_PO_4_ (2.65 g, 12.5 mmol, 2.5 equiv) and Pd(dppf)Cl_2_ (183 mg, 0.3 mmol, 0.05 equiv) successively. The mixture was stirred at 80 °C under Ar for 16.0 h. After reaction completed, the mixture was cooled to ambient temperature. The reaction mixture was filtered through a pad of Celite, and the filtrate was concentrated under reduced pressure. EtOAc (20 mL) and water (10 mL) were added and the layers were separated. The aqueous layer was extracted with EtOAc (4 × 20 mL). The combined organic layers were washed with brine (10 mL), dried over Na_2_SO_4_, filtered and concentrated under reduced pressure. The residue was purified via silica gel chromatography (0-5% MeOH/DCM) to afford the compound **006-3** (1.00 g, 72% yield).

To a solution of **006-3** (500 mg, 1.7 mmol) in MeOH was added 10% Pd/C catalyst. The reaction flask was charged with H_2_ (1 atm) and then stirred 2 h at room temperature. The reaction mixture was filtered over Celite under argon and concentrated to afford the crude intermediate **006-4**. TFA (1 mL) was added to a stirred solution of crude intermediate in DCM (3 mL) and then the mixture stirred at room temperature for 1.5 h. After reaction completed, the reaction mixture was quenched with 10 mL of saturated aqueous NaHCO₃ and diluted with 20 mL of DCM. The organic phase was separated, and the aqueous phase was extracted with DCM (20 mL × 3). The combined organic layers were washed with brine (10 mL), dried over Na_2_SO_4_, filtered and concentrated to afford the crude product, which was further purified via silica gel chromatography (0-10% MeOH/DCM) to afford intermediate **006-5** (188 mg, 56% yield for two steps) as a white solid. ^1^H NMR (600 MHz, DMSO-*d_6_*) *δ* 7.11 (d, *J* = 7.3 Hz, 1H), 6.92 (d, *J* = 7.6 Hz, 1H), 6.83-6.78 (m, 1H), 4.52 (t, *J* = 8.7 Hz, 2H), 3.38 – 3.35 (m, 1H), 3.35 – 3.33 (m, 1H), 3.17 (t, *J* = 8.7 Hz, 2H), 3.03 – 2.89 (m, 3H), 1.92 – 1.79 (m, 4H).

***N*-(2-chloro-4-(trifluoromethyl)phenyl)-2-(2-(4-(2,3-dihydrobenzofuran-7-yl)piperidin-1-yl)-5-ethyl-6-(4-(5-hydroxy-6-methylpyrimidine-4-carbonyl)piperazin-1-yl)-7-oxo-[1,2,4]triazolo[1,5-*a*]pyrimidin-4(7*H*)-yl)acetamide (GBA-006)*.*** Compound **GBA-006** was synthesized with the similar procedure as **GBA-001** from **006-5** and **19**. The residue was purified via flash chromatography (acetonitrile : water 0.1% CF_3_COOH = 10-100%) to afford the title compound **GBA-006** as a brown liquid (10 mg, 12% yield). ^1^H NMR (600 MHz, DMSO-*d_6_*) *δ* 10.34 (s, 1H), 10.24 (s, 1H), 8.58 (s, 1H), 8.04 (d, *J* = 8.6 Hz, 1H), 7.96 (d, *J* = 2.1 Hz, 1H), 7.71 (dd, *J* = 8.8, 2.1 Hz, 1H), 7.06 (d, *J* = 7.2 Hz, 1H), 6.92 (d, *J* = 7.6 Hz, 1H), 6.78-6.72 (m, 1H), 5.22 (s, 2H), 4.54 – 4.47 (m, 3H), 4.29 – 4.11 (m, 2H), 3.55 – 3.45 (m, 3H), 3.28 – 3.19 (m, 1H), 3.15 (t, *J* = 8.7 Hz, 2H), 3.02 – 2.91 (m, 5H), 2.90 – 2.82 (m, 1H), 2.80 – 2.76 (m, 1H), 2.63 – 2.58 (m, 1H), 2.44 (s, 3H), 1.77 – 1.72 (m, 2H), 1.70 – 1.59 (m, 2H), 1.16 (t, *J* = 7.5 Hz, 3H); ^13^C NMR (151 MHz, DMSO-*d_6_*) *δ* 166.22, 164.35, 163.98, 157.10, 156.86, 154.41, 153.61, 150.89, 148.92, 146.92, 146.37, 138.15, 127.00, 126.82, 126.79 (d, *J* = 3.8 Hz), 126.42, 123.36 (d, *J* = 33.3 Hz), 126.09, 125.53, 125.03, 124.76 (d, *J* = 3.9 Hz), 124.26, 123.36 (q, *J* = 272.3 Hz), 122.78, 121.57, 120.45, 70.50, 50.36, 50.16, 49.84, 47.02, 46.06, 41.96, 36.14, 30.38, 29.34, 20.73, 19.28, 12.91; ^19^F NMR (565 MHz, DMSO-*d_6_*) *δ* -60.82, -74.42 (CF_3_COOH); HRMS (ESI) calcd for C_39_H_40_ClF_3_N_10_O_5_ 821.2896 [M+H]^+^, found 821.2895; HPLC retention time = 10.921 min, purity = 97.34%, (λ = 254 nm).

**3-(4-bromophenyl)oxetane (007-3)*.*** To a solution of (4-bromophenyl)boronic acid **007-1** (709 mg, 3.5 mmol, 1.0 equiv) in isopropyl alcohol (5 mL) was added (*1R, 2R*)-2-aminocyclohexan-1-ol (31 mg, 0.3 mmol, 0.08 equiv ), NaHMDS (5 mL, 5.2 mmol, 1.5 equiv) and NiI_2_ (85 mg, 0.3 mmol, 0.08 equiv) was degassed with Ar for 5 min, then 3-iodooxetane **007-2** (500 mg, 2.7 mmol, 0.8 equiv) was added and the mixture was heated at 80 ^o^C under microwave irradiation for 0.5 h. After cooled to room temperature, the mixture was quenched by saturated NH_4_Cl solution (5 mL), extracted with EtOAc. The combined organic layers were washed with brine (5 mL), dried over anhydrous Na_2_SO_4_, filtered and concentrated. The residue was purified via silica gel chromatography (0-5% EA/PE) to afford the compound **007-3** (500 mg, 67% yield) as a colorless liquid. ^1^H NMR (500 MHz, Chloroform-*d*) *δ* 7.49 (d, *J* = 8.4 Hz, 2H), 7.27 (d, *J* = 8.4 Hz, 2H), 5.07 (dd, *J* = 8.4, 6.0 Hz, 2H), 4.74 – 4.68 (m, 2H), 4.21 – 4.13 (m, 1H).

**4,4,5,5-tetramethyl-2-(4-(oxetan-3-yl)phenyl)-1,3,2-dioxaborolane (007-4)*.*** To a solution of **007-3** (500 mg, 2.4 mmol, 1.0 equiv) and bis(pinacolato)diboron (337 mg, 4.7 mmol, 2.0 equiv) in 1,4-dioxane (6 mL) were added KOAc (694 mg, 7.1 mmol, 3.0 equiv) and Pd(dppf)Cl_2_ (345 mg, 0.5 mmol, 0.2 equiv) successively. The mixture was stirred at 80 °C under Ar for 16.0 h. After reaction completed, the mixture was cooled to ambient temperature. The reaction mixture was filtered through a pad of Celite, and the filtrate was concentrated under reduced pressure. EtOAc (20 mL) and water (10 mL) were added and the layers were separated. The aqueous layer was extracted with EtOAc (4 × 20 mL). The combined organic layers were washed with brine (10 mL), dried over Na_2_SO_4_, filtered and concentrated under reduced pressure. The residue was purified via silica gel chromatography (0-10% EA/PE) to afford the title compound **007-4** (478 mg, 78% yield) as a white solid. ^1^H NMR (500 MHz, Chloroform-*d*) *δ* 7.81 (d, *J* = 7.7 Hz, 2H), 7.40 (d, *J* = 7.6 Hz, 2H), 5.08 (dd, *J* = 8.4, 6.0 Hz, 2H), 4.81 – 4.74 (m, 2H), 4.24 (p, *J* = 7.6 Hz, 1H), 1.35 (s, 12H).

***N*-(2-chloro-4-(trifluoromethyl)phenyl)-2-(5-ethyl-2-(4-(oxetan-3-yl)phenyl)-7-oxo-6-(piperazin-1-yl)-[1,2,4]triazolo[1,5-*a*]pyrimidin-4(7*H*)-yl)acetamide (007-5)*.*** To a solution of **17** (200 mg, 0.3 mmol, 1.0 equiv) and **007-4** (117 mg, 0.5 mmol, 1.5 equiv) in 1,4-dioxane/H_2_O (2 mL/100 µL) were added K_3_PO_4_ (192 mg, 0.9 mmol, 3.0 equiv) and Pd(dppf)Cl_2_ (44 mg, 0.06 mmol, 0.2 equiv) successively. The mixture was stirred at 80 °C under Ar for 16.0 h. After reaction completed, the mixture was cooled to ambient temperature. The reaction mixture was filtered through a pad of Celite, and the filtrate was concentrated under reduced pressure. EtOAc (20 mL) and water (10 mL) were added and the layers were separated. The aqueous layer was extracted with EtOAc (3 × 10 mL). The combined organic layers were washed with brine (10 mL), dried over Na_2_SO_4_, filtered and concentrated under reduced pressure to afford the crude product. TFA (1 mL) was added to a stirred solution of crude product in DCM (3 mL) and then the mixture stirred at room temperature for 1.5 h. After reaction completed, the reaction mixture was quenched with 10 mL of saturated aqueous NaHCO₃ and diluted with 20 mL of DCM. The organic phase was separated, and the aqueous phase was extracted with DCM (20 mL × 3). The combined organic layers were washed with brine (10 mL), dried over Na_2_SO_4_, filtered and concentrated under reduced pressure. The residue was purified via silica gel chromatography (0-10% MeOH/DCM) to afford intermediates **007-5** (43 mg, 23% yield) as a yellow liquid.

***N*-(2-chloro-4-(trifluoromethyl)phenyl)-2-(5-ethyl-6-(4-(5-hydroxy-6-methylpyrimidine-4-carbonyl)piperazin-1-yl)-2-(4-(oxetan-3-yl)phenyl)-7-oxo-[1,2,4]triazolo[1,5-*a*]pyrimidin-4(7*H*)-yl)acetamide (GBA-007)*.*** To a solution of **9** (13 mg, 0.8 mmol, 1.2 equiv) in DMF (3 mL) were added HOBt (2 mg, 0.02 mmol, 0.2 equiv) and EDCI (20 mg, 0.1 mmol, 1.5 equiv) successively. The mixture was stirred room temperature under Ar for 0.5 h, then **007-5** (43 mg, 0.07 mmol, 1.0 equiv) and DIPEA (37 μL, 0.2 mmol, 3.0 equiv ) was added and the mixture was stirred room temperature under Ar for 2.0 h. After reaction completed, the reaction mixture was diluted with 15 mL of DCM and washed with water and brine. The organic phase was dried and removed through rotary evaporation to afford the crude product, which was further purified through flash chromatography (acetonitrile : water 0.1% HCOOH = 10-100%) to afford the title compound **GBA-007** (42 mg, 80% yield) as a white solid. ^1^H NMR (500 MHz, DMSO-*d_6_*) *δ* 10.44 (s, 1H), 8.56 (s, 1H), 8.11 (d, *J* = 8.0 Hz, 2H), 8.05 (d, *J* = 8.6 Hz, 1H), 7.96 (d, *J* = 2.1 Hz, 1H), 7.71 (dd, *J* = 8.8, 2.1 Hz, 1H), 7.55 (d, *J* = 8.1 Hz, 2H), 5.40 (s, 2H), 4.96 (dd, *J* = 8.4, 5.9 Hz, 2H), 4.70 – 4.59 (m, 2H), 4.57 – 4.48 (m, 1H), 4.36 – 4.25 (m, 1H), 3.59 – 3.50 (m, 3H), 3.28 – 3.23 (m, 1H), 3.04 – 2.96 (m, 3H), 2.87 – 2.80 (m, 1H), 2.72 – 2.60 (m, 1H), 2.44 (s, 3H), 1.21 (t, *J* = 7.4 Hz, 3H); ^13^C NMR (151 MHz, DMSO-*d_6_*) *δ* 166.10, 164.49, 160.78, 156.89, 156.79, 154.07, 151.78, 148.54, 146.85, 144.29, 138.12, 128.76, 128.18, 127.39, 127.07, 126.81 (q, *J* = 3.9 Hz), 126.38 (q, *J* = 33.1 Hz), 126.01, 125.43, 124.76 (q, *J* = 3.7 Hz), 123.32 (q, *J* = 272.1 Hz), 121.65, 77.28, 50.50, 50.30, 49.78, 46.96, 41.87, 40.06, 21.05, 19.29, 12.79; ^19^F NMR (471 MHz, DMSO-*d_6_*) *δ* -60.83; HRMS (ESI) calcd for C_35_H_33_O_5_N_9_ClF_3_^+^ 752.2318 [M+H]^+^, found 752.2313; HPLC retention time = 14.130 min, purity = 97.36%, (λ = 254 nm).

***N*-(2-chloro-4-(trifluoromethyl)phenyl)-2-(5-ethyl-6-(4-(5-hydroxy-6-methylpyrimidine-4-carbonyl)piperazin-1-yl)-2-(4-(2-methoxyphenyl)piperidin-1-yl)-7-oxo-[1,2,4]triazolo[1,5-*a*]pyrimidin-4(7*H*)-yl)acetamide (GBA-008)*.*** Compound **GBA-008** was synthesized with the similar procedure as **GBA-001** from **19** and 4-(2-Methoxyphenyl)piperidine. **GBA-008** was obtained as a yellow solid (12 mg, 15% yield). ^1^H NMR (500 MHz, Chloroform-*d*) *δ* 11.79 (s, 1H), 9.18 (s, 1H), 8.58 (s, 1H), 8.47 (d, *J* = 8.6 Hz, 1H), 7.64 (s, 1H), 7.53 (d, *J* = 8.6 Hz, 1H), 7.22 – 7.16 (m, 1H), 7.13 (d, *J* = 7.5 Hz, 1H), 6.98 – 6.90 (m, 1H), 6.87 (d, *J* = 8.1 Hz, 1H), 5.75 – 5.50 (m, 1H), 5.03 (s, 2H), 4.87 – 4.63 (m, 1H), 4.53 – 4.28 (m, 2H), 3.89 – 3.73 (m, 5H), 3.54 – 3.38 (m, 1H), 3.20 – 3.00 (m, 6H), 2.91 – 2.65 (m, 2H), 2.55 (s, 3H), 1.93 – 1.83 (m, 2H), 1.75 – 1.64 (m, 2H), 1.30 (t, *J* = 7.4 Hz, 3H); ^13^C NMR (151 MHz, DMSO-*d_6_*) *δ* 166.21, 164.42, 161.04, 156.92, 156.69, 154.14, 151.75, 150.93, 148.96, 148.87, 146.93, 146.41, 138.16, 129.71, 126.86 (d, *J* = 4.2 Hz), 126.84, 126.49 (d, *J* = 33.0 Hz), 126.14, 125.55, 124.81 (d, *J* = 3.8 Hz), 123.37 (q, *J* = 272.2 Hz), 122.15, 121.62, 120.16, 111.81, 109.55, 55.66, 55.57, 50.52, 50.34, 49.82, 47.03, 41.96, 35.17, 29.08, 21.09, 19.31, 12.85; ^19^F NMR (565 MHz, DMSO-*d_6_*) *δ* -60.83; HRMS (ESI) calcd for C_38_H_40_ClF_3_N_10_O_5_ 809.2896 [M+H]^+^, found 809.2894; HPLC retention time = 10.924 min, purity = 96.87%, (λ = 254 nm).

**4-bromo-1-ethylpyridin-2(1*H*)-one (029-2)*.*** To a solution of 4-bromopyridin-2(1H)-one **029-1** (200 mg, 1.2 mmol, 1.0 equiv) in dry THF (5 mL) was added sodium hydride (30 mg, 1.3 mmol, 1.1 equiv) successively. The mixture was stirred at 0 °C under Ar for 0.5 h, then iodoethane (300 μL, 3.5 mmol, 3.0 equiv) was added and the mixture was heated at 50 ^o^C for 24.0 h. After cooled to room temperature, the mixture was quenched by saturated NH_4_Cl solution (10 mL), extracted with EtOAc. The combined organic layers was washed with brine (10 mL), dried over anhydrous Na_2_SO_4_, filtered and concentrated. The residue was purified via silica gel chromatography (0-40% EA/PE) to afford the compound **029-2** (150 mg, 65% yield) as a yellow solid. ^1^H NMR (500 MHz, Chloroform-*d*) *δ* 7.12 (d, *J* = 7.2 Hz, 1H), 6.76 (d, *J* = 2.2 Hz, 1H), 6.30 (dd, *J* = 7.2, 2.2 Hz, 1H), 3.91 (q, *J* = 7.2 Hz, 2H), 1.30 (t, *J* = 7.3 Hz, 3H).

**1-ethyl-4-(4,4,5,5-tetramethyl-1,3,2-dioxaborolan-2-yl)pyridin-2(1*H*)-one (029-3)*.*** To a solution of **029-2** (100 mg, 0.5 mmol, 1.0 equiv) and bis(pinacolato)diboron (314 mg, 1.2 mmol, 2.5 equiv) in 1,4-dioxane (6 mL) were added KOAc (146 mg, 1.5 mmol, 3.0 equiv) and Pd(dppf)Cl_2_ (18 mg, 0.03 mmol, 0.05 equiv) successively. The mixture was stirred at 100 °C under Ar for 16.0 h. After reaction completed, the mixture was cooled to ambient temperature. The reaction mixture was filtered through a pad of Celite, and the filtrate was concentrated under reduced pressure. EtOAc (20 mL) and water (10 mL) were added and the layers were separated. The aqueous layer was extracted with EtOAc (4 × 20 mL). The combined organic layers were washed with brine (10 mL), dried over Na_2_SO_4_, filtered and concentrated under reduced pressure to afford the crude product **029-3**. The crude product **029-3** was carried directly to the next step.

***N*-(2-chloro-4-(trifluoromethyl)phenyl)-2-(5-ethyl-2-(1-ethyl-2-oxo-1,2-dihydropyridin-4-yl)-6-(4-(5-hydroxy-6-methylpyrimidine-4-carbonyl)piperazin-1-yl)-7-oxo-[1,2,4]triazolo[1,5-*a*]pyrimidin-4(7*H*)-yl)acetamide (GBA-029).** Compound **GBA-029** was synthesized with the similar procedure as **GBA-007**. The residue was purified via flash chromatography (acetonitrile : water 0.1% CF_3_COOH = 10-100%) to afford the title compound **GBA-029** as a white solid (36 mg, 71% yield). ^1^H NMR (600 MHz, DMSO-*d_6_*) *δ* 10.42 (s, 1H), 10.25 (s, 1H), 8.58 (s, 1H), 8.03 (d, *J* = 8.6 Hz, 1H), 7.97 (d, *J* = 2.1 Hz, 1H), 7.85 (d, *J* = 7.0 Hz, 1H), 7.71 (dd, *J* = 8.8, 2.1 Hz, 1H), 7.03 (d, *J* = 1.8 Hz, 1H), 6.84 (dd, *J* = 6.9, 1.9 Hz, 1H), 5.38 (s, 2H), 4.60 – 4.47 (m, 1H), 3.94 (q, *J* = 7.1 Hz, 2H), 3.55 – 3.46 (m, 3H), 3.32 – 3.23 (m, 1H), 3.12 – 2.94 (m, 3H), 2.87 – 2.77 (m, 1H), 2.68 – 2.64 (m, 1H), 2.44 (s, 3H), 1.27 – 1.19 (m, 6H); ^13^C NMR (151 MHz, DMSO-*d_6_*) *δ* 165.99, 164.35, 160.99, 158.57, 157.40, 156.84, 153.94, 151.85, 148.89, 146.80, 146.38, 140.17, 139.59, 138.06, 126.81 (q, *J* = 3.9 Hz), 126.44 (d, *J* = 32.8 Hz), 126.10, 125.52, 124.74 (q, *J* = 3.8 Hz), 123.31 (q, *J* = 272.3 Hz), 121.77, 116.99, 102.51, 50.54, 50.25, 49.74, 46.96, 43.76, 41.89, 21.10, 19.24, 14.40, 12.75; ^19^F NMR (565 MHz, DMSO-*d_6_*) *δ* -60.83, -74.32 (CF_3_COOH); HRMS (ESI) calcd for C_33_H_32_ClF_3_N_10_O_5_ 741.2270 [M+H]^+^, found 741.2266; HPLC retention time = 7.811 min, purity = 96.55%, (λ = 254 nm).

***N*-(2-chloro-4-(trifluoromethyl)phenyl)-2-(2-(3,6-dihydro-2*H*-pyran-4-yl)-5-ethyl-6-(4-(5-hydroxy-6-methylpyrimidine-4-carbonyl)piperazin-1-yl)-7-oxo-[1,2,4]triazolo[1,5-*a*]pyrimidin-4(7*H*)-yl)acetamide**.(**HRO761**)**.** The WRN inhibitor **HRO761** was synthesized following an established synthesis route^1^.

**(*S*,*E*)-*N*-(1-cyclopropyl-3-(methylsulfonyl)allyl)-2-(1,1-difluoroethyl)-4-phenoxypyrimidine-5-carboxamide (VVD-214).** The WRN inhibitor **VVD-214** (CAS No. 3026500-20-6) was available to purchase from Bide Pharmatech Co., Ltd.

**^1^H NMR of GBA-001**

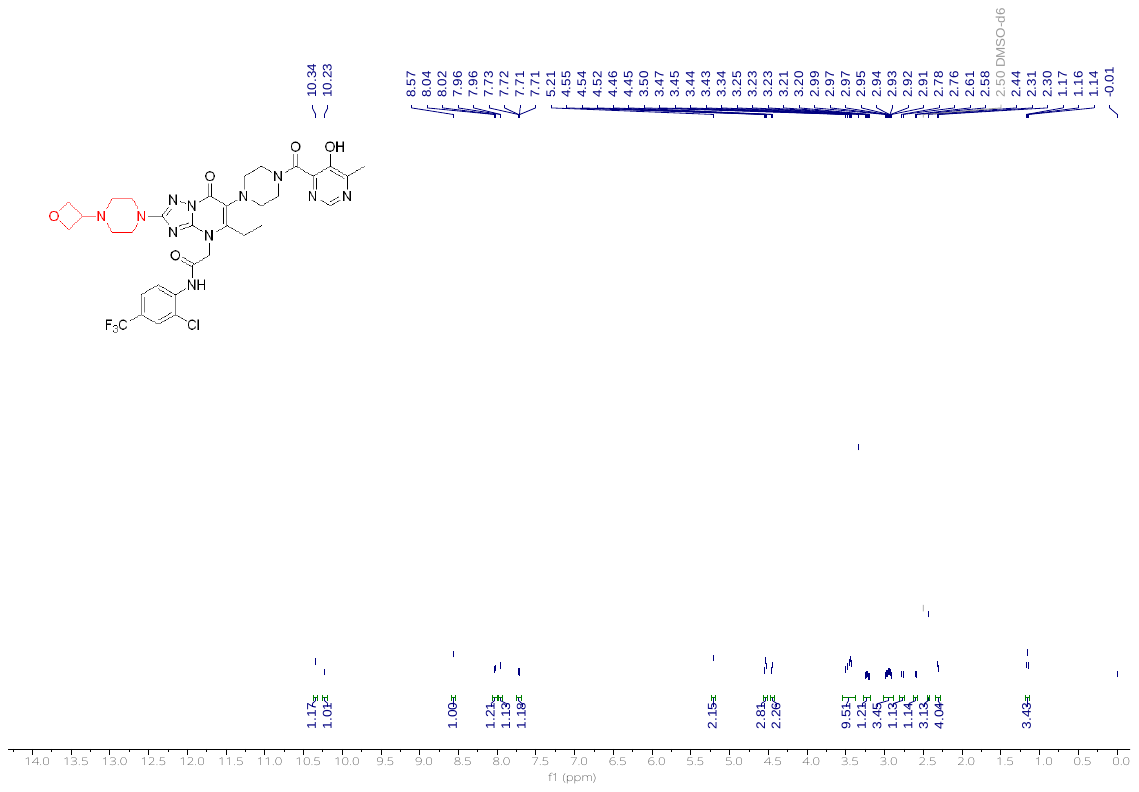

**^13^C NMR of GBA-001**

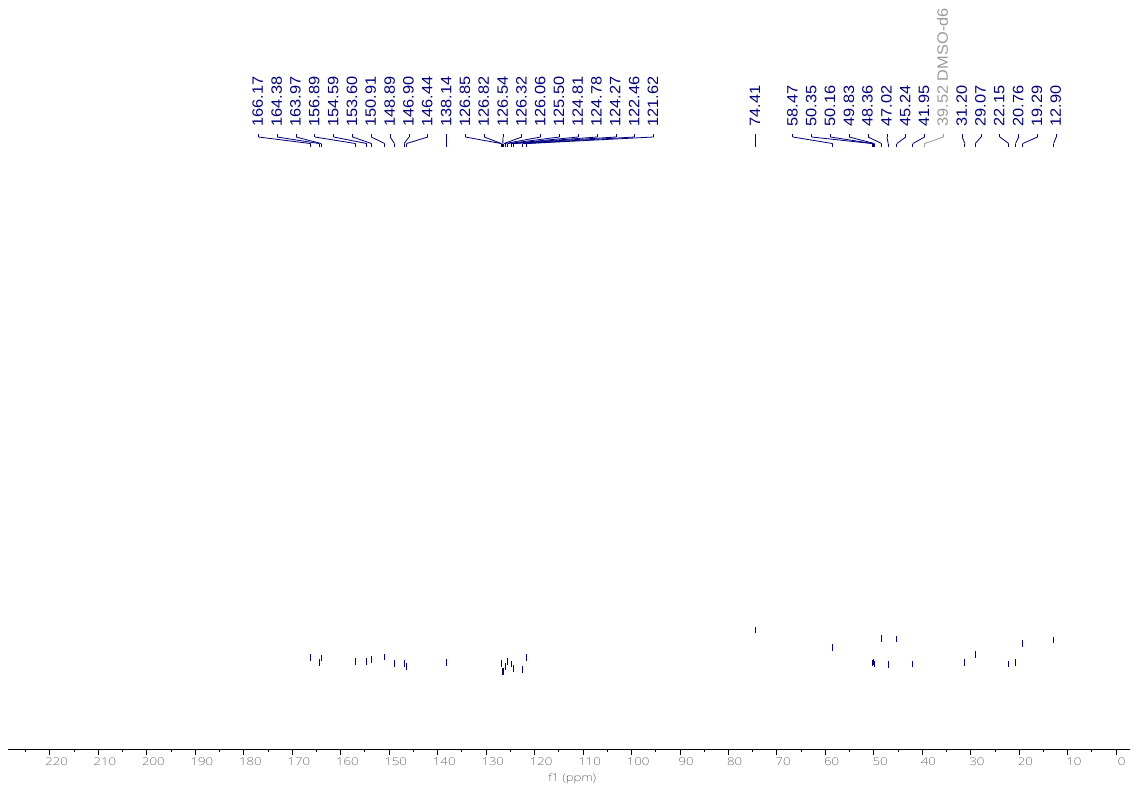

**^19^F NMR of GBA-001**

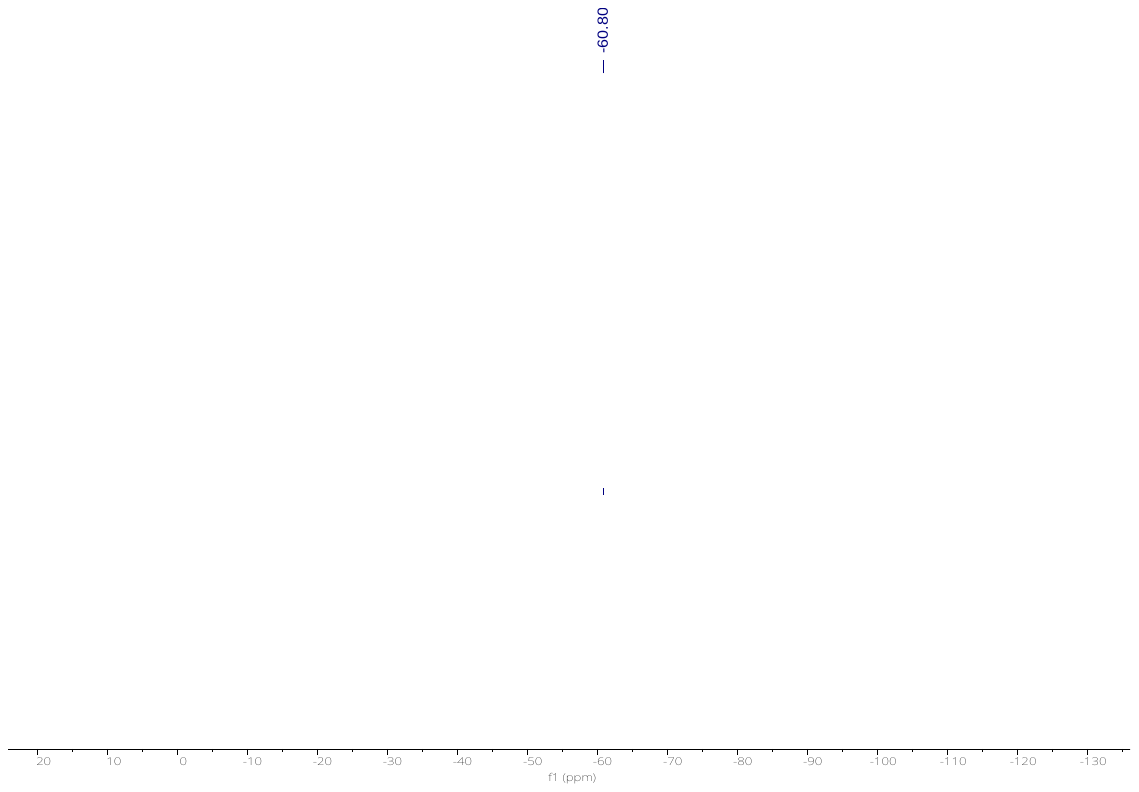

**HRMS of GBA-001**

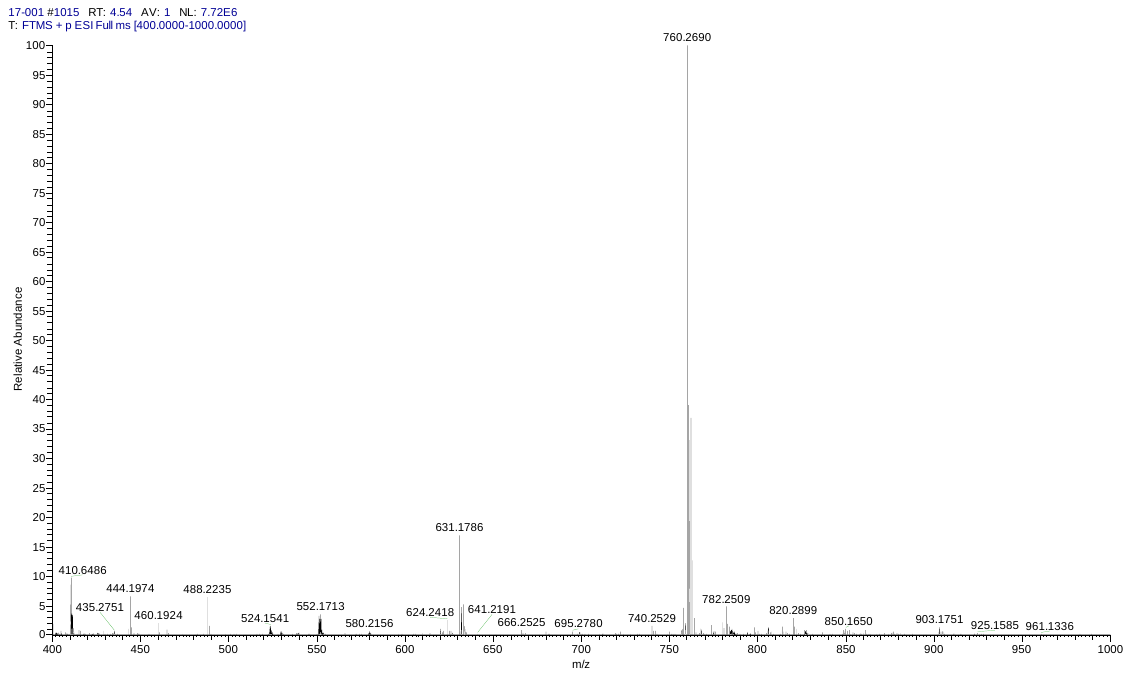

**HPLC of GBA-001**

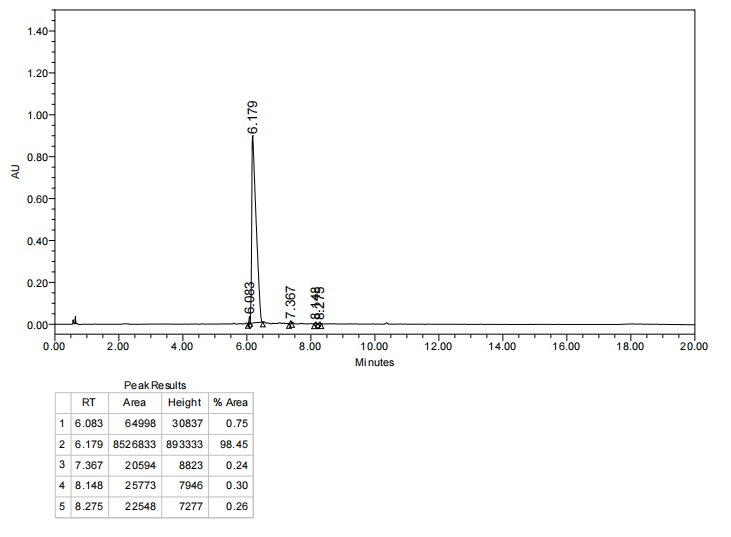

**^1^H NMR of GBA-006**

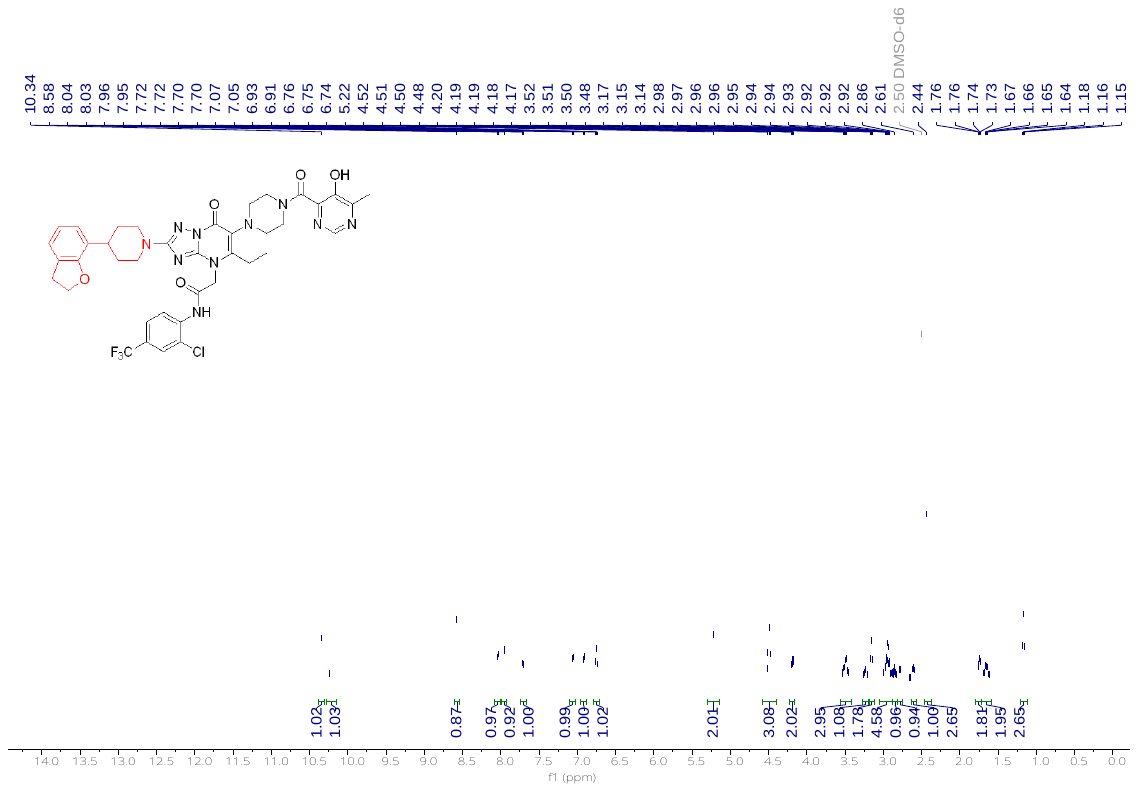

**^13^C NMR of GBA-006**

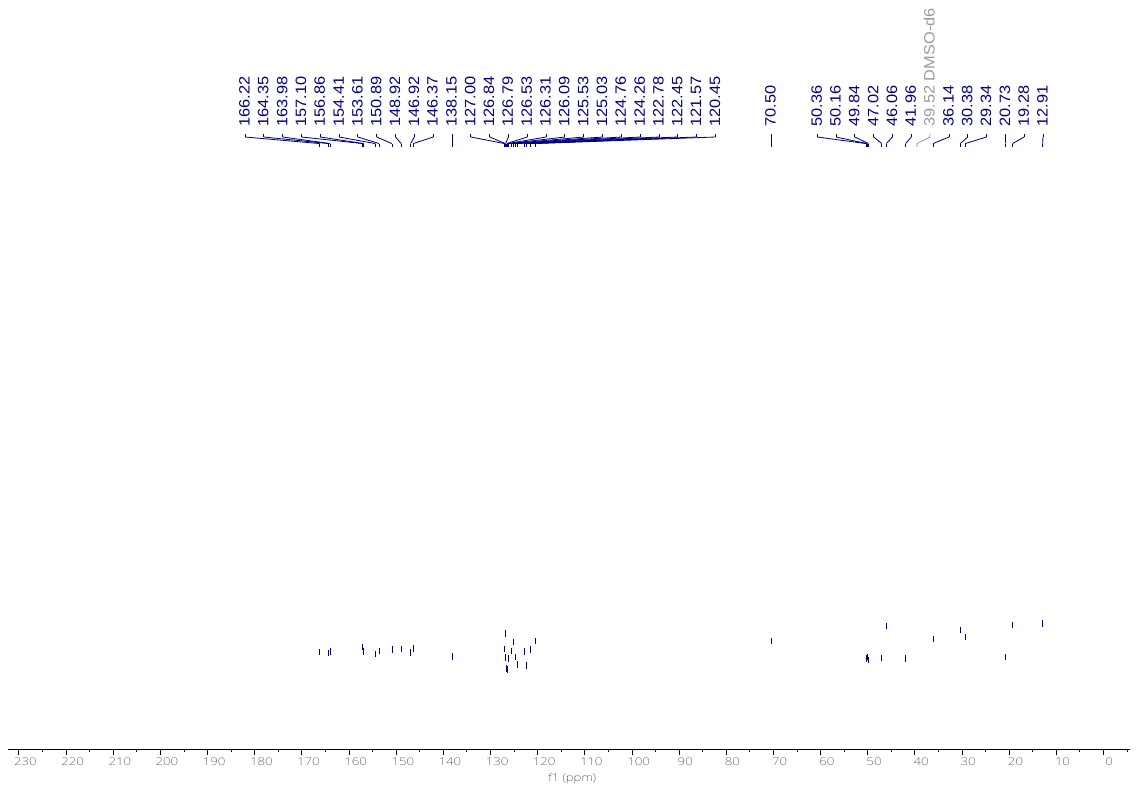

**^19^F NMR of GBA-006**

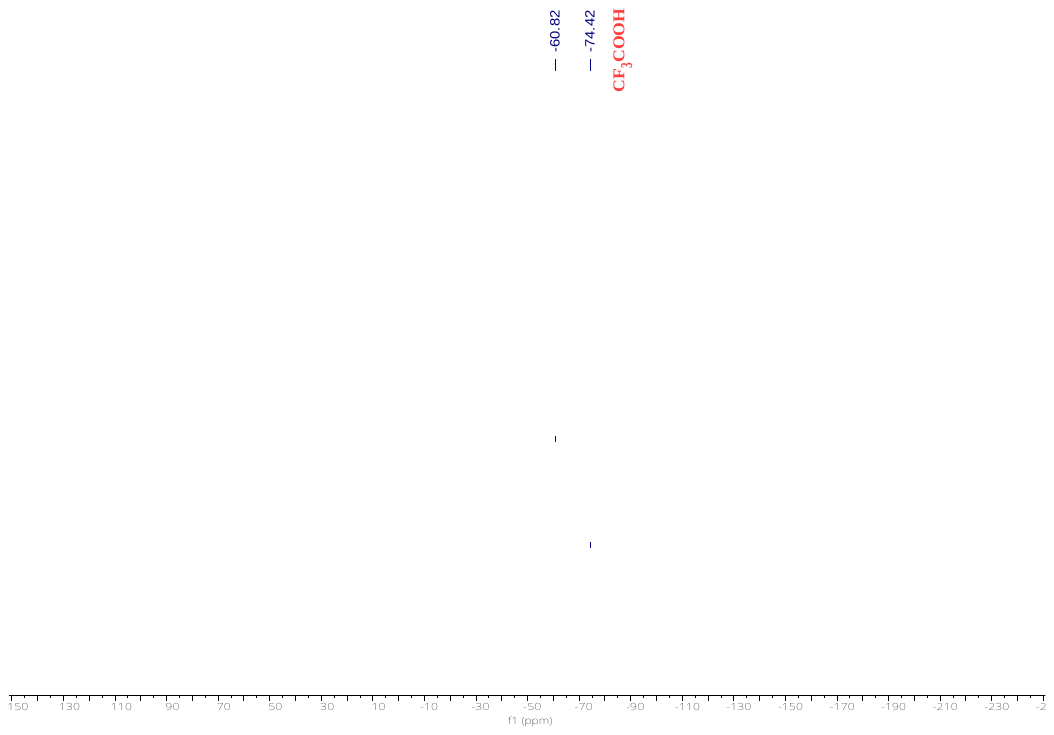

**HRMS of GBA-006**

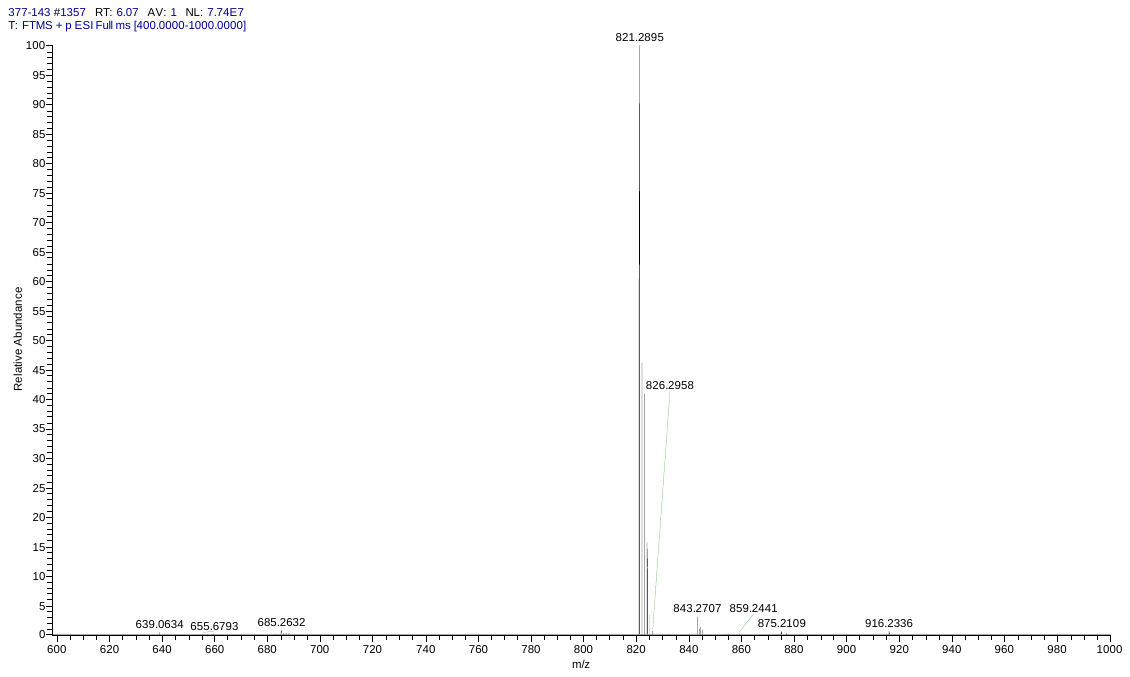

**HPLC of GBA-006**

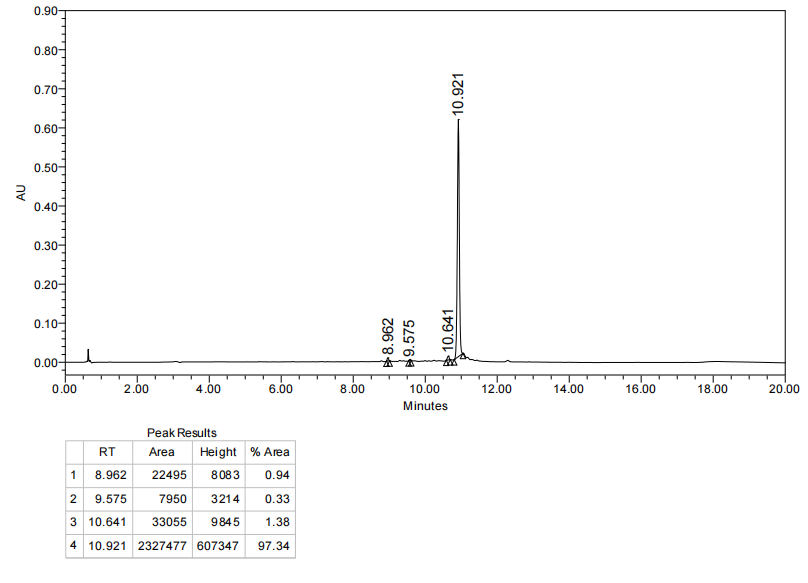

**^1^H NMR of GBA-007**

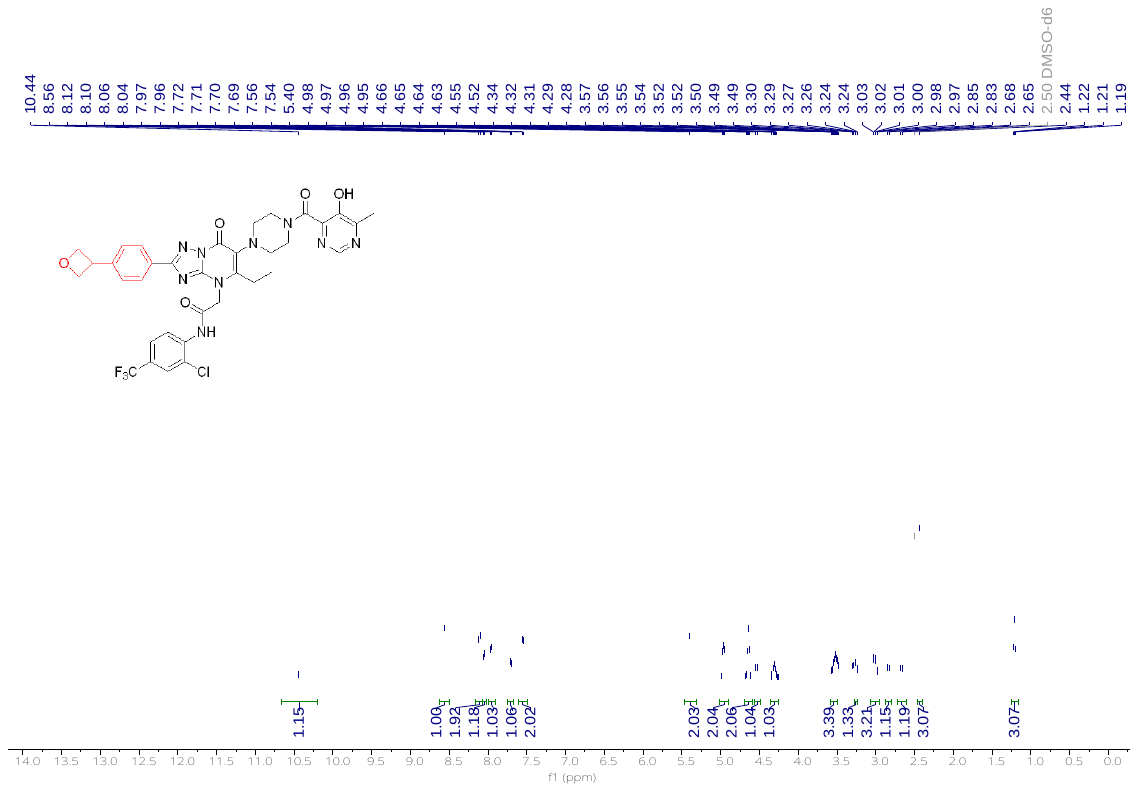

**^13^C NMR of GBA-007**

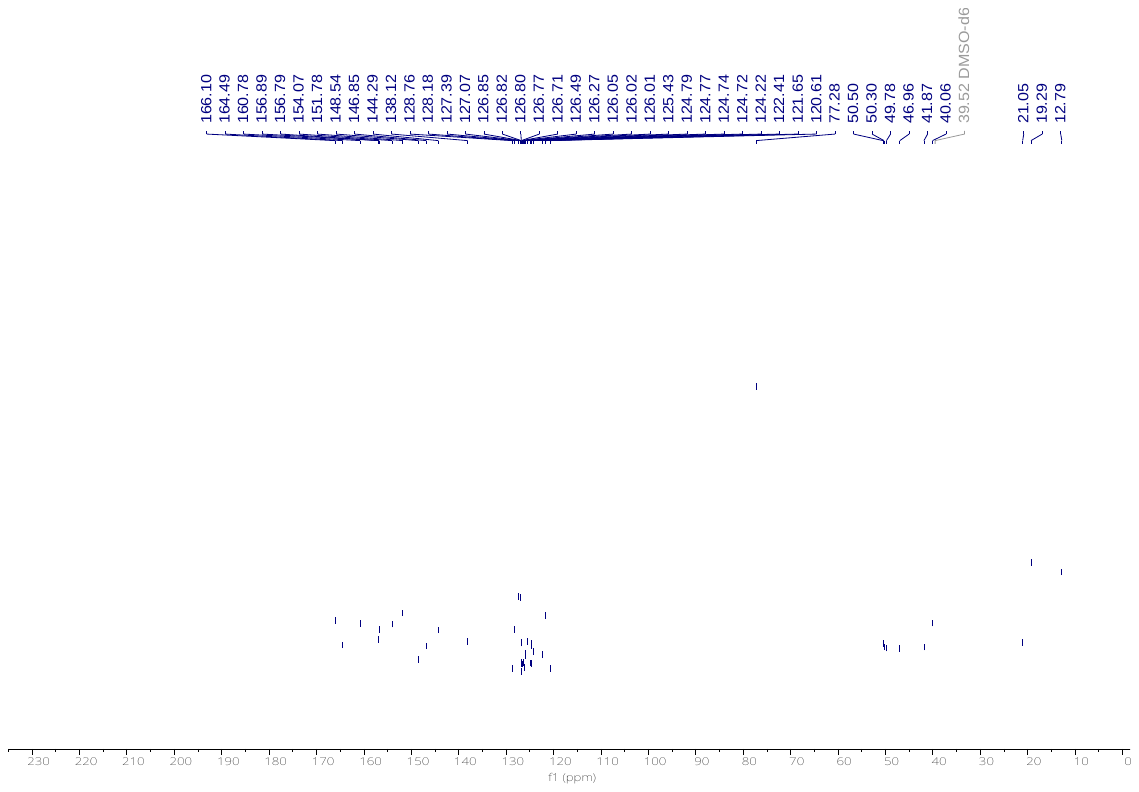

**^19^F NMR of GBA-007**

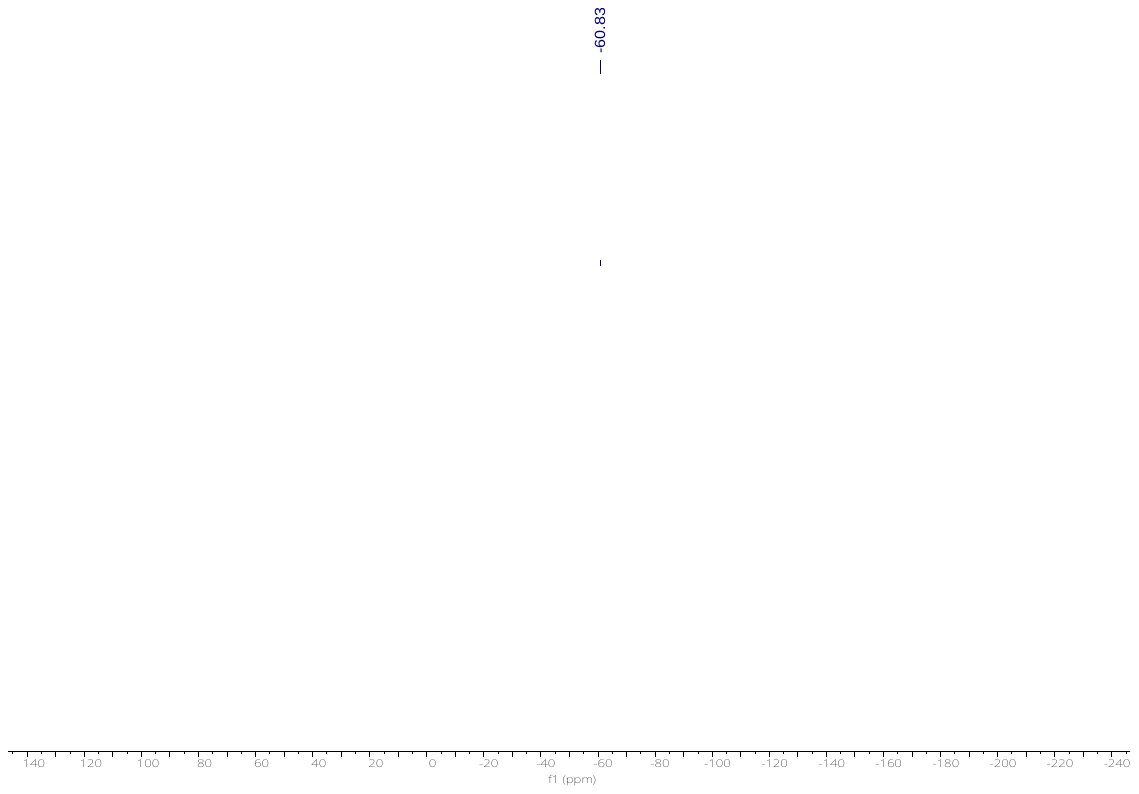

**HRMS of GBA-007**

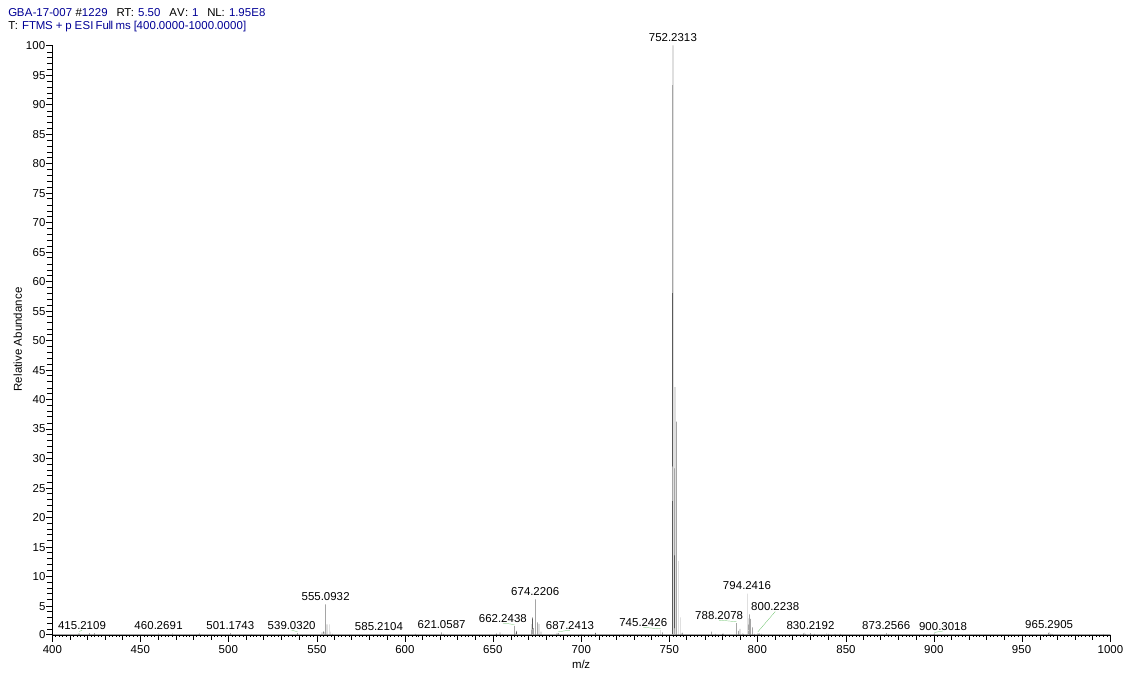

**HPLC of GBA-007**

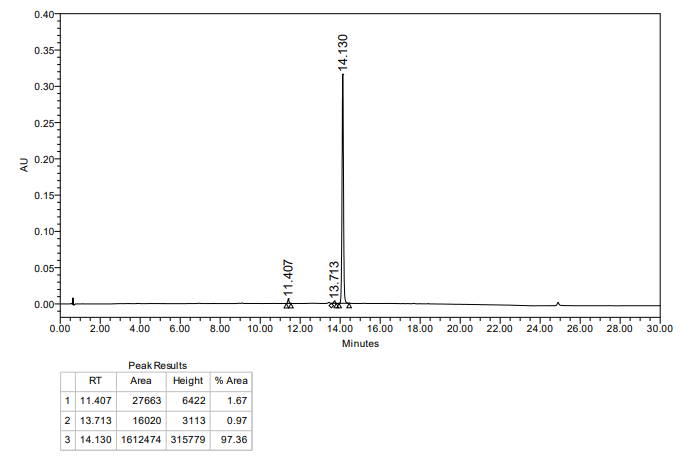

**^1^H NMR of GBA-008**

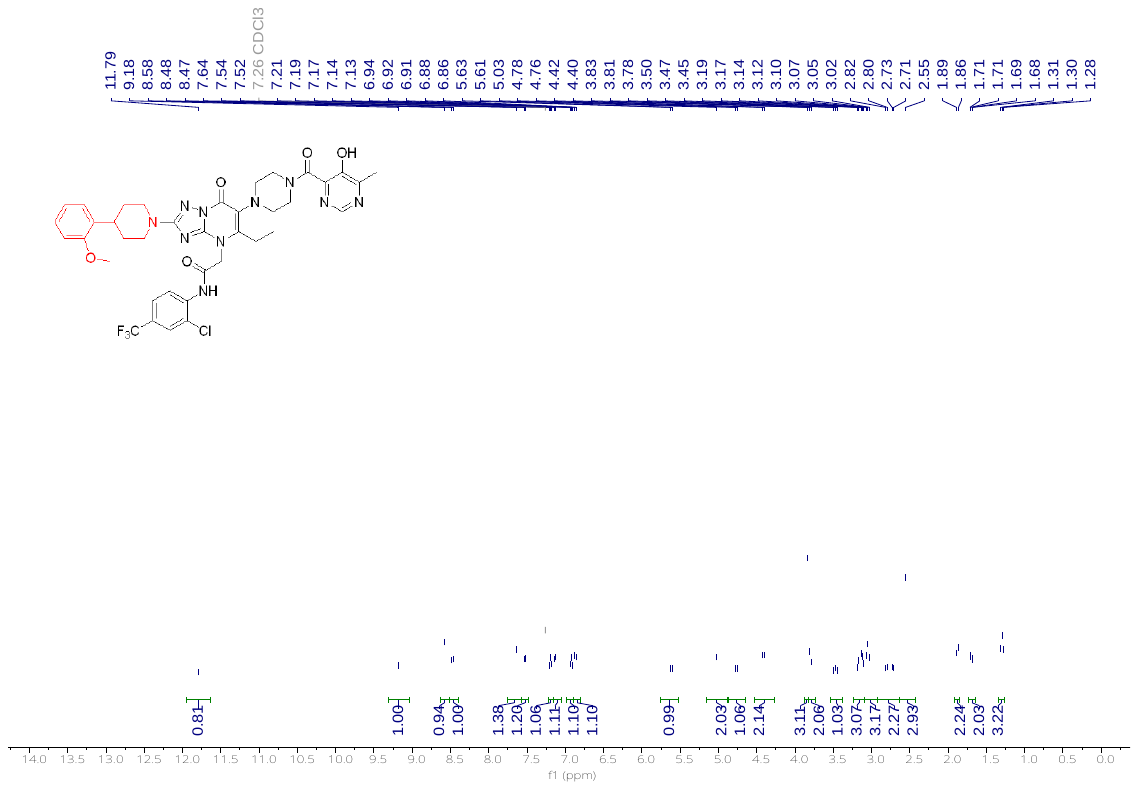

**^13^C NMR of GBA-008**

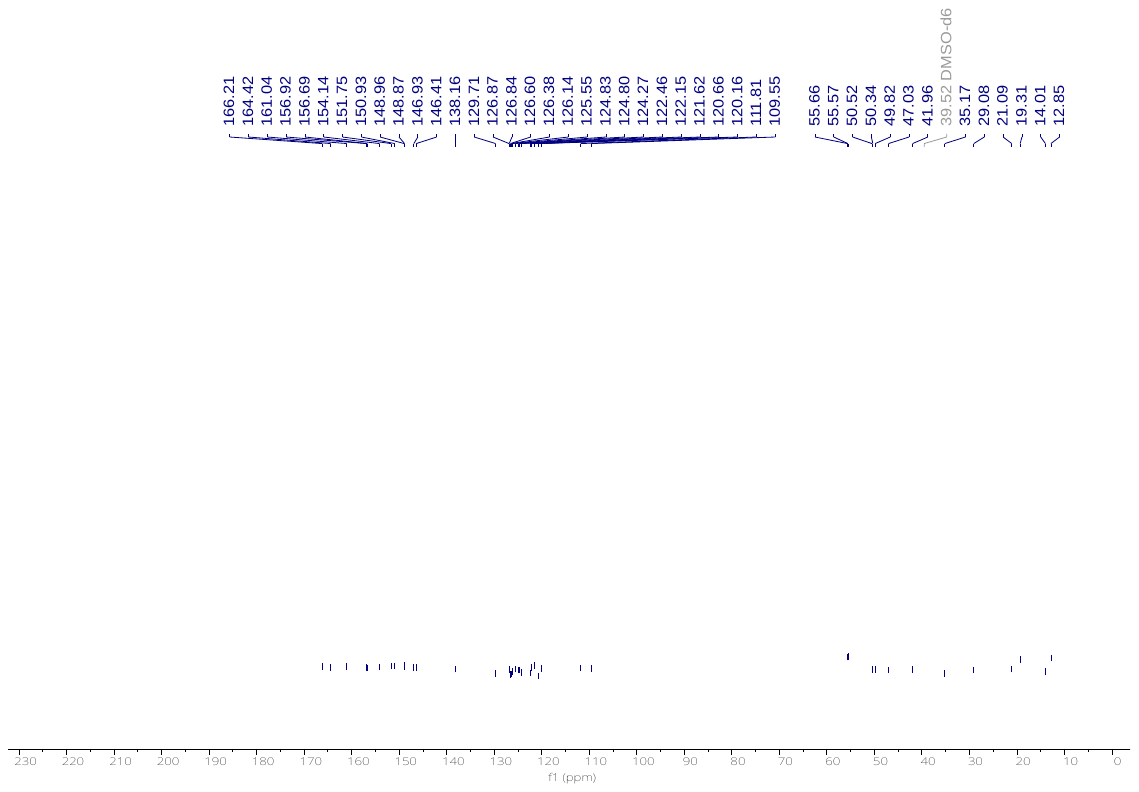

**^19^F NMR of GBA-008**

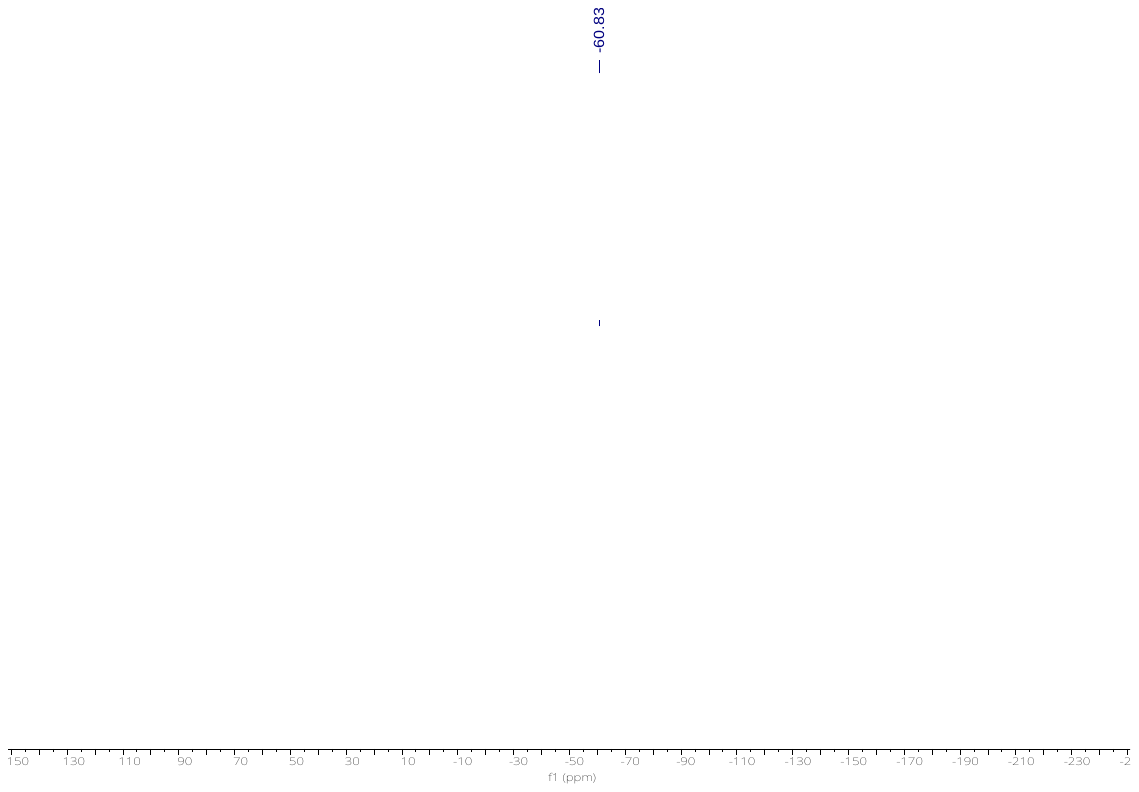

**HRMS of GBA-008**

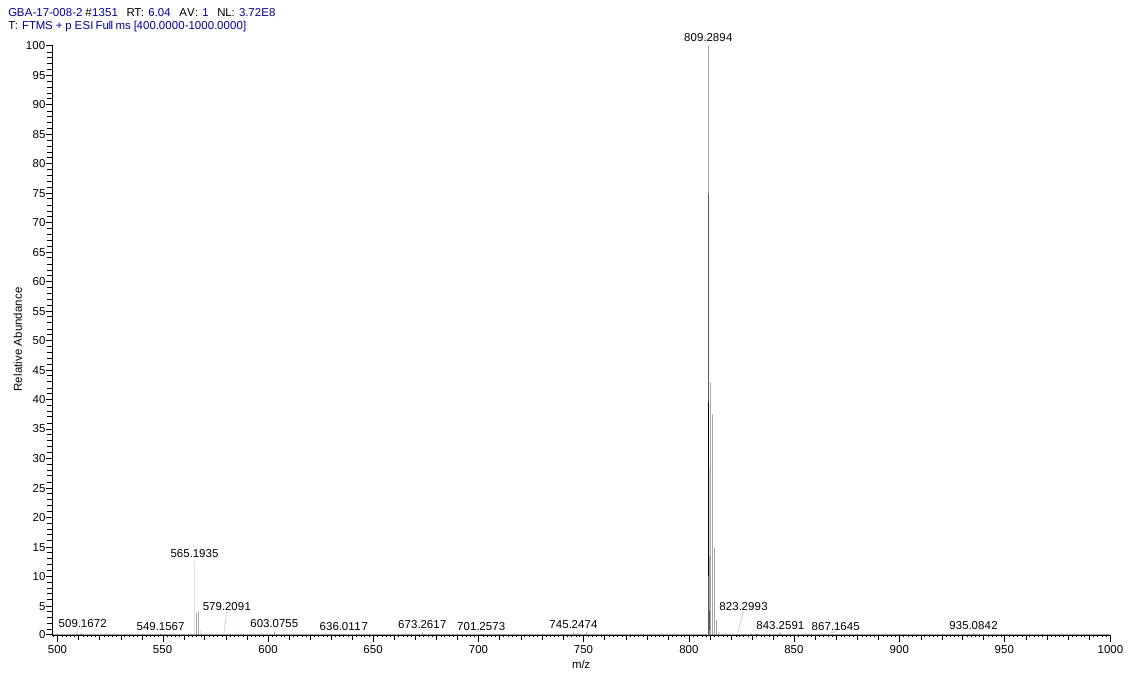

**HPLC of GBA-008**

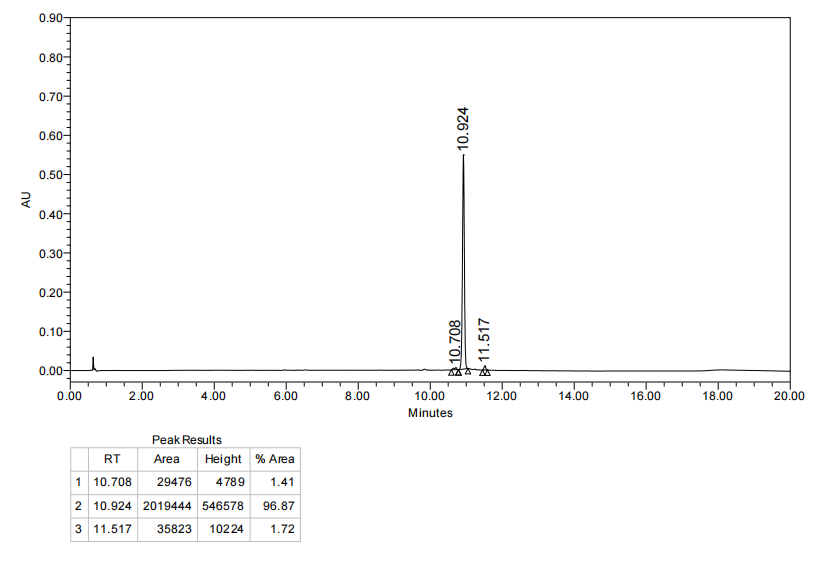

**^1^H NMR of GBA-029**

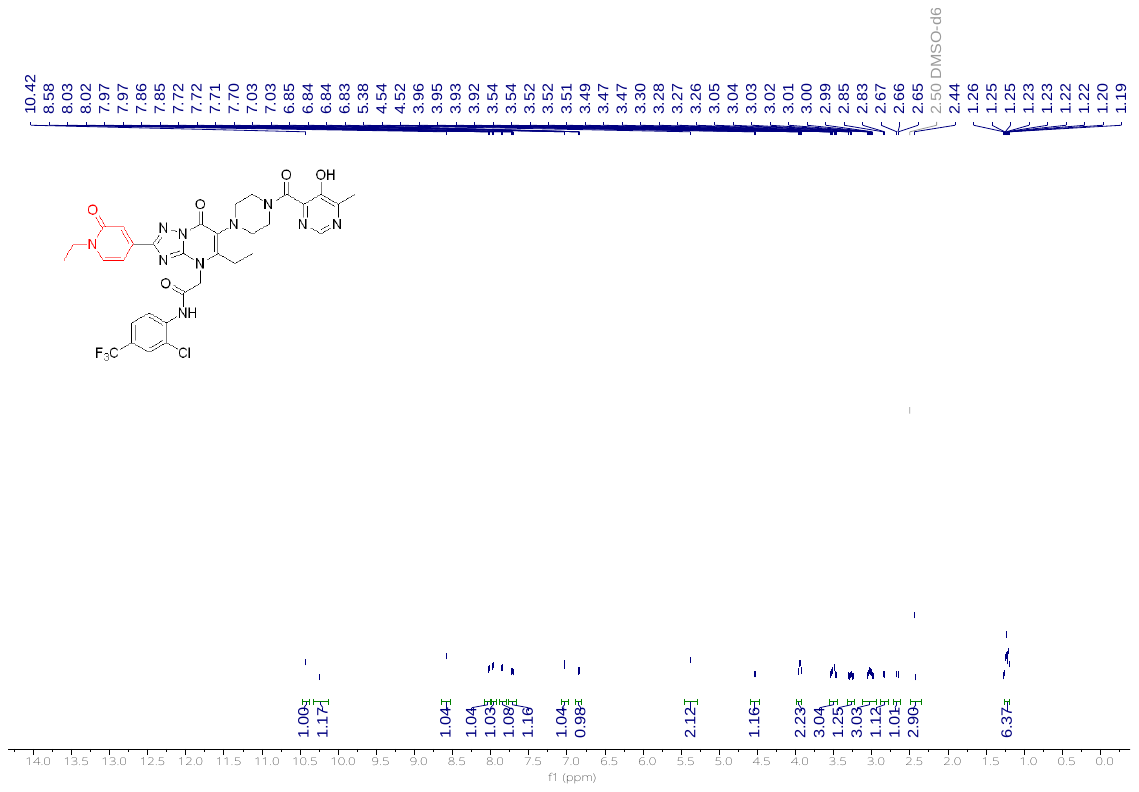

**^13^C NMR of GBA-029**

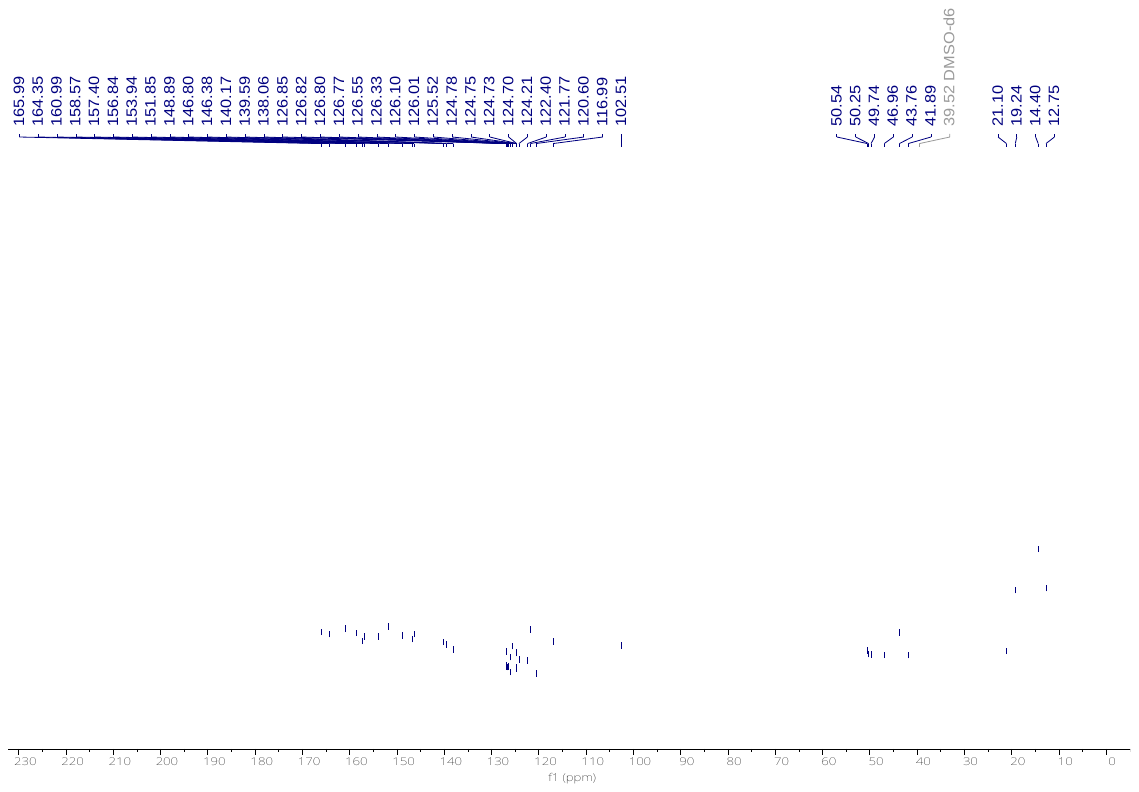

**^19^F NMR of GBA-029**

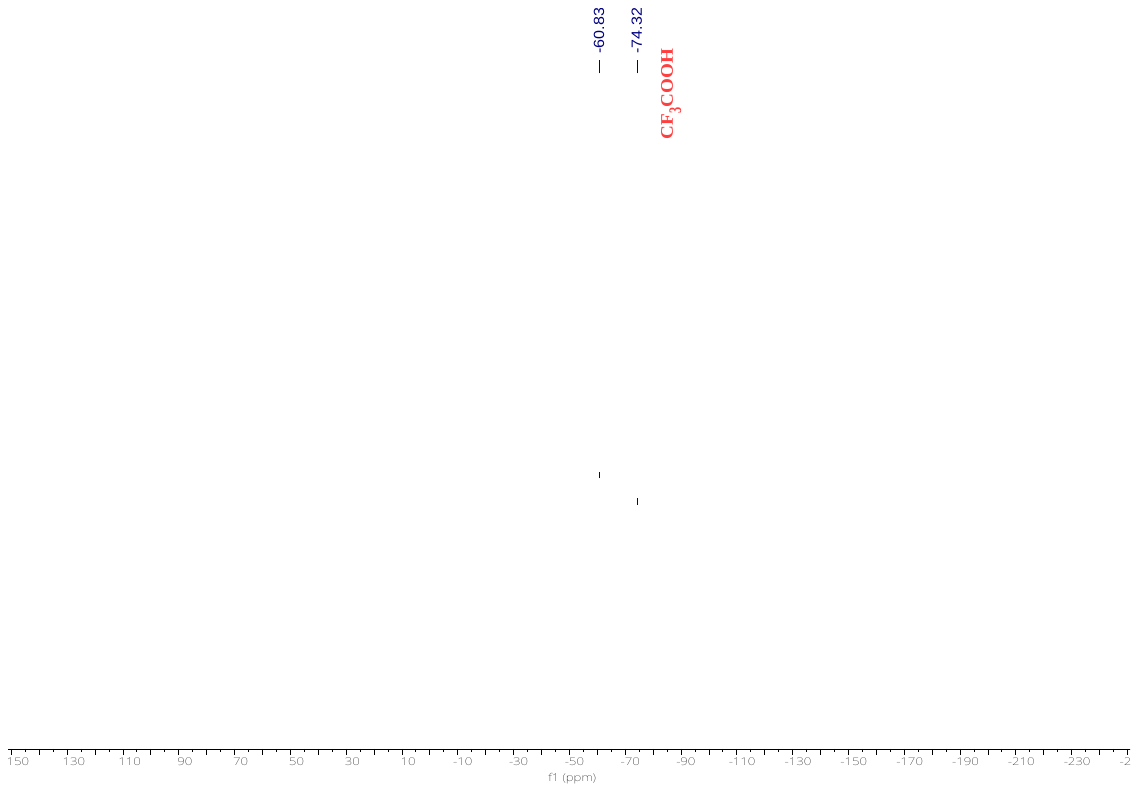

**HRMS of GBA-029**

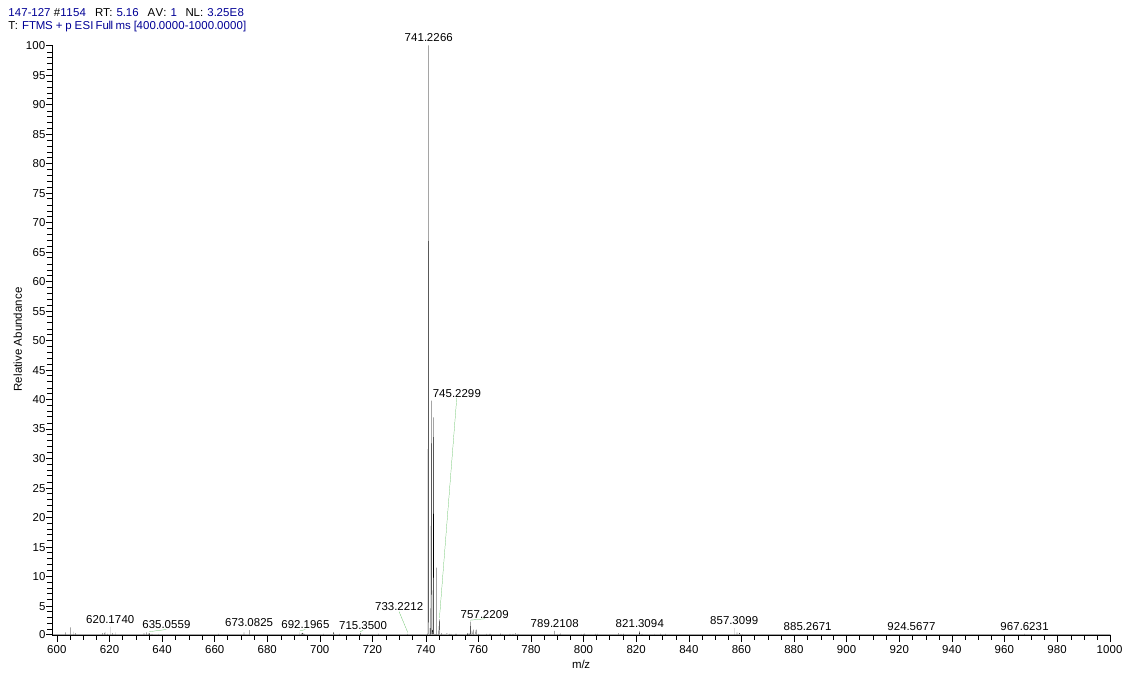

**HPLC of GBA-029**

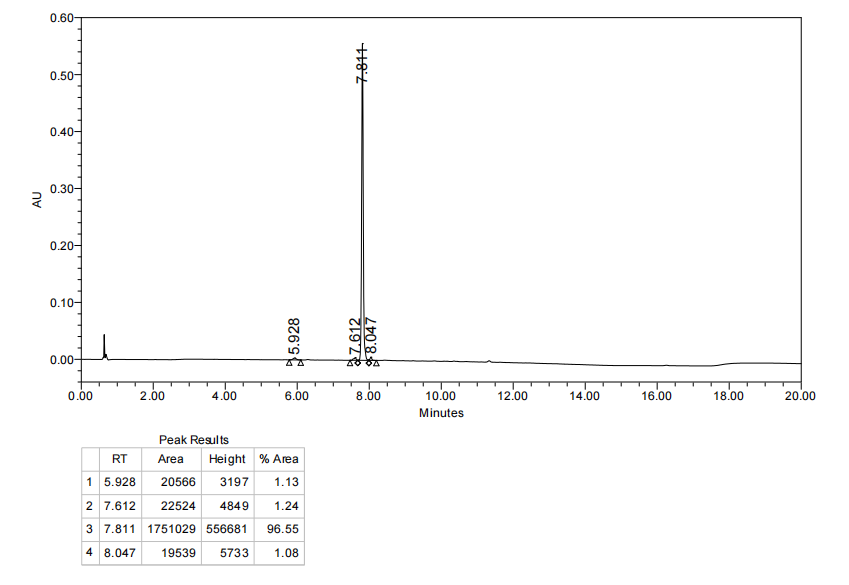

**Uncropped image corresponding to Figure 2a.**

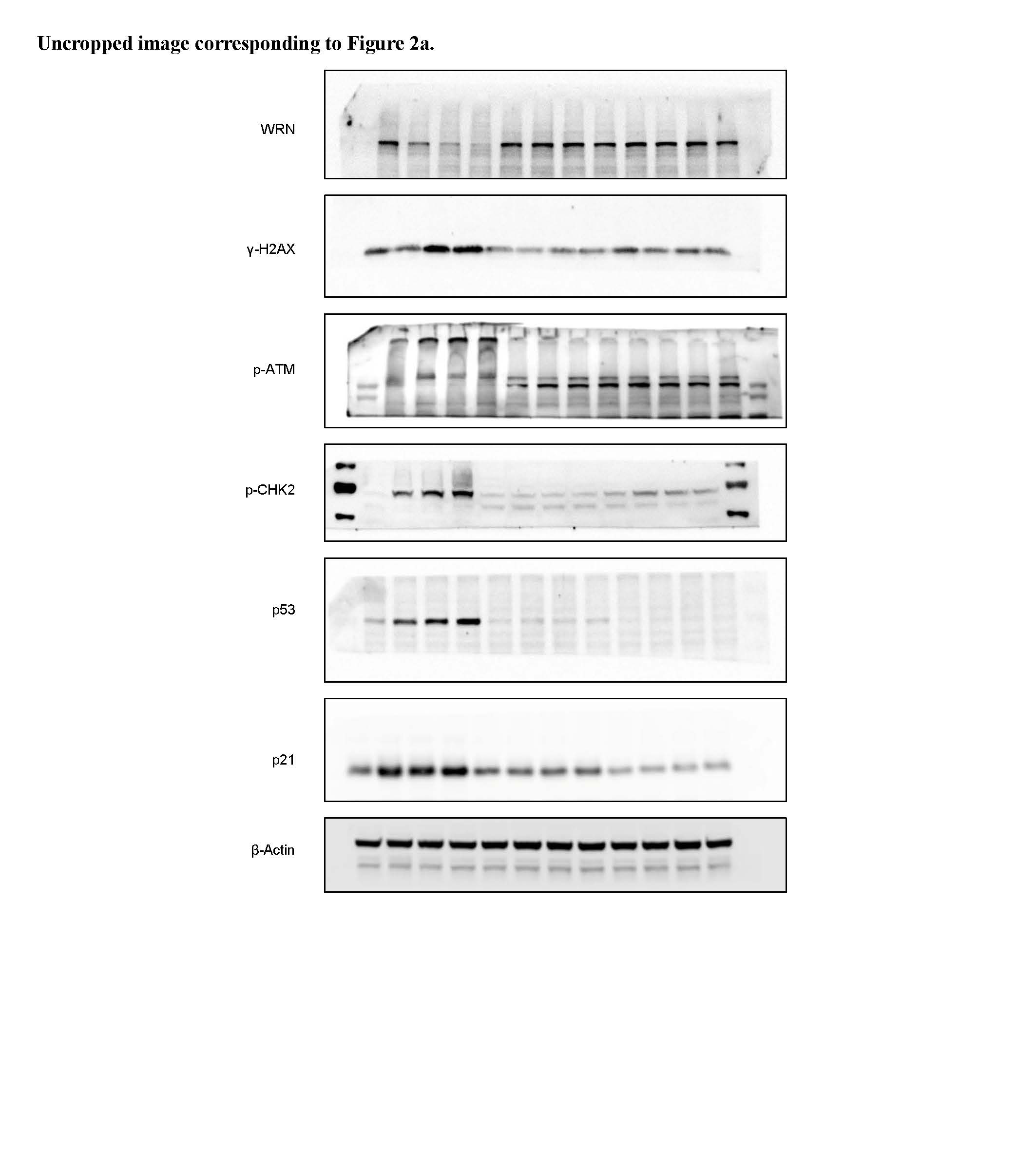

**Uncropped image corresponding to Figure 2b.**

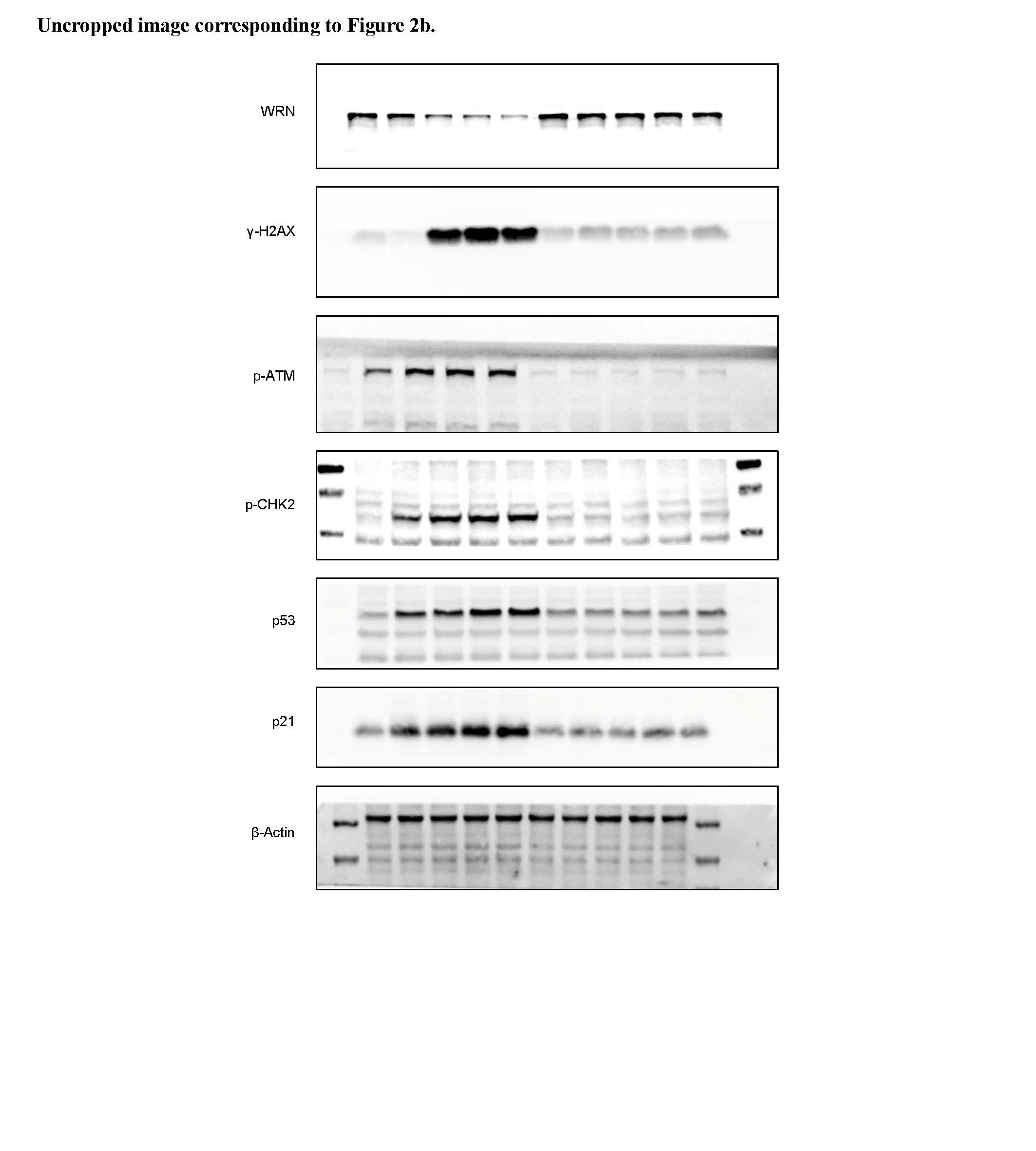

**Uncropped image corresponding to Figure 4g.**

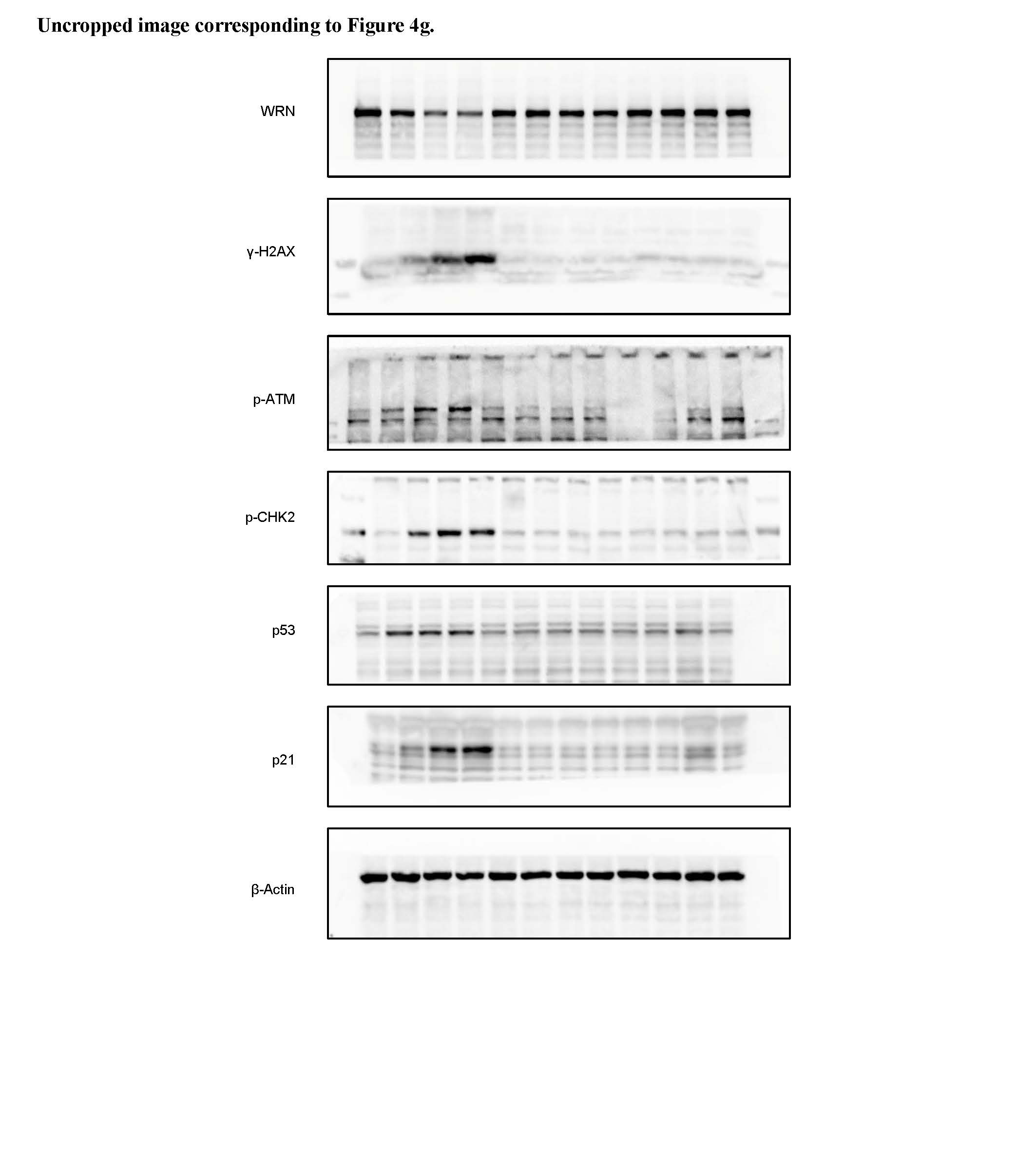

**Uncropped image corresponding to Figure 4h.**

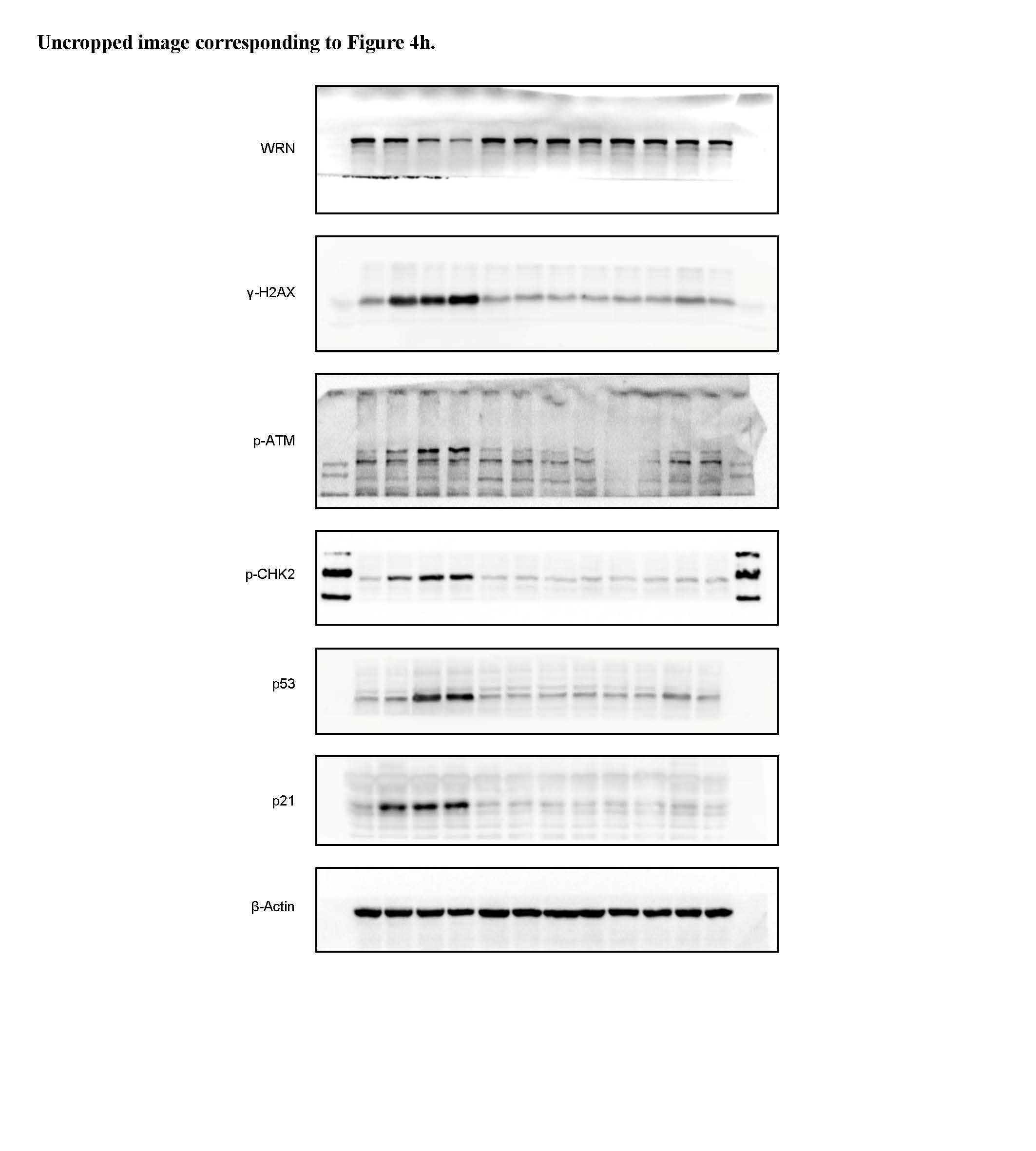

**Uncropped image corresponding to Figure 4i.**

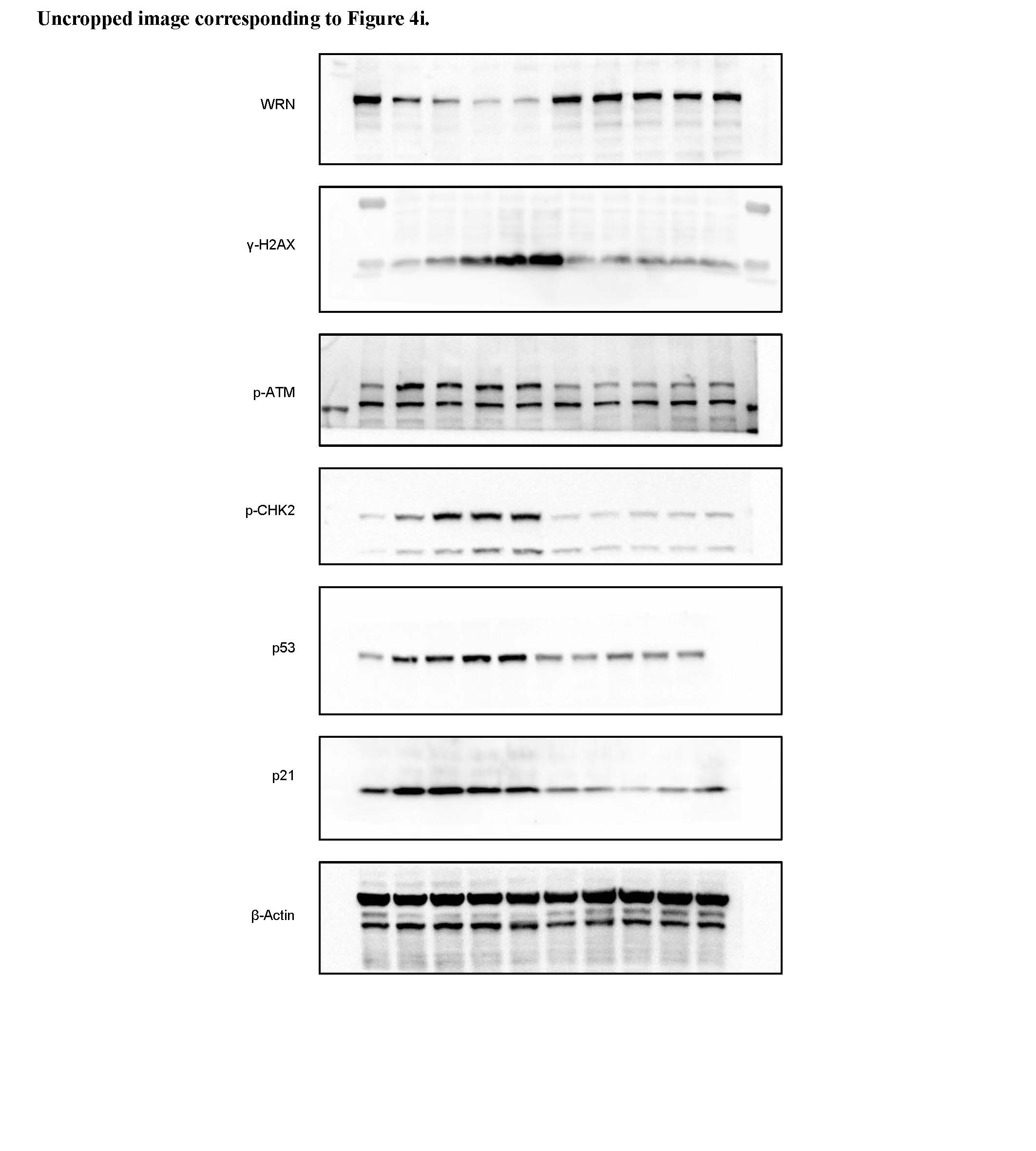

**Uncropped image corresponding to Figure 4j.**

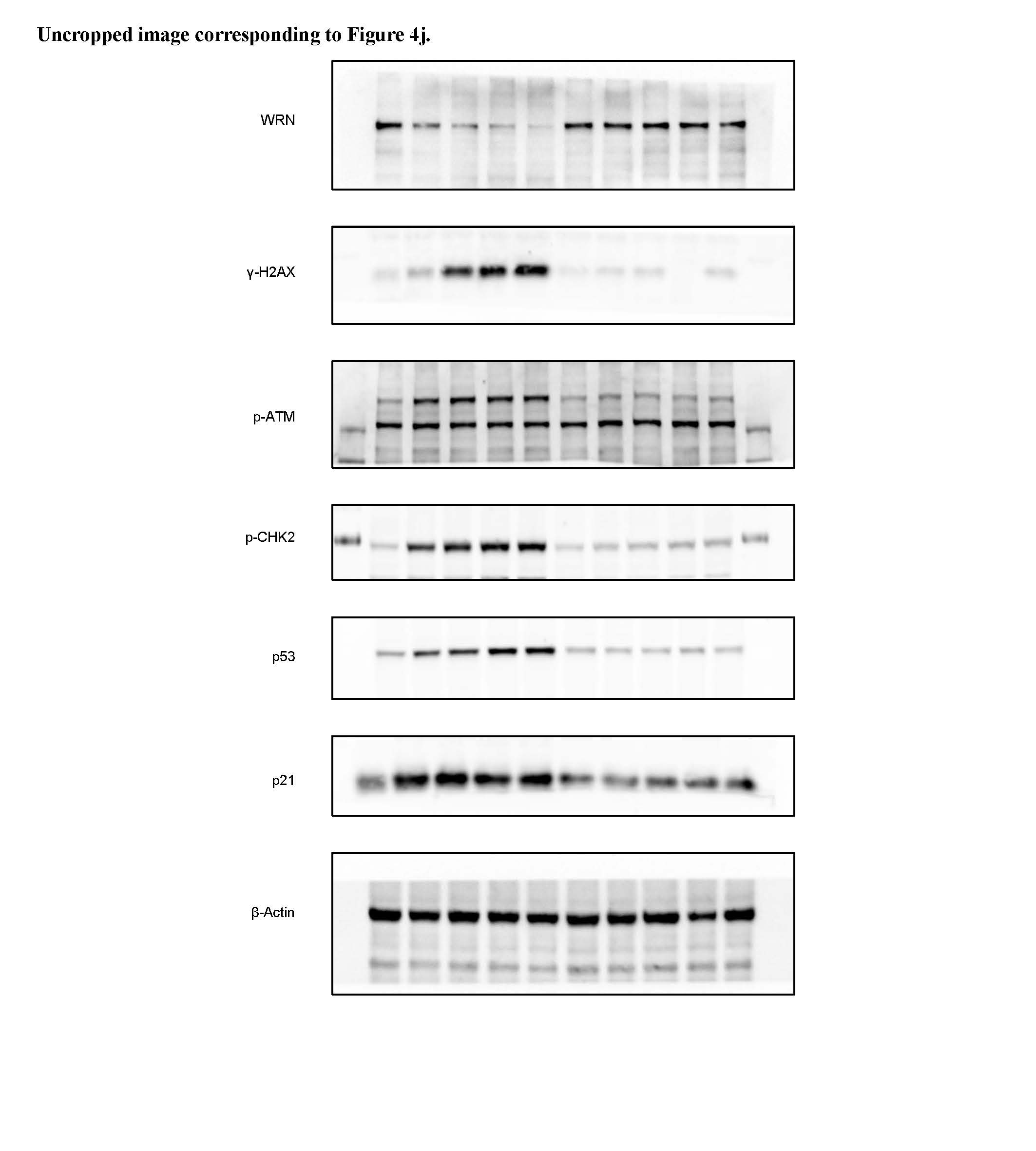

**Uncropped image corresponding to Figure 5g.**

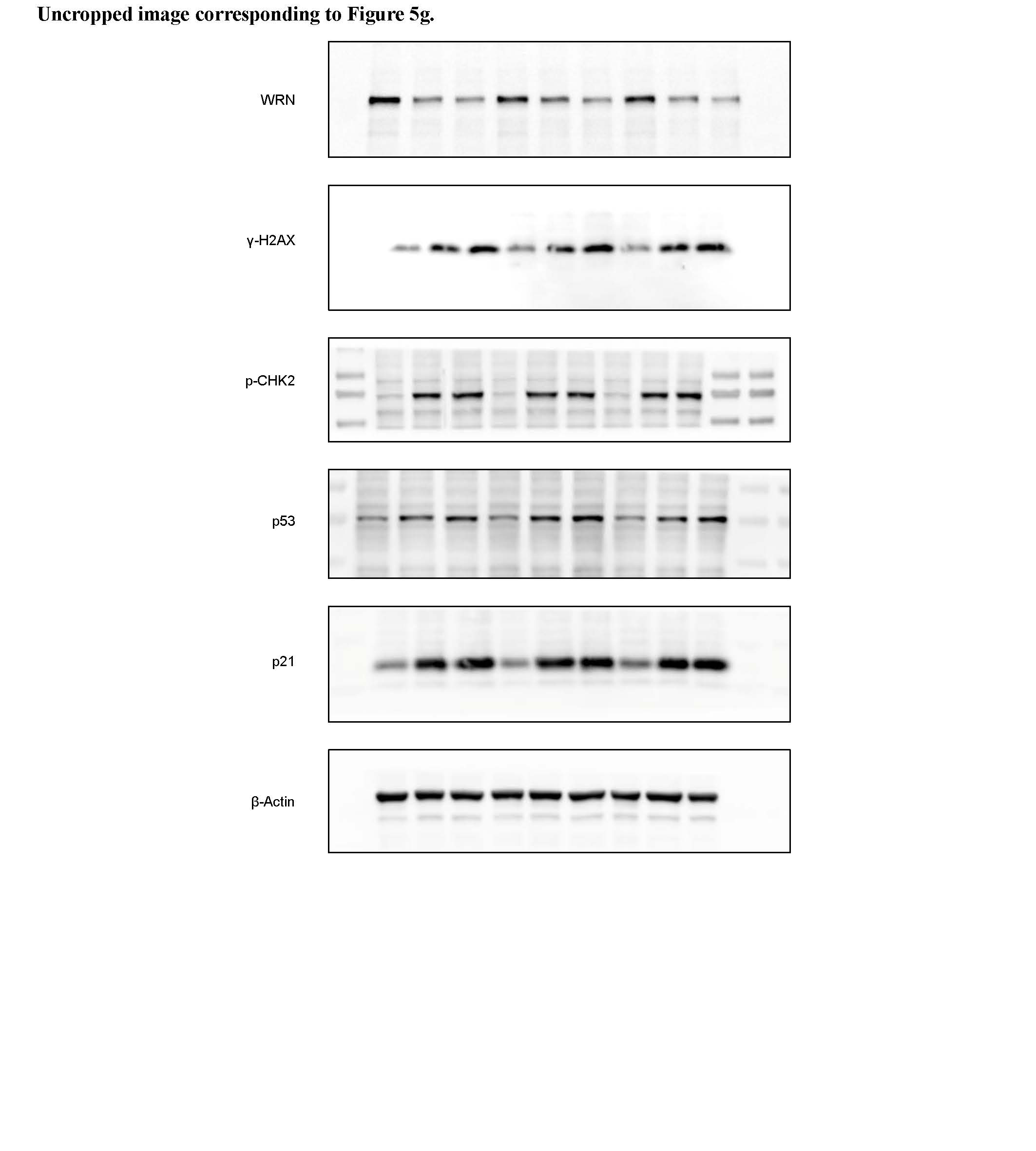

**Uncropped image corresponding to Figure 5h.**

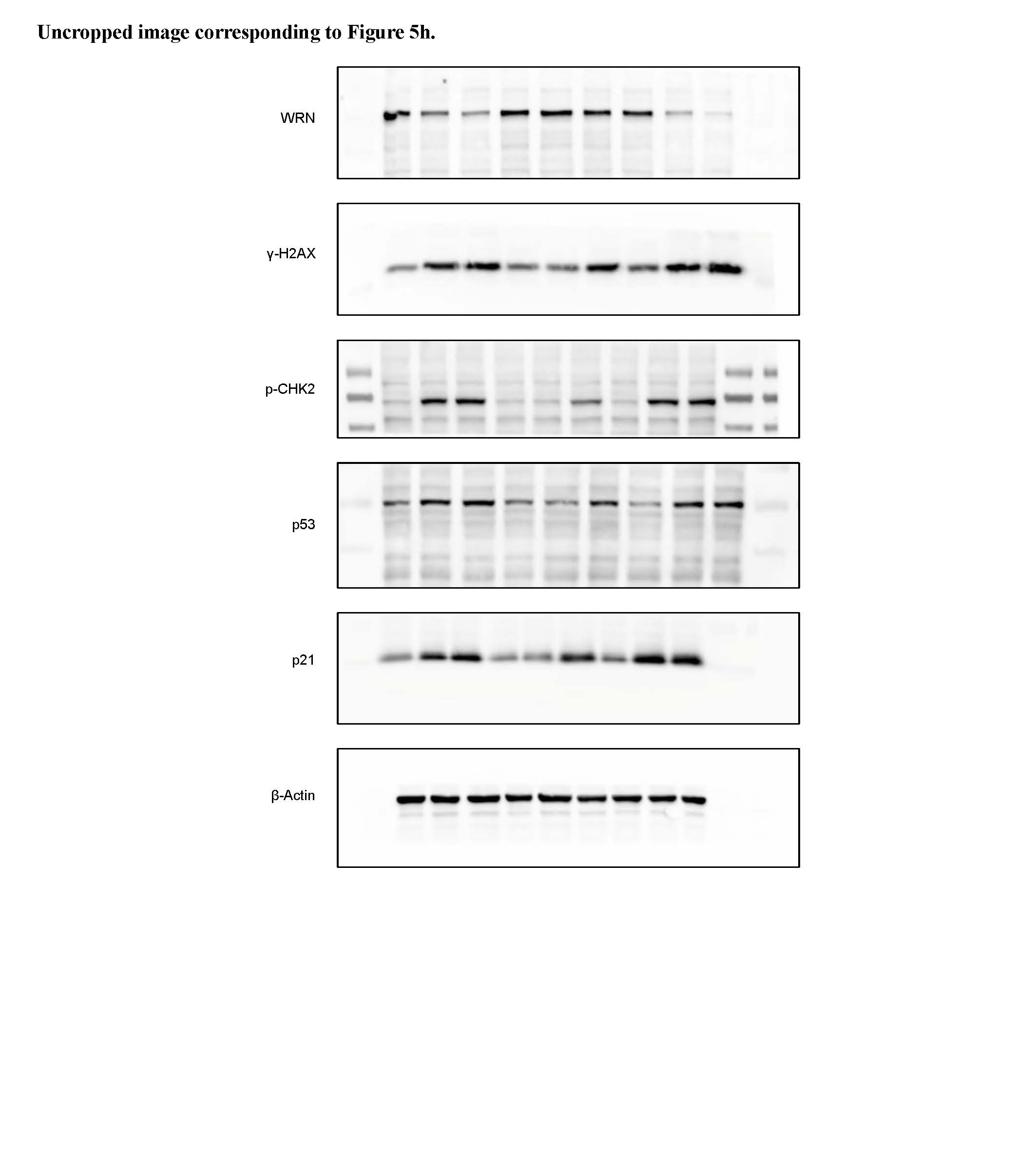

**Uncropped image corresponding to Figure 5i.**

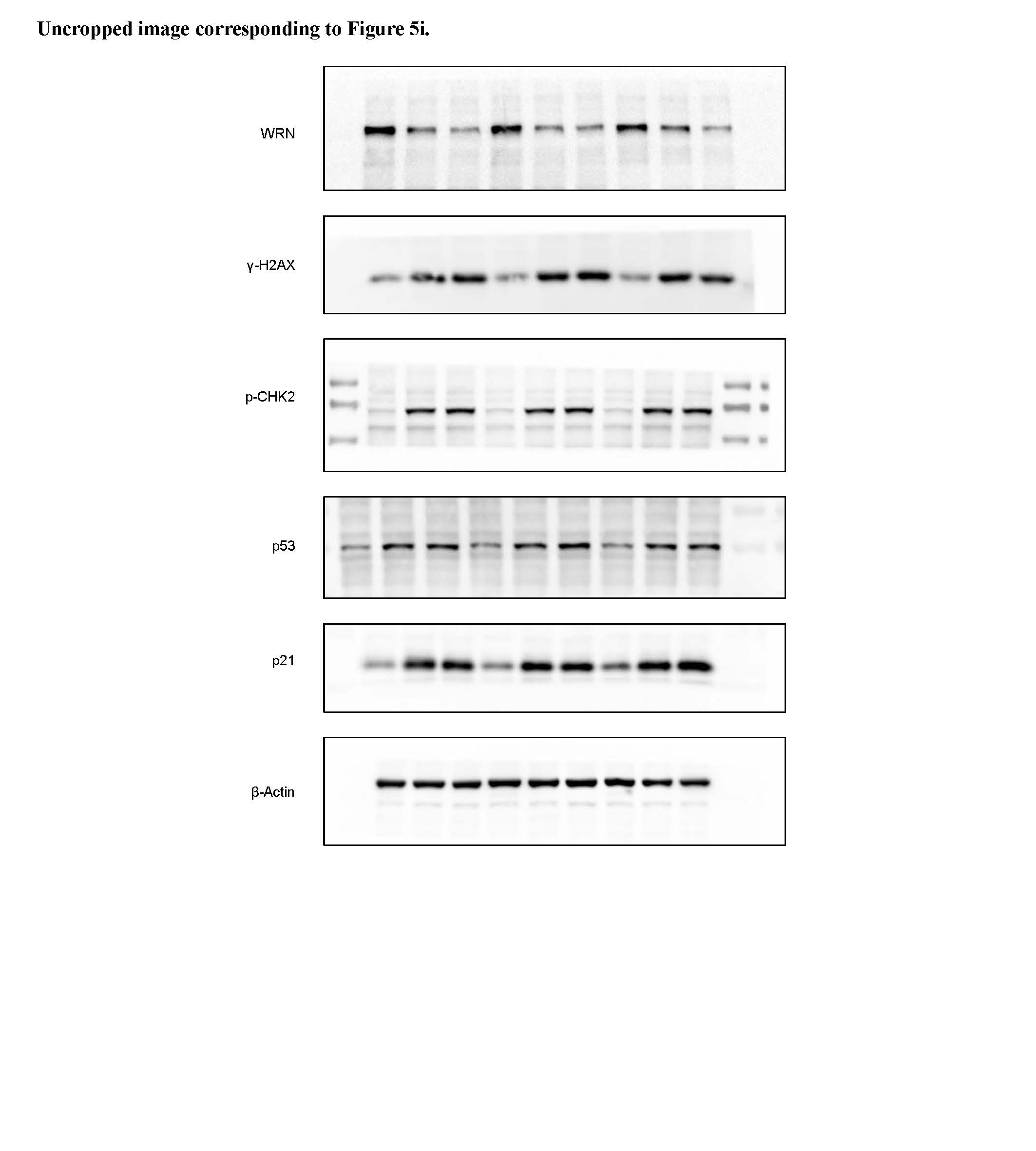

**Uncropped image corresponding to Figure 6j.**

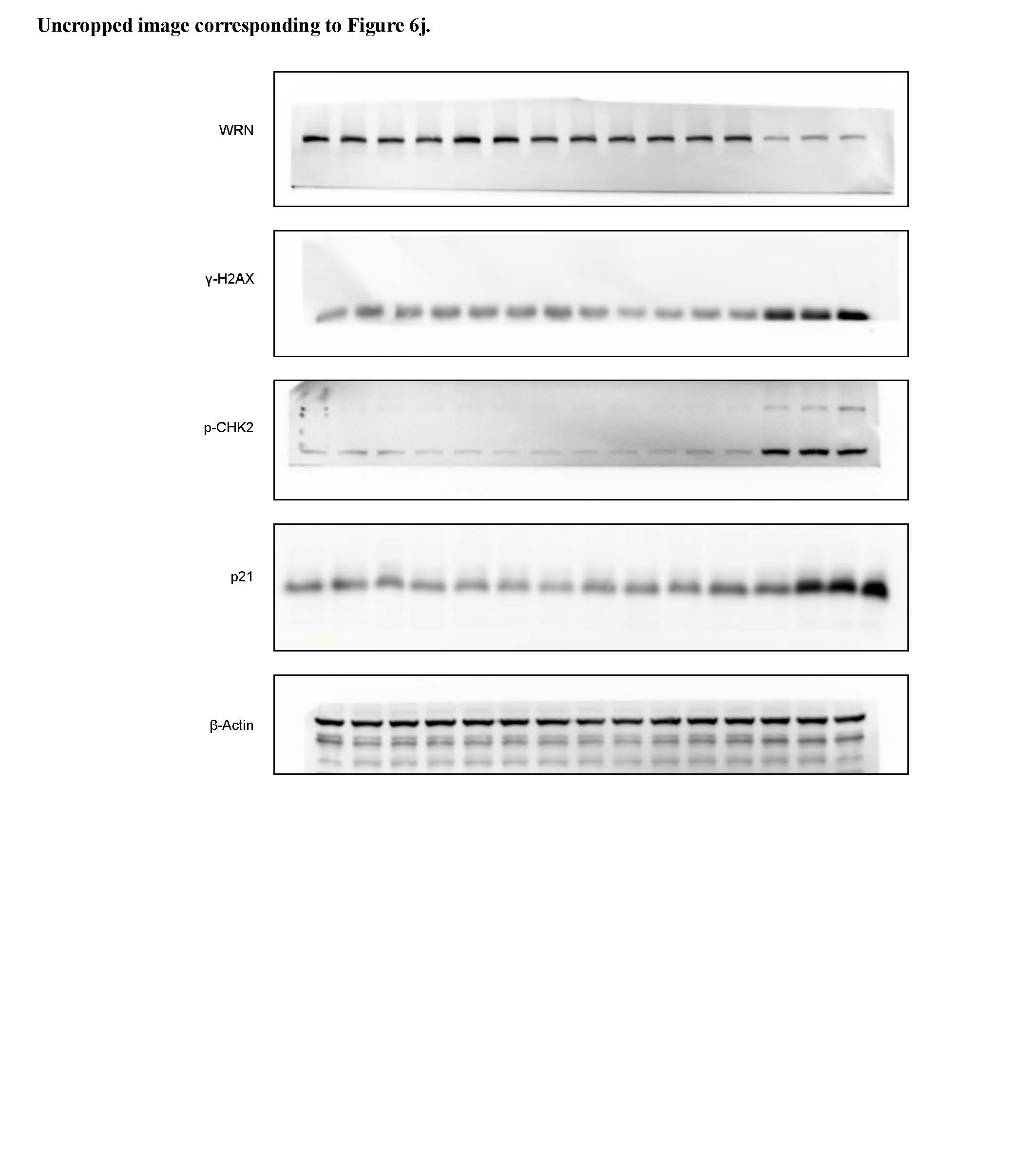

**Uncropped image corresponding to Figure 8d.**

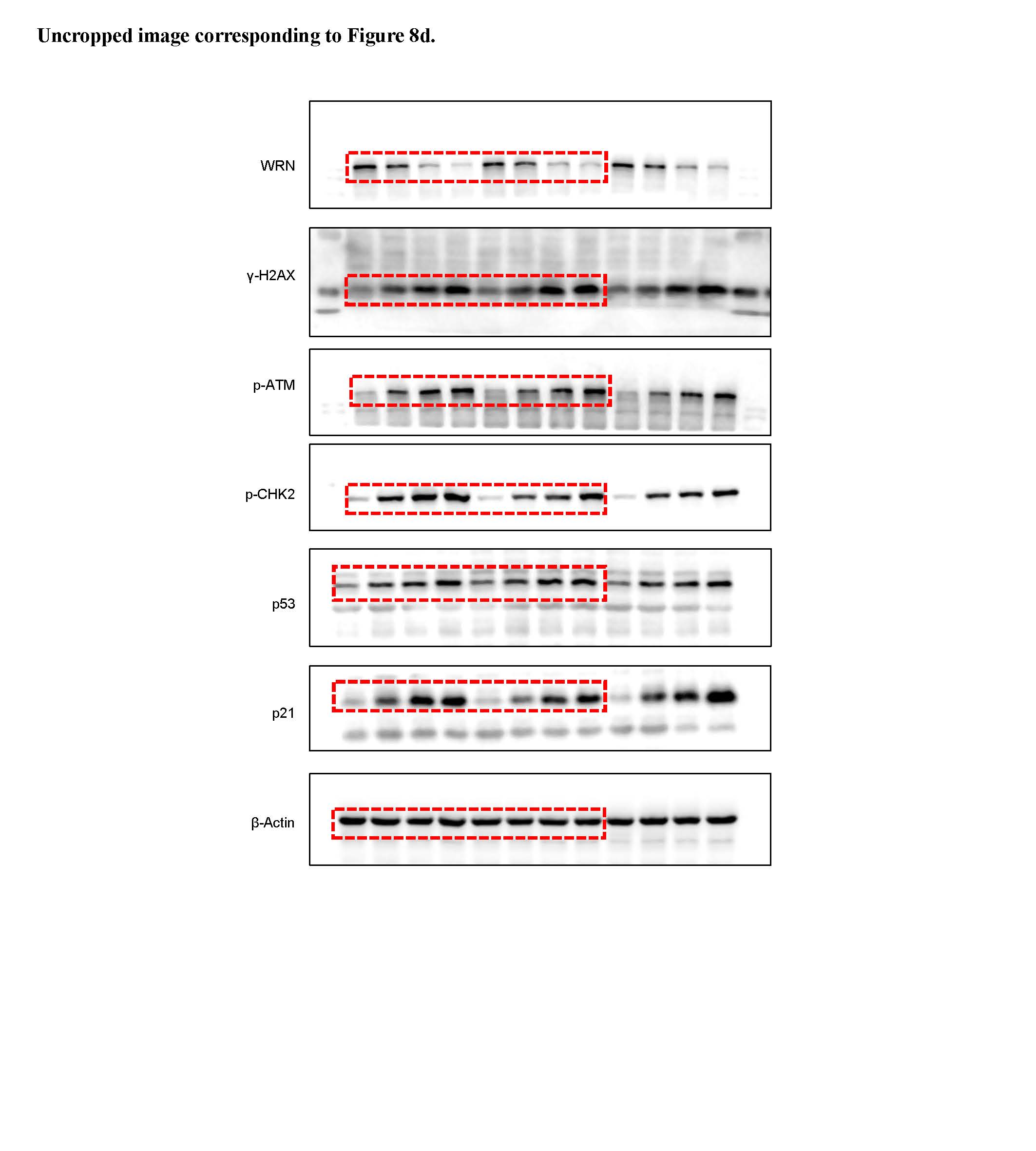

**Uncropped image corresponding to Figure 8i.**

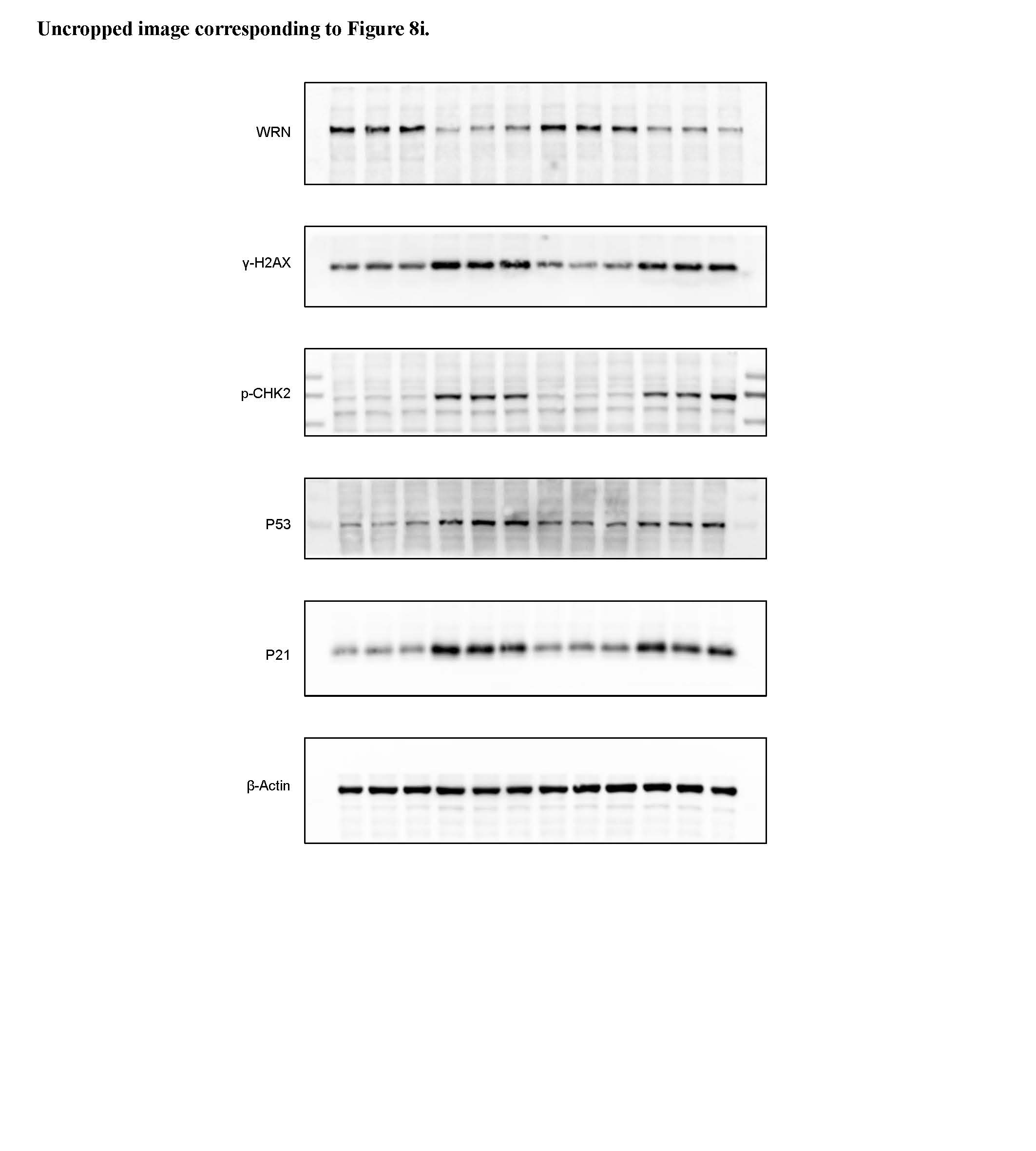

**Uncropped image corresponding to Supplementary Figure S1a.**

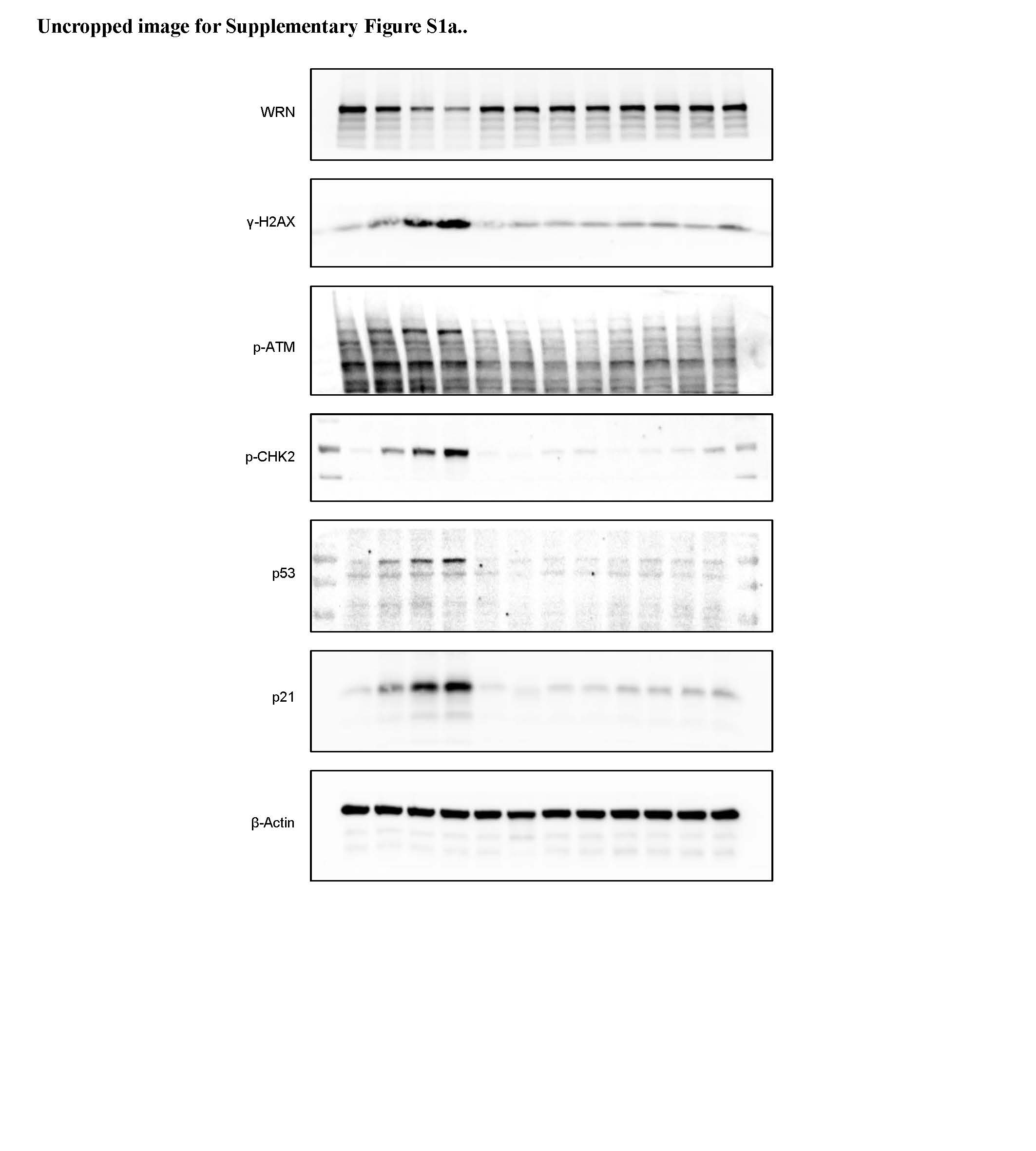

**Uncropped image corresponding to Supplementary Figure S1b.**

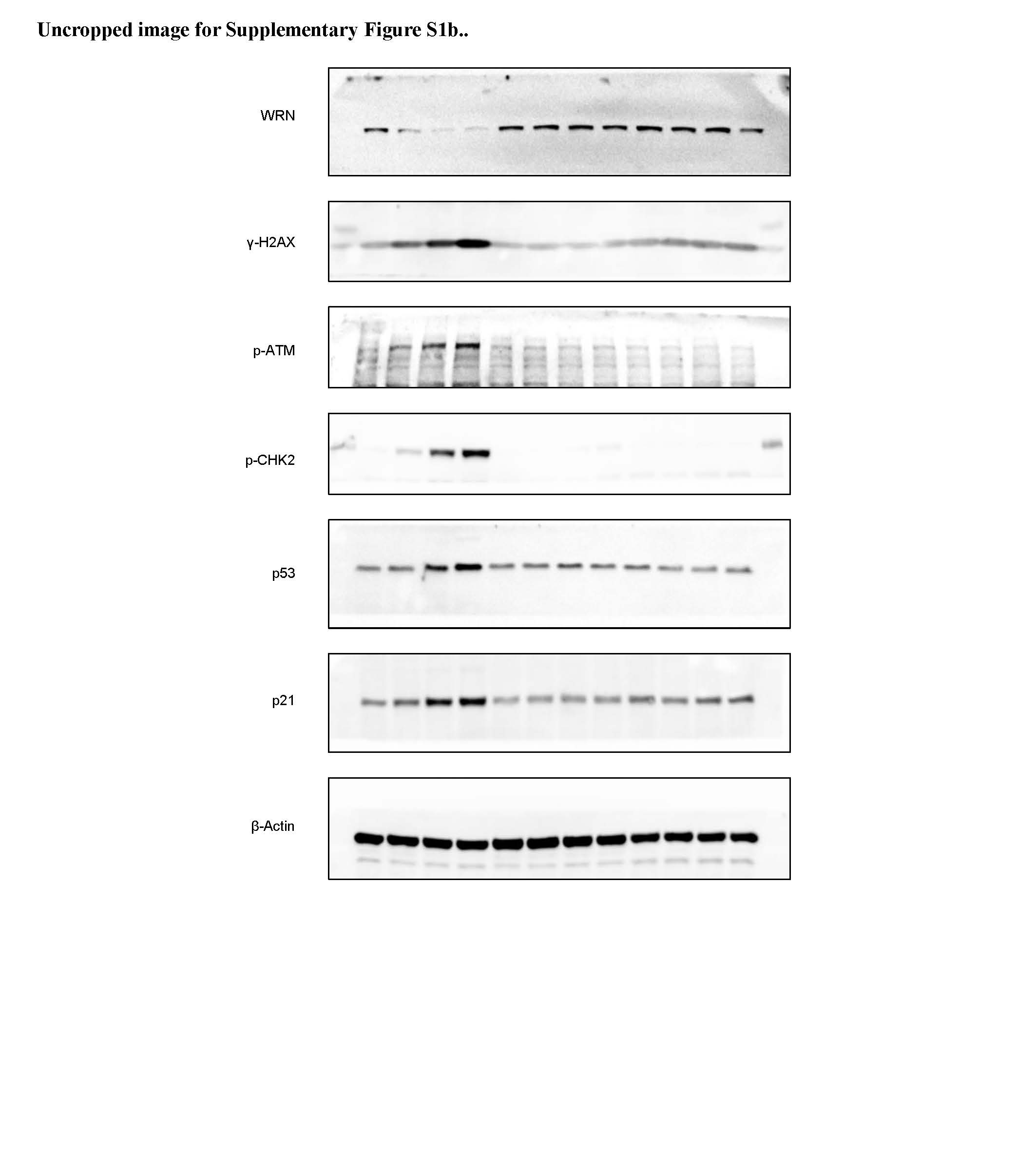

**Uncropped image corresponding to Supplementary Figure S1c.**

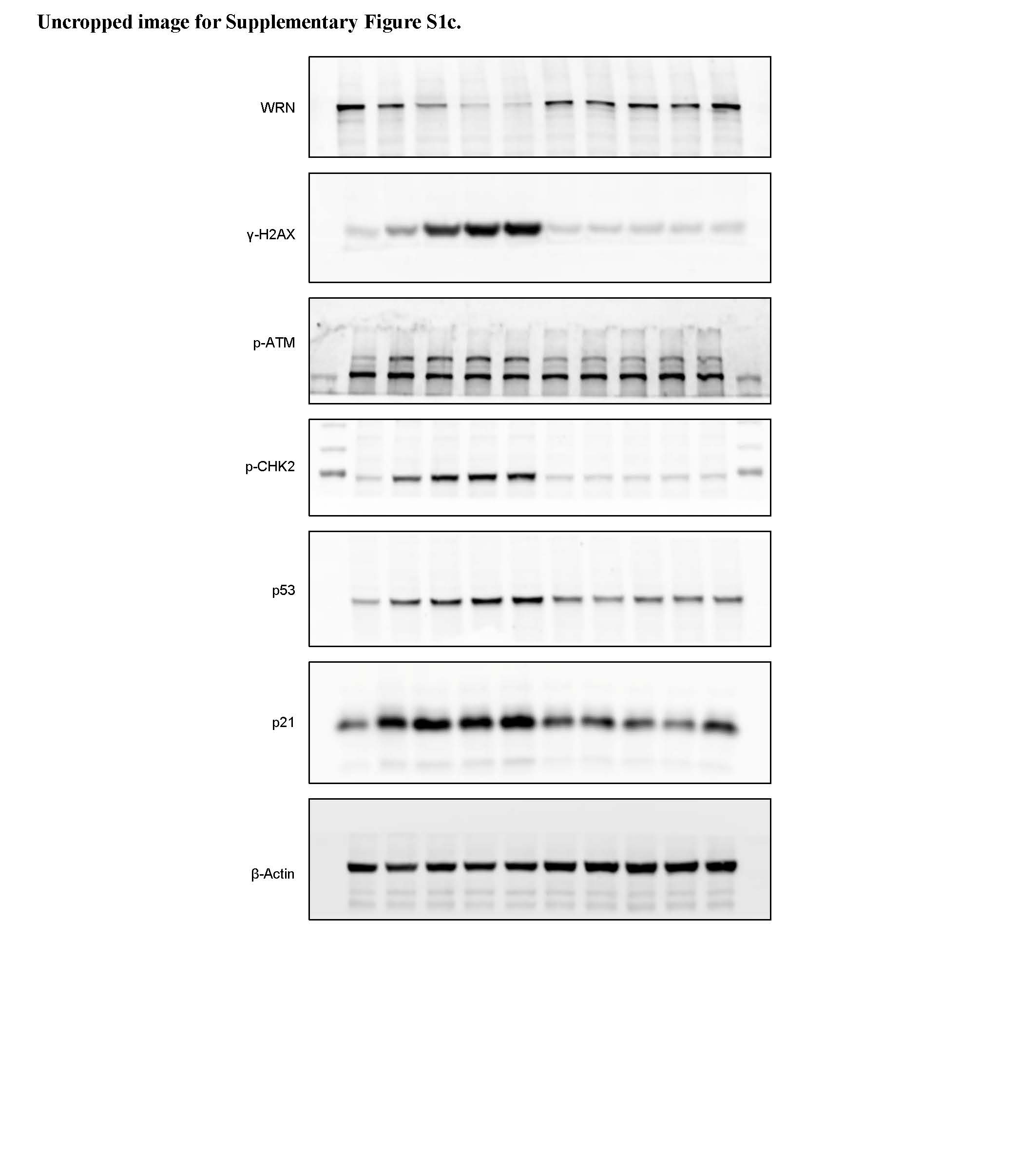

**Uncropped image corresponding to Supplementary Figure S1d.**

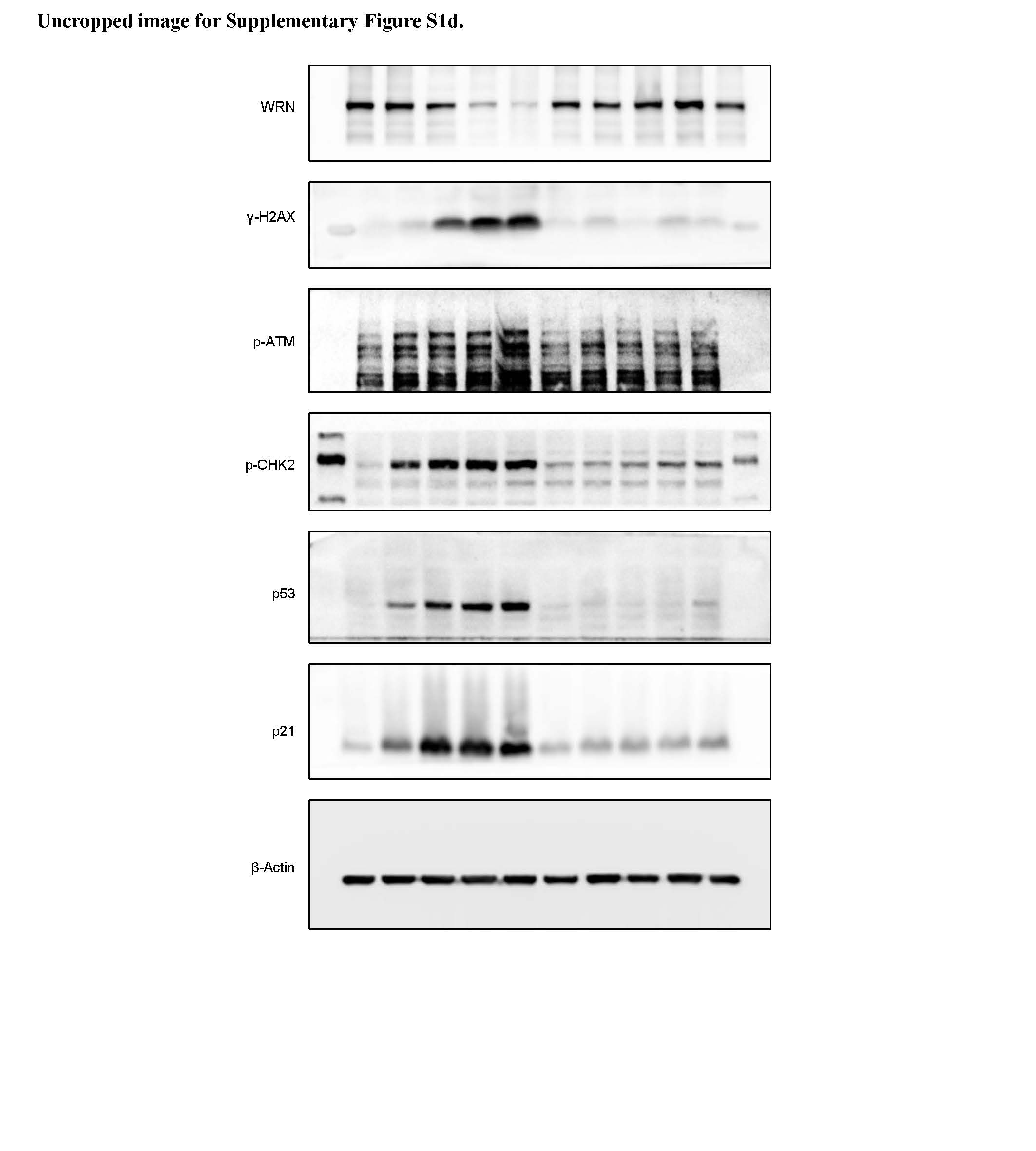

**Uncropped image corresponding to Supplementary Figure S1e.**

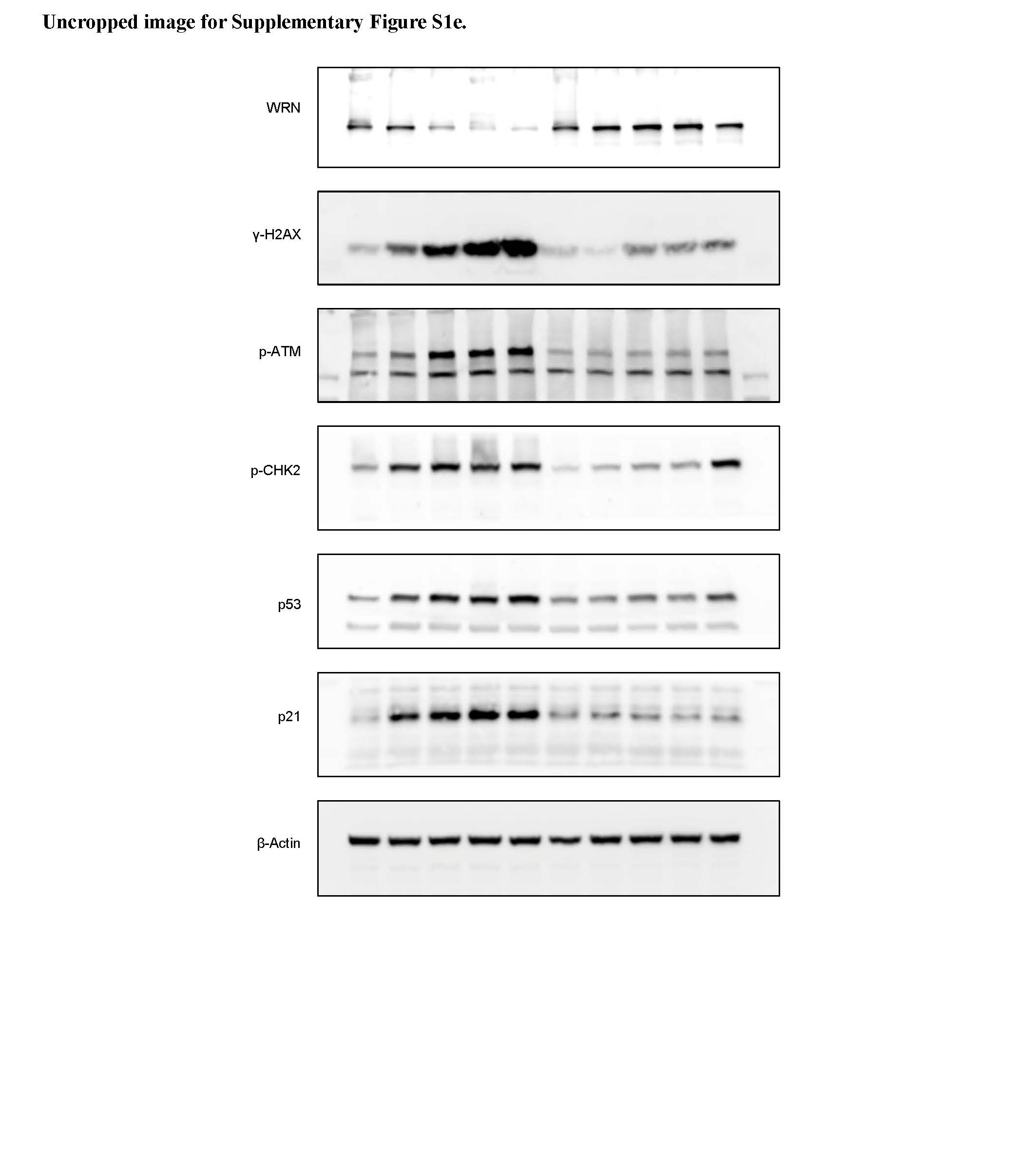

**Uncropped image corresponding to Supplementary Figure S1f.**

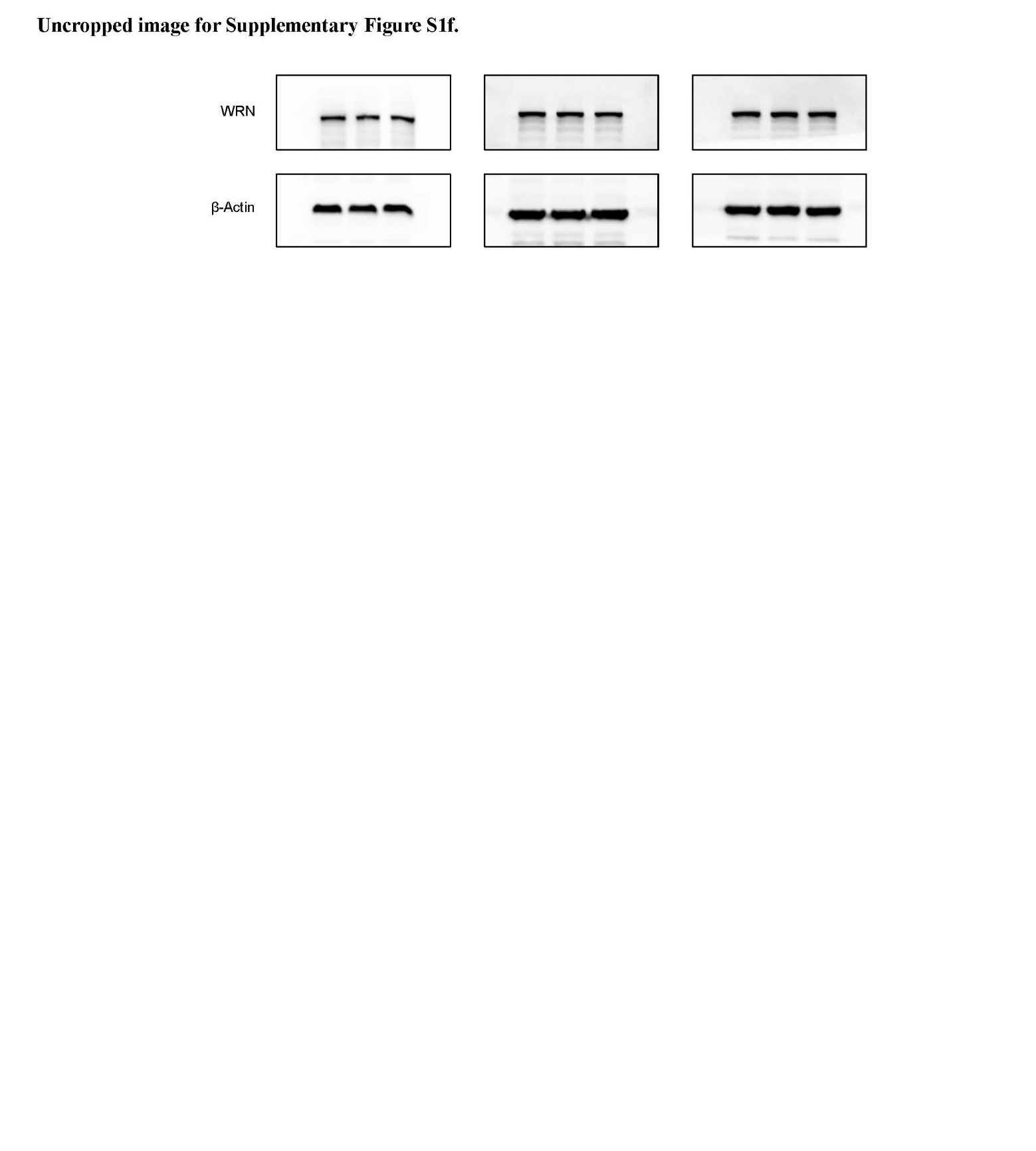

**Uncropped image corresponding to Supplementary Figure S1g.**

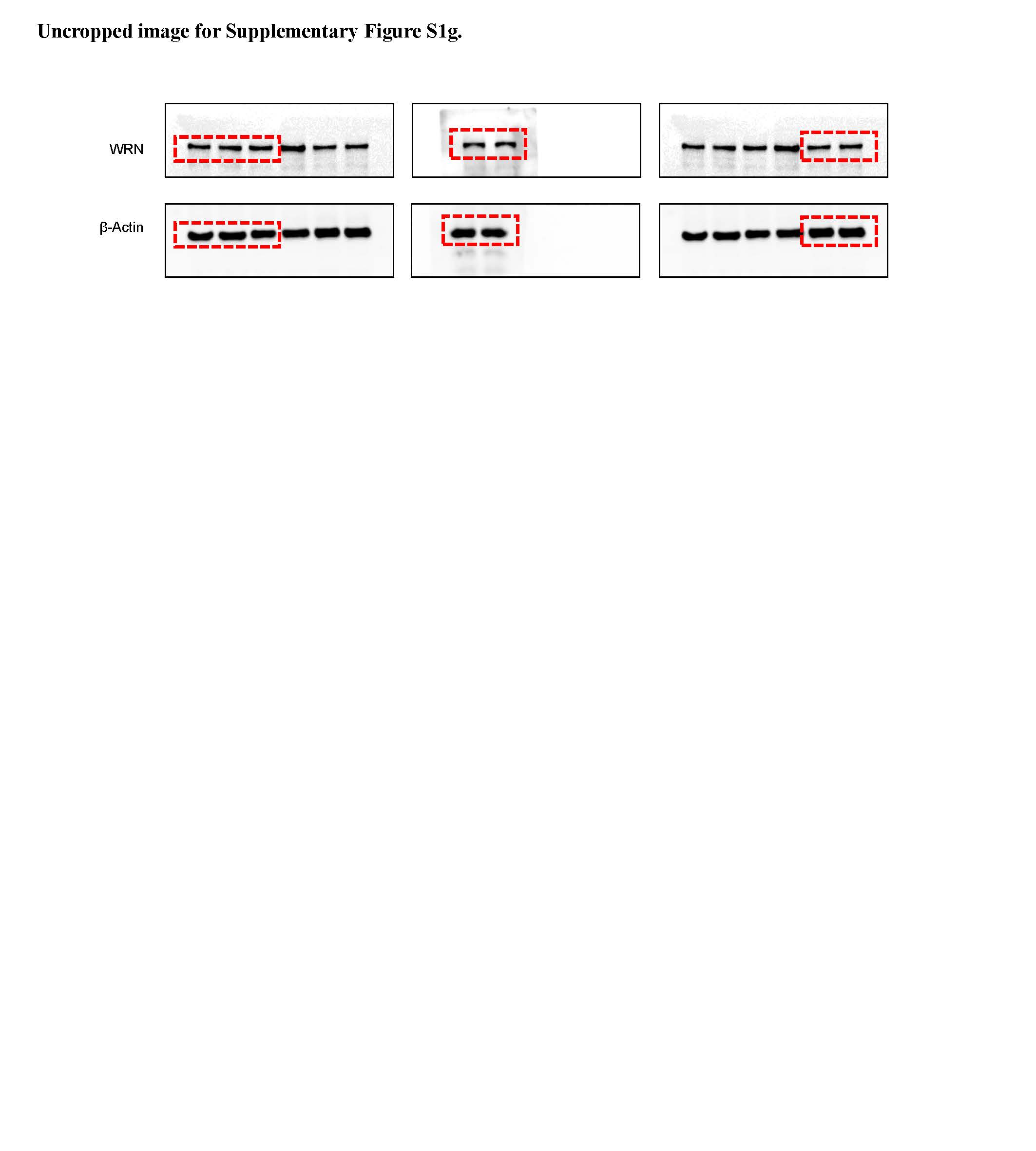

**Uncropped image corresponding to Supplementary Figure S2a.**

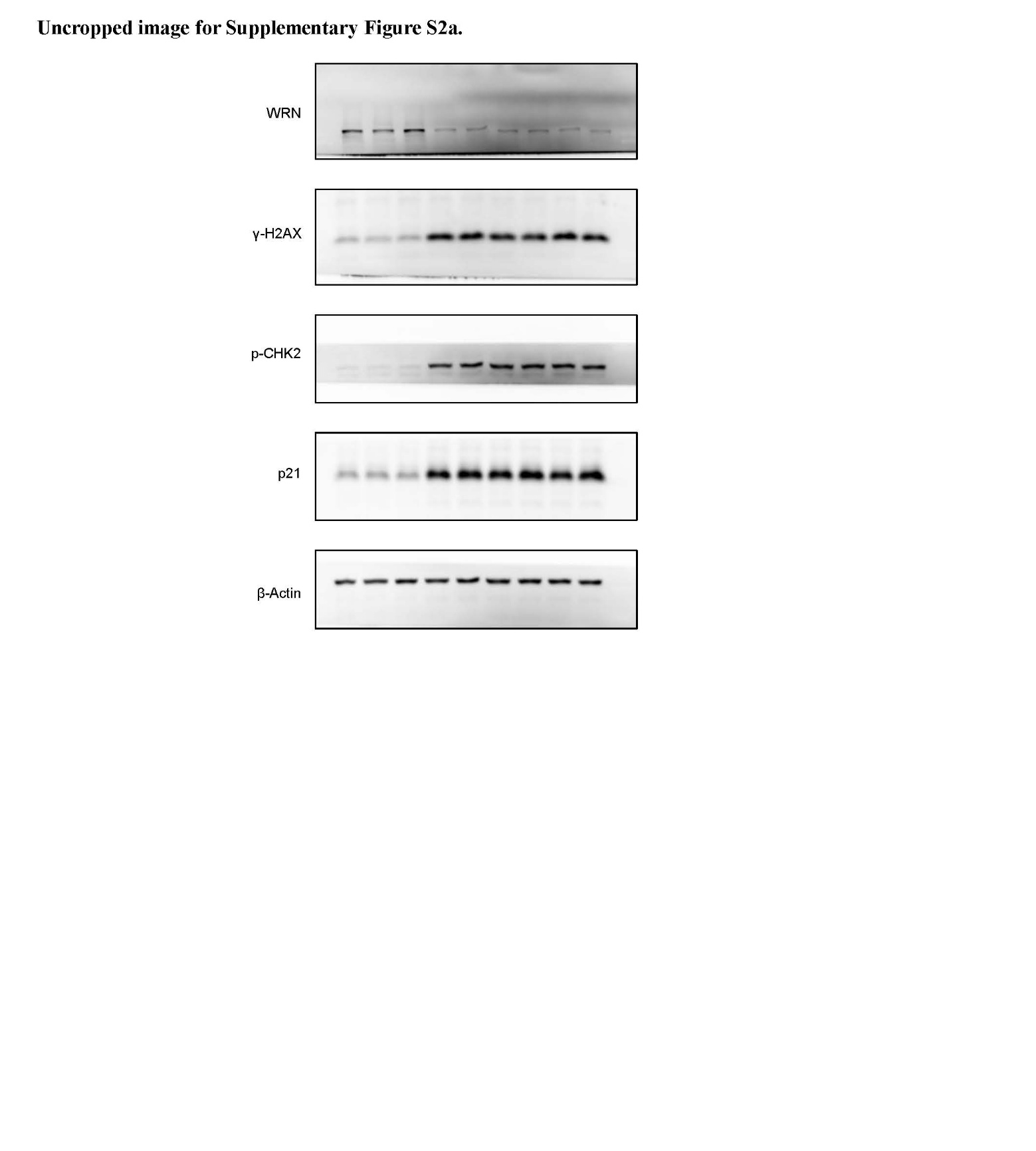

**Uncropped image corresponding to Supplementary Figure S2b.**

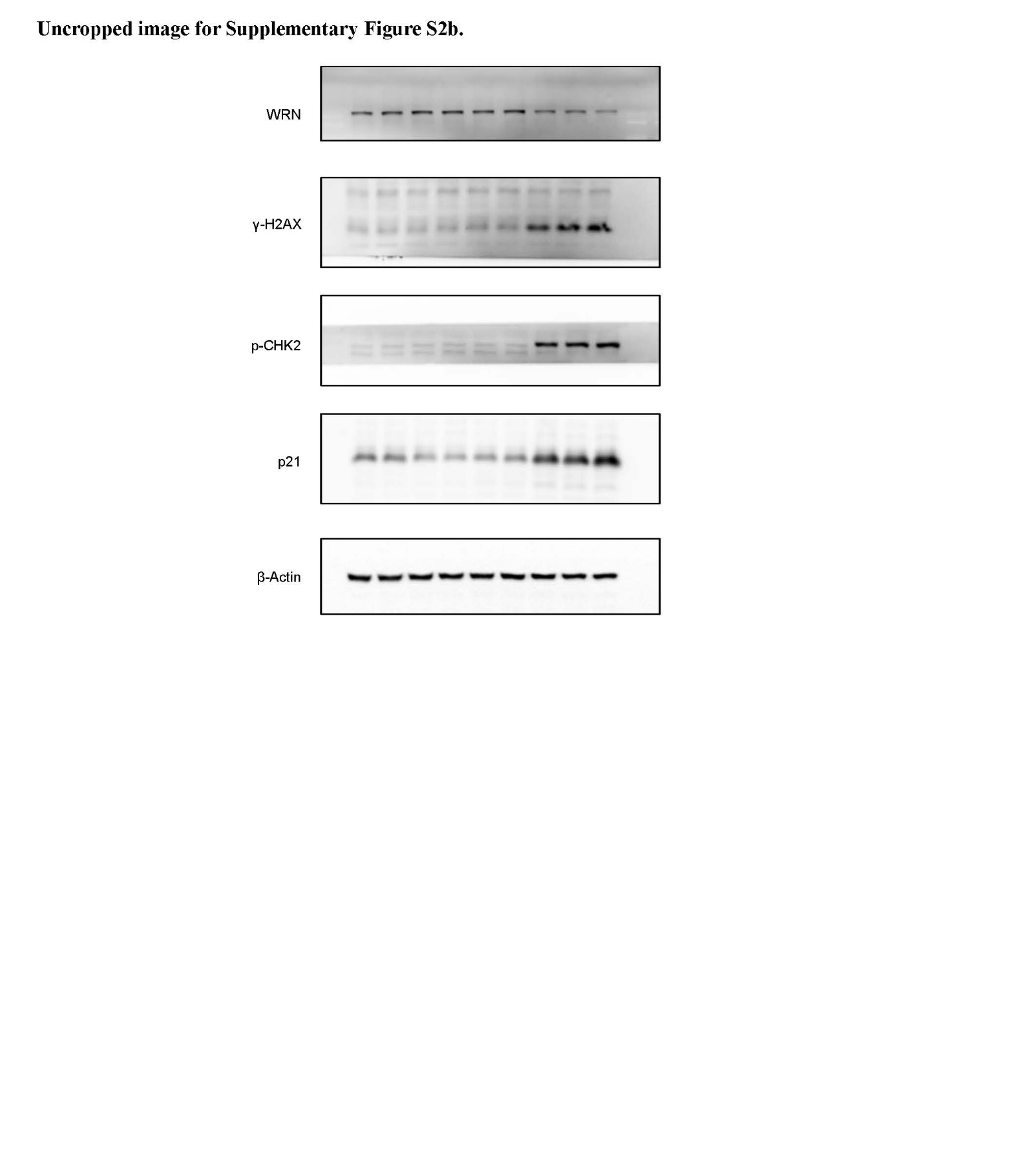

**Uncropped image corresponding to Supplementary Figure S2c.**

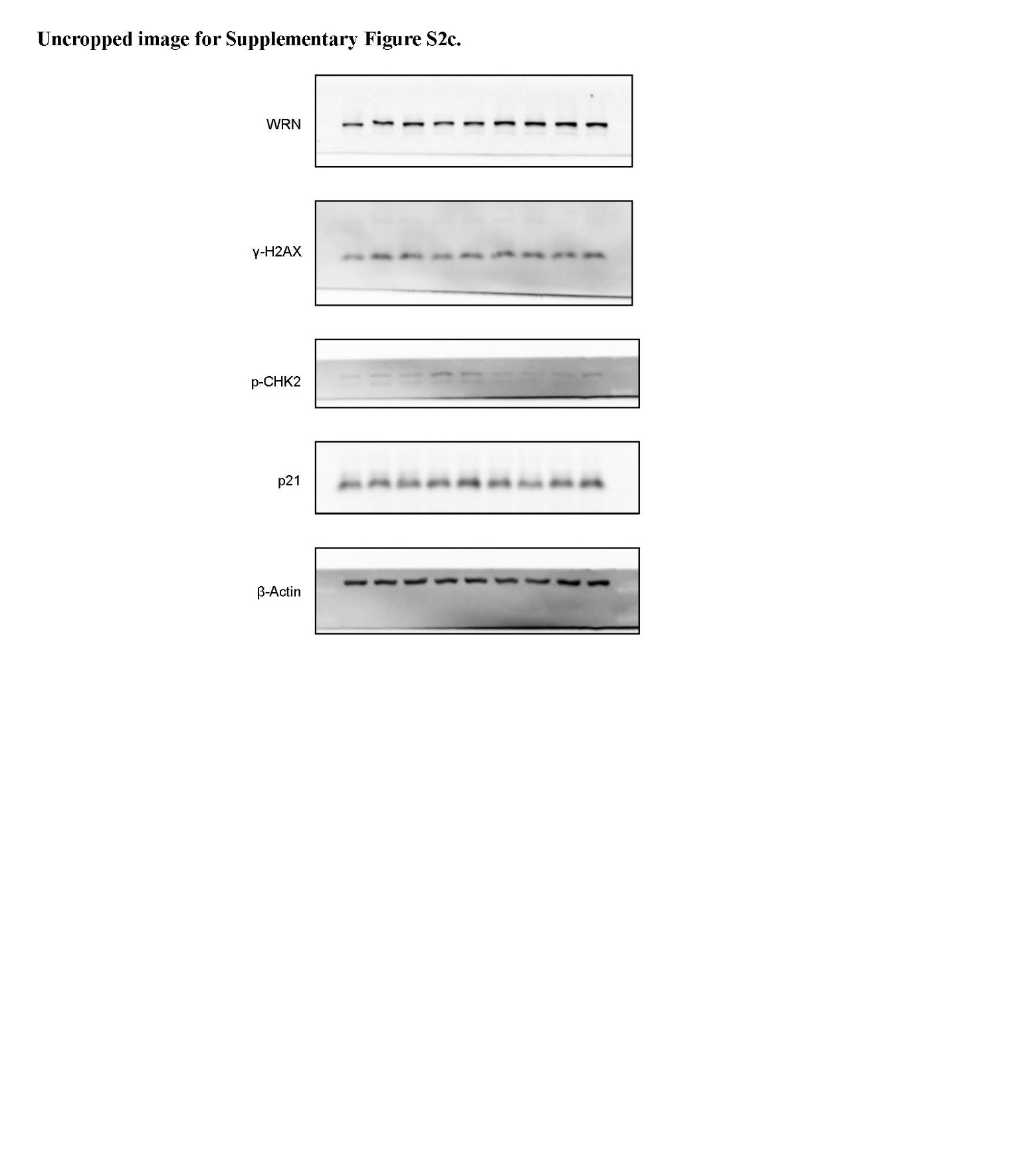

**Uncropped image corresponding to Supplementary Figure S2d.**

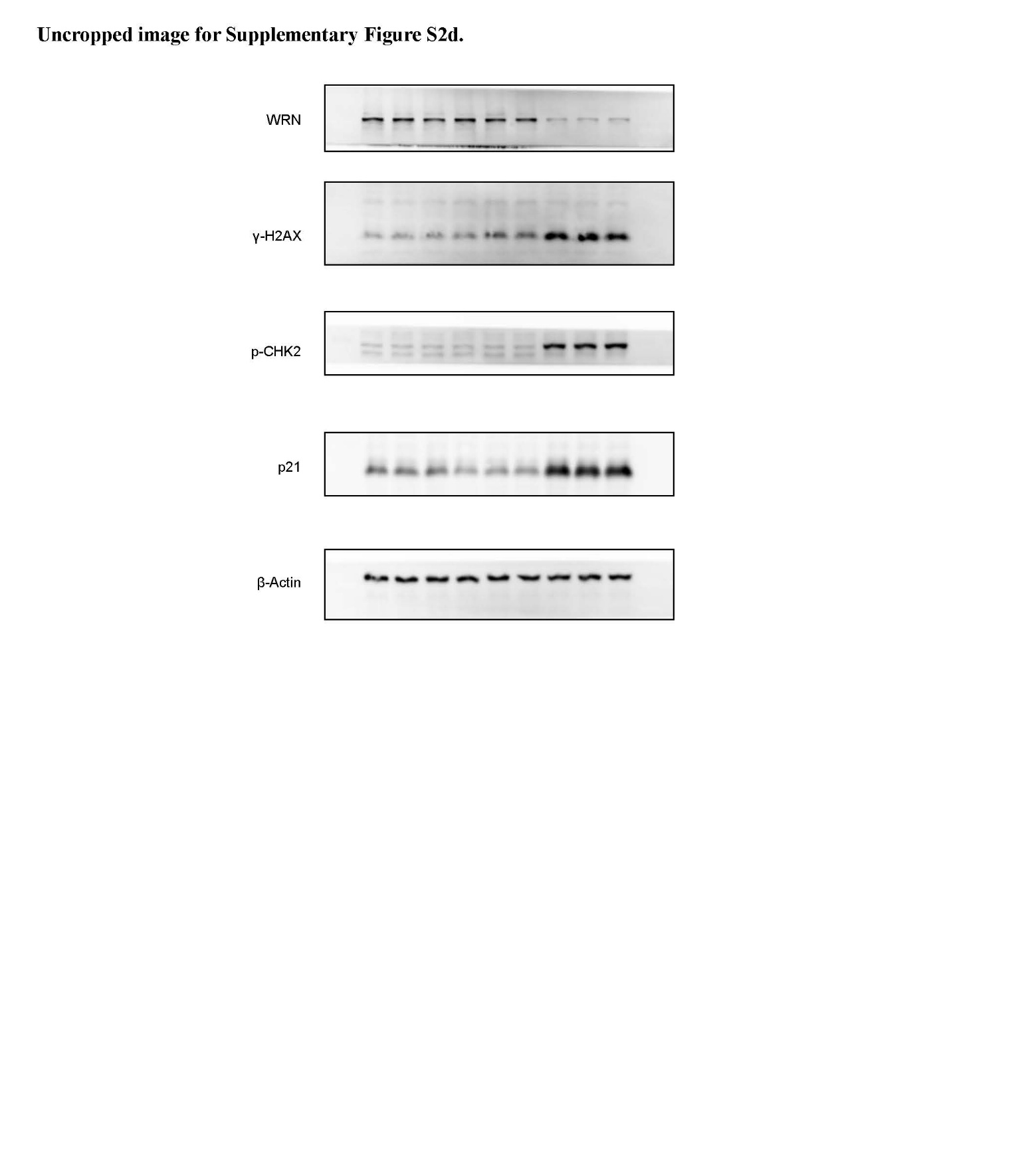

**Uncropped image corresponding to Supplementary Figure S2e.**

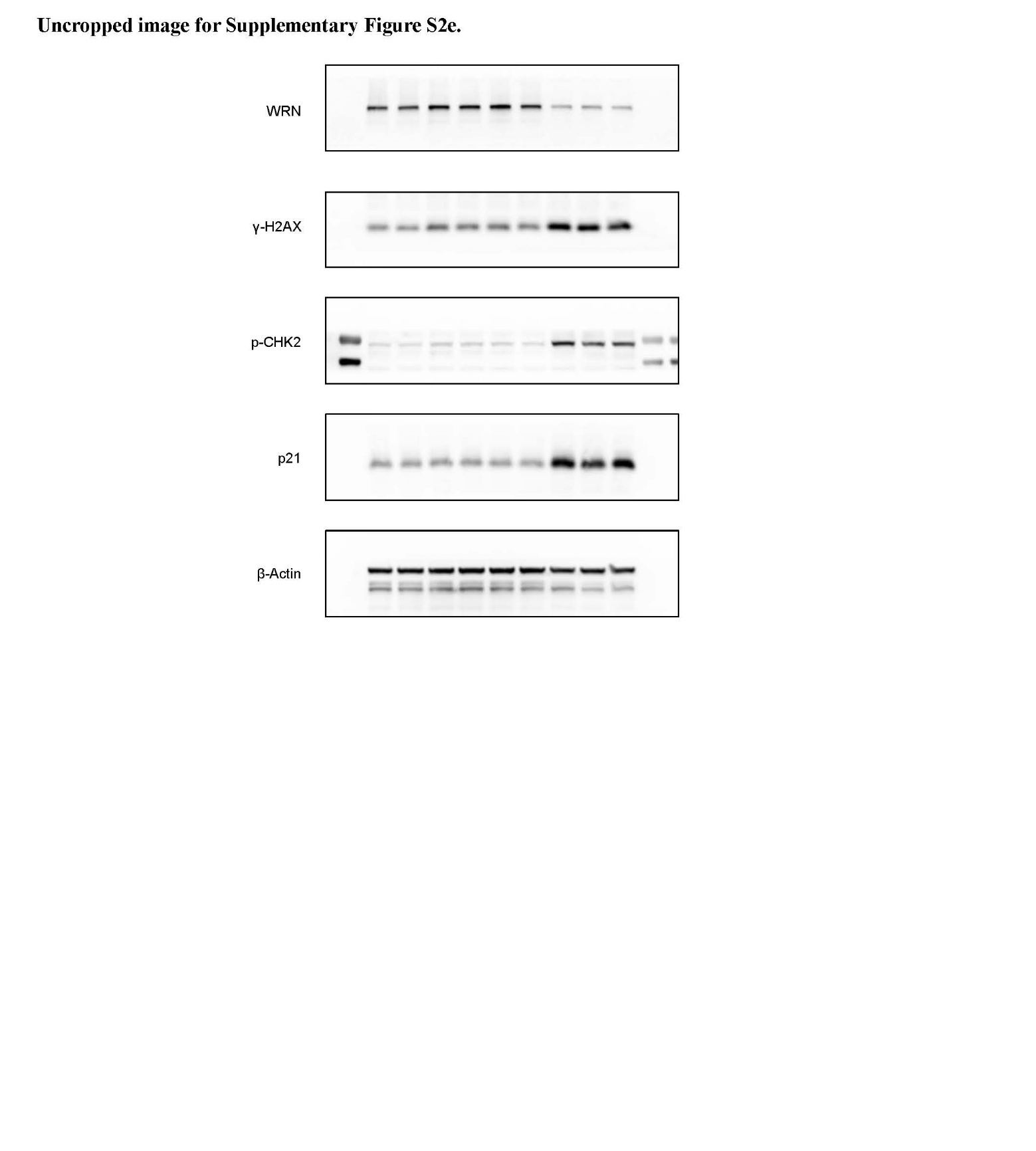

**Uncropped image corresponding to** **Supplementary Figure S2f.**

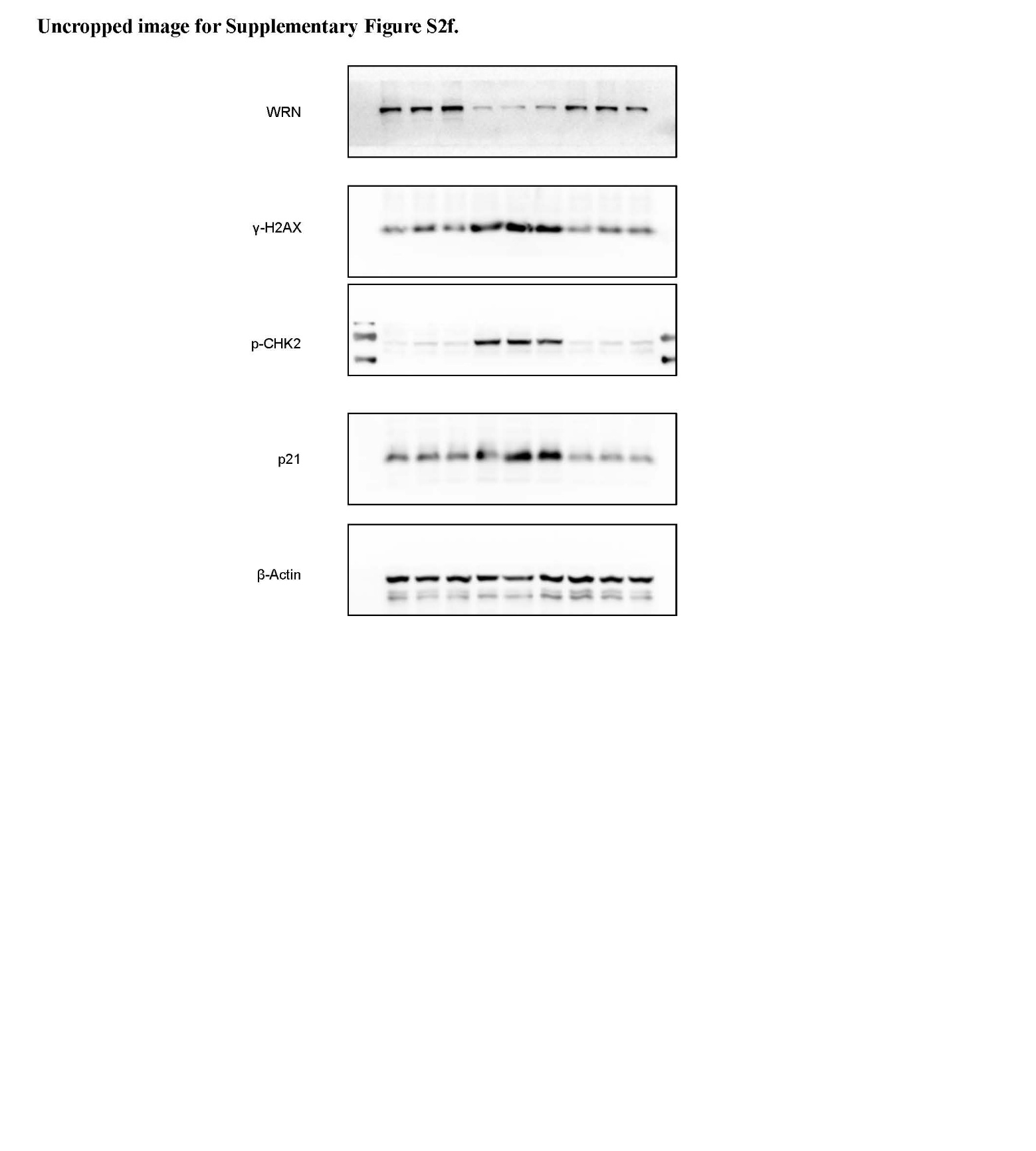

**Uncropped image corresponding to Supplementary Figure S2g.**

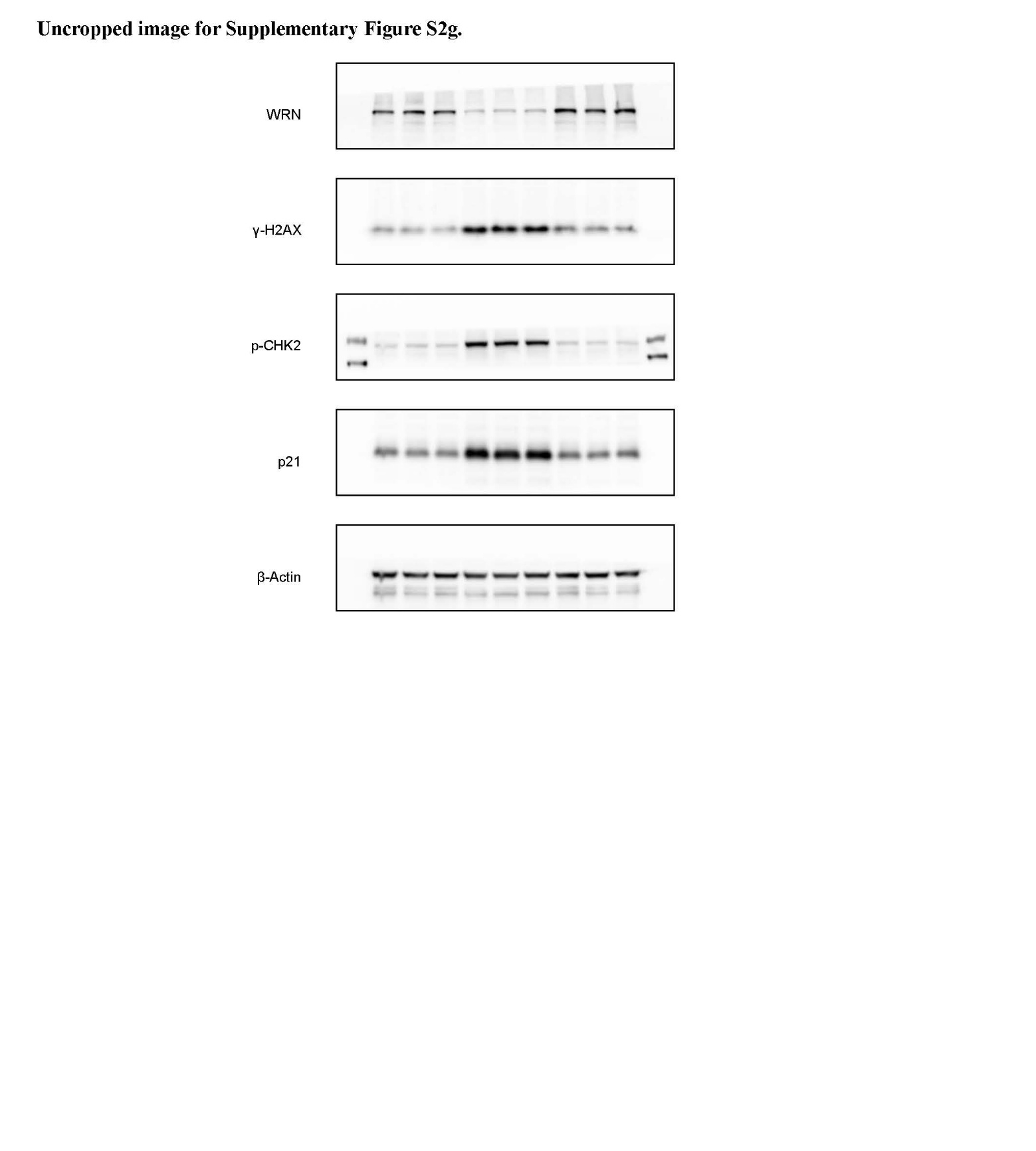
